# Interleukin-1 prevents SARS-CoV-2-induced membrane fusion to restrict viral transmission via induction of actin bundles

**DOI:** 10.1101/2024.05.16.594569

**Authors:** Xu Zheng, Shi Yu, Yanqiu Zhou, Kuai Yu, Yuhui Gao, Mengdan Chen, Dong Duan, Yunyi Li, Xiaoxian Cui, Jiabin Mou, Yuying Yang, Xun Wang, Min Chen, Yaming Jiu, Jincun Zhao, Guangxun Meng

## Abstract

Innate immune responses triggered by severe acute respiratory syndrome coronavirus 2 (SARS-CoV-2) infection play pivotal roles in the pathogenesis of COVID-19, while host factors including pro-inflammatory cytokines are critical for viral containment. By utilizing quantitative and qualitative models, we discovered that soluble factors secreted by human monocytes potently inhibit SARS-CoV-2-induced cell-cell fusion in viral-infected cells. Through cytokine screening, we identified that interleukin-1β (IL-1β), a key mediator of inflammation, inhibits syncytia formation mediated by various SARS-CoV-2 strains. Mechanistically, IL-1β activates RhoA/ROCK signaling through a non-canonical IL-1 receptor-dependent pathway, which drives the enrichment of actin bundles at the cell-cell junctions, thus prevents syncytia formation. Notably, *in vivo* infection experiments in mice confirms that IL-1β significantly restricted SARS-CoV-2 spreading in the lung epithelia. Together, by revealing the function and underlying mechanism of IL-1β on SARS-CoV-2-induced cell-cell fusion, our study highlights an unprecedented antiviral function for cytokines during viral infection.

## Introduction

The COVID-19 pandemic caused by severe acute respiratory syndrome coronavirus 2 (SARS-CoV-2) infection has spread globally, with at least 755 million people diagnosed and the death toll is over 6.8 million. SARS-CoV-2 variants of concern, including Alpha, Beta, Delta, and Omicron, continue to evolve and increase transmissibility and the ability to escape host immune responses. These variants have brought huge challenges to the design and development of vaccines and therapeutic reagents (Zhou et al., 2020). In order to discover novel strategies to control the virus, it is important to understand host responses to SARS-CoV-2 infection.

SARS-CoV-2 infection induces cell-cell fusion (also known as syncytia formation) in multiple cell types including lung epithelial cells, neurons and glia (Martínez-Mármol et al., 2023). Syncytia of SARS-CoV-2 infected cells with neighboring cells could potentially contribute to increased viral transmission and pathogenicity in the infected host (Rajah, Bernier, Buchrieser, & Schwartz, 2022), which also makes the virus insensitive to extracellular neutralizing antibodies (Li et al., 2022; Yu et al., 2023). Moreover, syncytia formation among pneumocytes with long-term persistence of viral RNA has been observed in the lung autopsy of deceased COVID-19 donors, which may contribute to prolonged clearance of the virus and long COVID symptoms (Bussani et al., 2020). Therefore, inhibiting syncytia formation is critical to ensure viral clearance and to control viral transmission.

It has been reported that SARS-CoV-2 mediated syncytia formation is effectively inhibited by multiple interferon (IFN)-stimulating genes (ISGs) (Pfaender et al., 2020; Wang et al., 2020). However, low IFN levels with impaired ISG responses were observed during early SARS-CoV-2 infection, which may have compromised the antiviral responses of IFN in severe COVID-19 patients (Blanco-Melo et al., 2020; Hadjadj et al., 2020). Thus, identifying other endogenous host factors that regulate syncytia formation is of great significance for harnessing the transmission of SARS-CoV-2.

Of note, a variety of cells are involved in the host responses to SARS-CoV-2 infection (Ren et al., 2021). Lung epithelial cells are the primary target of SARS-CoV-2 infection and transmission, which subsequently recruit and activate innate immune cells, leading to COVID-19 pathology (Barnett et al., 2023). In such process, tissue-resident macrophages and circulating monocytes contribute to local and systemic inflammation largely through releasing inflammatory cytokines (Sefik et al., 2022). Among these cytokines induced by SARS-CoV-2, the combination of TNF-α and IFN-γ induces inflammatory cell death, resulting in clear tissue damage, while other cytokines’ function remains obscure (Karki et al., 2021). In addition, when corticosteroids were applied to suppress the inflammatory response in patients infected by SARS-CoV (N. Lee et al., 2004) or MERS-CoV (Arabi et al., 2018), the clearance of viral RNA was obviously delayed, suggesting the importance of innate immune factors in viral clearance.

Innate immune cells express toll-like receptors (TLRs), and TLR-mediated signaling induces robust production of inflammatory cytokines (Medzhitov, 2001). In the current work, we screened the role of soluble pro-inflammatory cytokines on the SARS-CoV-2 spike-induced cell-cell fusion. Notably, we identified that IL-1β, which is the key factor of inflammatory response, inhibited SARS-CoV-2 spike-induced syncytia formation in various cells by activating RhoA/ROCK pathway to initiate actin bundle formation at cell-cell interface between SARS-CoV-2 infected cells and neighboring cells. Importantly, IL-1β significantly reduced SARS-CoV-2 transmission among lung epithelia in experimental mice *in vivo*. Therefore, our data highlight an important role for pro-inflammatory cytokines against viral infection.

## Results

### Host factors secreted by activated innate immune cells inhibit SARS-CoV-2-induced cell-cell fusion

We have previously established quantitative and qualitative models for SARS-CoV-2 spike-induced cell-cell fusion by bioluminescence assay, immunoblotting and fluorescence imaging (Yu et al., 2022). In order to explore the potential effect of cytokines on SARS-CoV-2-induced cell-cell fusion, human monocyte cell line THP-1 and human peripheral blood mononuclear cells (PBMCs) were used in this study. We applied several TLR ligands to stimulate such innate immune cells and collected the cell culture supernatants for subsequent experiments **(Figure 1A)**. Of note, cell culture supernatants of THP-1 cells stimulated by TLR ligands significantly reduced the bioluminescence signal, while neither untreated THP-1 cell culture supernatant nor the medium control had any effect on the bioluminescence signal reflecting cell-cell fusion **(Figure 1B)**. SARS-CoV-2 spike engagement of ACE2 primed the cleavage of S2’ fragment in target cells, a key proteolytic event coupled with spike-mediated membrane fusion (Yu et al., 2022). In parallel with bioluminescence assay, a large amount of enriched S2’ cleavage was detected in HEK293T-Spike and HEK293T-ACE2 co-cultured group and co-culture incubated with untreated THP-1 cell culture supernatant, while S2’ cleavage was clearly reduced upon treatment with TLR ligands-stimulated THP-1 cell culture supernatants **(Figure 1C)**. Syncytia formation was also visualized using cells co-expressing spike and a ZsGreen fluorescent reporter. The area of syncytium were significantly reduced by the treatment with TLR ligands-stimulated THP-1 cell culture supernatants **(Figure 1D and Figure 1—figure supplement 1A)**. Considering the presence of TLR ligands in such cell culture supernatants, we tested their potential direct effects. As expected, TLR ligands alone did not reduce the bioluminescence signal and S2’ cleavage compared to the control groups, as well as no effect on syncytia formation **(Figure 1—figure supplement 1B-E)**.

**Figure 1.**
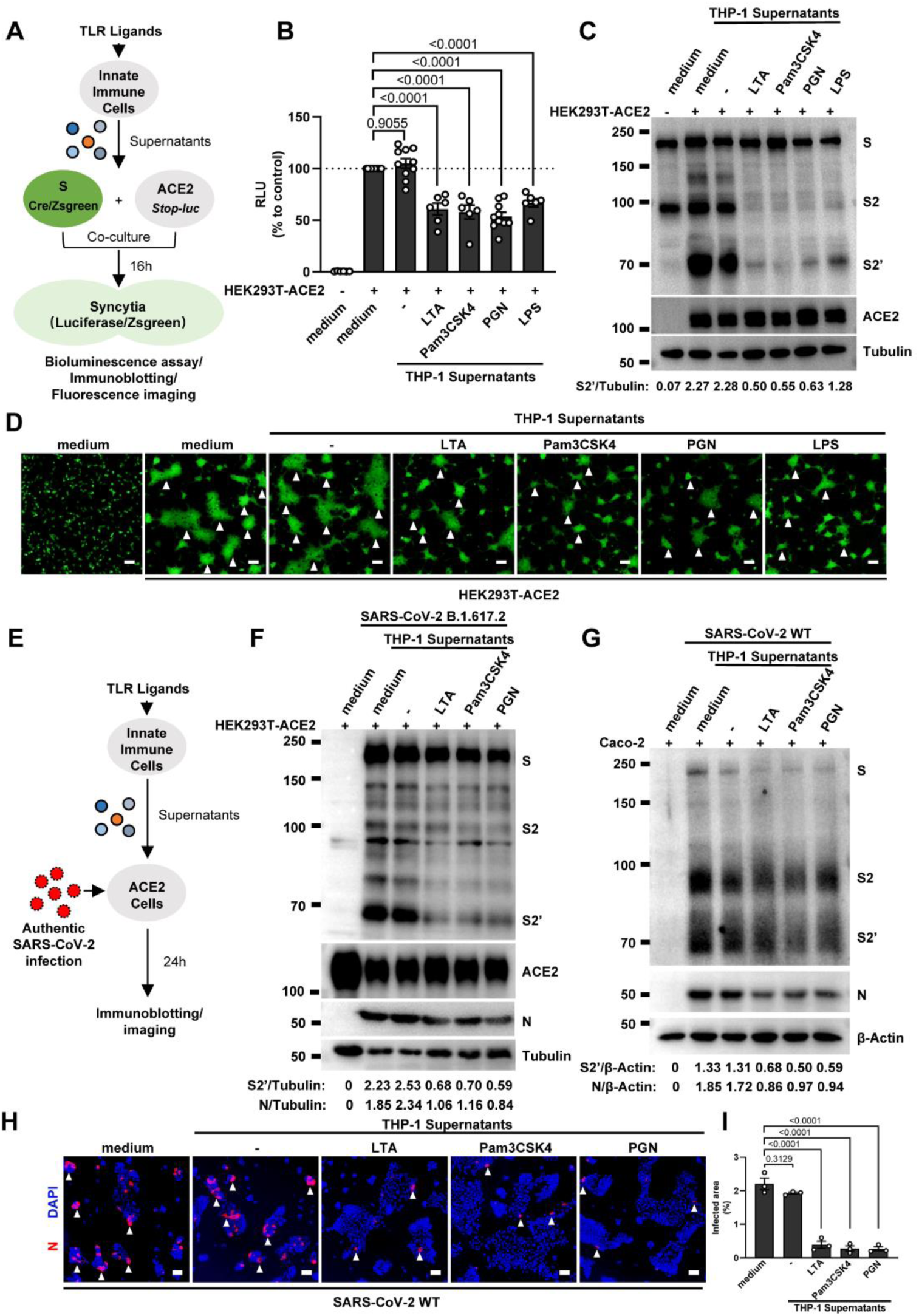
Host factors secreted by activated innate immune cells inhibit SARS-CoV-2-induced cell-cell fusion. (**A**) Schematics of the cell-cell fusion model used to quantify spike-mediated syncytium formation upon treatment with cell culture supernatants from TLR ligands-stimulated innate immune cells. Cells co-expressing SARS-CoV-2 spike and Cre, were co-cultured with ACE2 and *Stop-luc* co-expressing HEK293T cells for 16 hours, before cell lysates were collected for bioluminescence assay and immunoblotting. Cells co-expressing SARS-CoV-2 spike and Zsgreen, were co-cultured with ACE2 expressing HEK293T cells for 16 hours before fluorescence imaging. (**B**) Luciferase activity (RLU) measured from HEK293T cell lysates collected from THP-1 supernatants-treated HEK293T-S and HEK293T-ACE2 described in (A) for 16 hours. FBS free RPMI 1640 served as medium control. Data are representative of six individual repeats and displayed as individual points with mean ± standard error of the mean (SEM). (**C**) Immunoblots showing full-length spike, S2, cleaved S2’ and ACE2 collected from THP-1 supernatants-treated HEK293T-S and HEK293T-ACE2 described in (A) for 16 hours. Blots are representative of three independent experiments. Numbers below the blots indicated the intensity of S2’ versus Tubulin. (**D**) Representative fluorescent image captured at 488 nm from THP-1 supernatants-treated HEK293T-S-Zsgreen and HEK293T-ACE2 for 16 hours. (**E**) Schematic presentation of THP-1 supernatants pre-treatment on authentic SARS-CoV-2 infected cells. Pre-treatment of HEK293T-ACE2 cells with THP-1 supernatants for 1 hour, then inoculated with 0.5 multiplicity of infection (MOI) Delta or WT authentic SARS-CoV-2 virus. Imaging was performed at 24 hours post-infection (hpi) before cell lysates were harvested for immunoblotting. (**F**) Immunoblots of Delta SARS-CoV-2 S, S2, cleaved S2’, N and ACE2 proteins collected from HEK293T-ACE2 cells 24 hpi as described in (E). Blots are representative of three individual experiments. Numbers below the blots indicated the intensity of S2’ or N versus Tubulin. (**G**) Immunoblots of WT SARS-CoV-2 S, S2, cleaved S2’ and N proteins collected from Caco-2 cells 24 hpi as described in (E). Blots are representative of three individual experiments. Numbers below the blots indicated the intensity of S2’ or N versus β-Actin. (**H**) Immunofluorescent images showing morphology of SARS-CoV-2 infected Caco-2 cells pre-treated with THP-1 supernatants. Anti-SARS-CoV-2 N was stained with Alexa fluor 555, and nuclei were counterstained with DAPI, respectively. White arrow heads (D and H) indicate syncytia formation or infected cells, scale bars are indicative of 50 μm and images are representative of three independent experiments. (**I**) Quantification of the infected area in (H).

Concurrently, we also tested the effect of PBMCs culture supernatants on SARS-CoV-2 spike-induced cell-cell fusion. Consistent with the results from THP-1 cells, TLR ligands stimulated PBMCs culture supernatants treatment also strongly reduced the bioluminescence signal, S2’ cleavage, and the area of syncytium compared with the medium group **(Figure 1—figure supplement 1F-I)**. These results thus suggested that activated innate immune cells released host factors to inhibit SARS-CoV-2 spike-induced cell-cell fusion.

To validate the effect of innate immune cell culture supernatants on cell-cell fusion in authentic SARS-CoV-2 infection, we pre-treated ACE2-expressing cells with THP-1 cell culture supernatants before inoculation with SARS-CoV-2 B.1.617.2 (Delta) or wild type (WT) strains. Cell lysates were used for the detection of SARS-CoV-2 spike and N protein 24 hours post infection (hpi) **(Figure 1E)**. Result of this experiment revealed that TLR ligands-stimulated-THP-1 cell culture supernatants reduced S2’ cleavage and N protein during Delta or WT SARS-CoV-2 infection in HEK293T-ACE2 cells, while untreated THP-1 cell culture supernatant had no effect **(Figure 1F and Figure 1—figure supplement 2A)**. In addition, TLR ligands-stimulated-THP-1 cell culture supernatants reduced the area of syncytium induced by Delta or WT SARS-CoV-2 infection **(Figure 1—figure supplement 2B and C)**. Furthermore, we infected the human colon epithelial carcinoma cell line Caco-2 with WT SARS-CoV-2, and found that S2’ cleavage and N protein were reduced after TLR ligands-stimulated-THP-1 cell culture supernatants pre-treatment **(Figure 1G)**. Accordingly, immunofluorescent staining also showed that TLR ligands-stimulated-THP-1 cell culture supernatants significantly reduced the area of syncytium during SARS-CoV-2 infection in Caco-2 cells **(Figure 1H and I)**. Therefore, these data suggested that host factors secreted by activated innate immune cells inhibit authentic SARS-CoV-2 induced cell-cell fusion.

### IL-1β inhibits SARS-CoV-2-induced cell-cell fusion

To explore which host factor(s) inhibited SARS-CoV-2 induced cell-cell fusion, we first detected mRNA levels of different cytokines in THP-1 cells stimulated by TLR ligands. It was found that the expression levels of *IL1A*, *IL1B*, *IL6* and *IL8* were significantly increased upon TLR ligands stimulation, while *IL4*, *IL12A*, *IFNA1*, *IFNB1* and *IFNG* mRNA levels were not changed or undetected **(Figure 2—figure supplement 1A)**. In addition, we also detected the mRNA levels of cytokine receptors in HEK293T modelling cells, confirming that *IL1R1*, *IL4R*, *IL6ST*, *IL8RA*, *IFNAR1*, *IFNGR1* were expressed in such cells, while *IL2RA* and *IL12RB1* were undetectable **(Figure 2—figure supplement 1B)**.

We next selected recombinant IL-1α, IL-1β, IL-6, and IL-8 to test whether individual cytokine may play a role in affecting SARS-CoV-2 spike induced cell-cell fusion **(Figure 2A)**. Interestingly, IL-1α and IL-1β significantly reduced the bioluminescence signal compared to the control group, while IL-6 and IL-8 had little or no effect **(Figure 2B)**. In addition, fluorescence images of cells expressing Zsgreen reporter also confirmed that IL-1α and IL-1β significantly inhibited SARS-CoV-2 spike induced syncytia formation **(Figure 2—figure supplement 1C and D)**. Furthermore, IL-1β and IL-1α both reduced the bioluminescence signal and S2’ cleavage **(Figure 2 C and D, and Figure 2—figure supplement 2A)** in cell lysates in a dose-dependent manner. Moreover, the syncytia formation was inhibited with increasing concentrations of IL-1β or IL-1α **(Figure 2— figure supplement 2B-E)**. Intriguingly, when we added both IL-1α and IL-1β, there was no synergistic inhibition on cell-cell fusion compared to either cytokine alone **(Figure 2—figure supplement 2F-H)**, suggesting a saturation of IL-1 receptor binding to these homologues. Since both IL-1α and IL-1β activate the downstream pathway through the same receptor IL-1R1, these data suggested that IL-1α or IL-1β may inhibit cell-cell fusion through the same pathway. Considering the higher mRNA level of *IL1B* than *IL1A,* as well as the classical release pathway of IL-1β from innate immune cells (Weber, Wasiliew, & Kracht, 2010), we applied IL-1β for further experiments.

**Figure 2.**
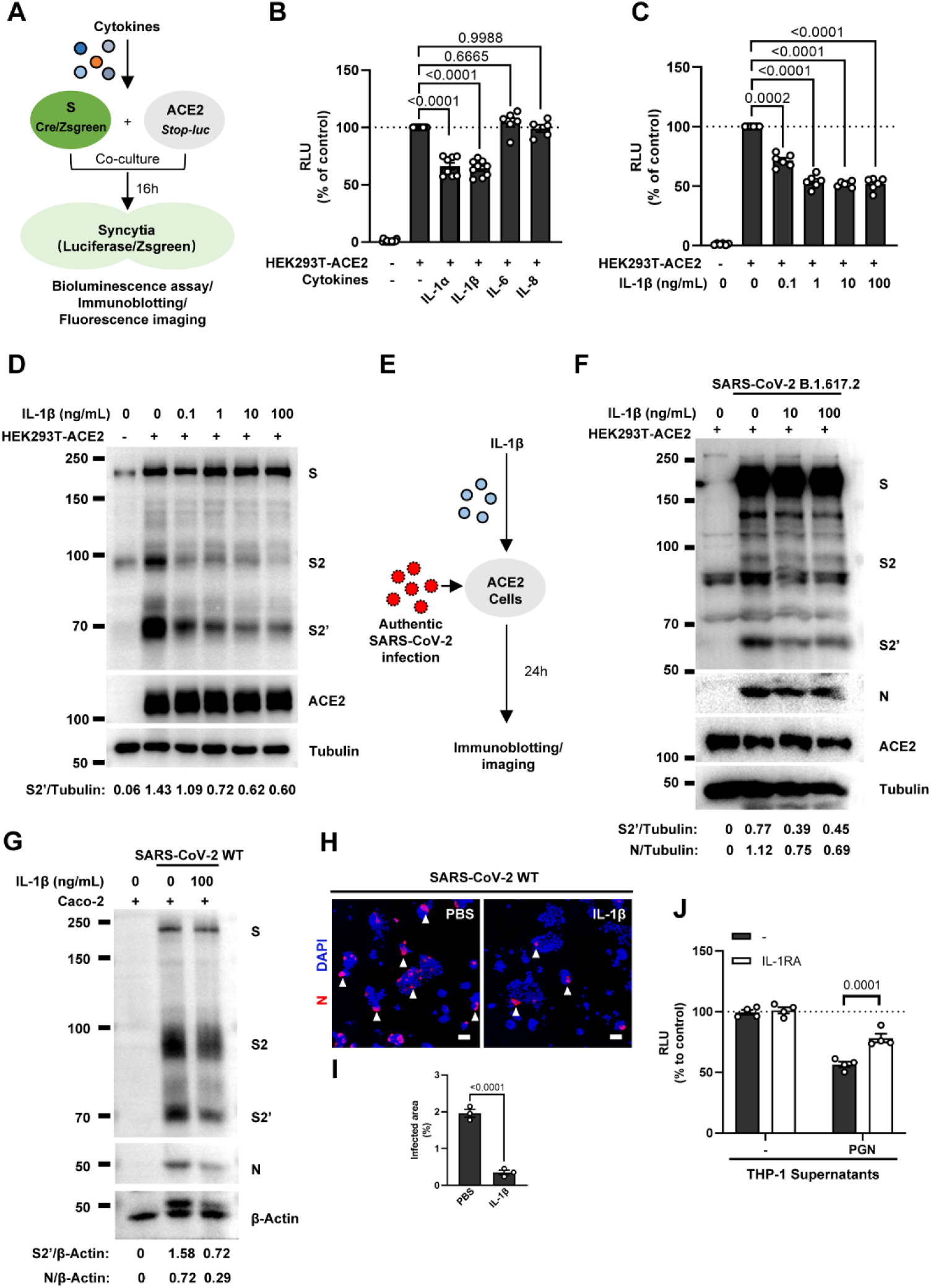
IL-1β inhibits SARS-CoV-2-induced cell-cell fusion. (**A**) Schematics of the cell-cell fusion model used to quantify spike-mediated syncytium formation upon treatment with different cytokines. Cells co-expressing SARS-CoV-2 spike and Cre, were co-cultured with ACE2 and *Stop-luc* co-expressing HEK293T cells for 16 hours, before cell lysates were collected for bioluminescence assay and immunoblotting. Cells co-expressing SARS-CoV-2 spike and Zsgreen, were co-cultured with ACE2 expressing HEK293T cells for 16 hours before fluorescence imaging. (**B**) Luciferase activity (RLU) measured from HEK293T cell lysates collected from different cytokines-treated HEK293T-S and HEK293T-ACE2 described in (A) for 16 hours. IL-1α (10 ng/mL), IL-1β (1 ng/mL), IL-6 (100 ng/mL) or IL-8 (100 ng/mL) were added into the cell-cell fusion system. Data are representative of six individual repeats and displayed as individual points with mean ±SEM. (**C**) Luciferase activity (RLU) measured from HEK293T cell lysates collected from different concentrations of IL-1β-treated HEK293T-S and HEK293T-ACE2 for 16 hours. Data are representative of six individual repeats and displayed as individual points with mean ±SEM. (**D**) Immunoblots showing full-length spike, S2, cleaved S2’ and ACE2 collected from different concentrations of IL-1β-treated HEK293T-S and HEK293T-ACE2 for 16 hours. Blots are representative of three independent experiments. Numbers below the blots indicated the intensity of S2’ versus Tubulin. (**E**) Schematic presentation of IL-1β pre-treatment on authentic SARS-CoV-2 infected cells. Pre-treatment of HEK293T-ACE2 cells with different concentrations of IL-1β for 1 hour, then inoculated with 0.5 MOI Delta or WT authentic SARS-CoV-2 virus. Brightfield images were captured at 24 hours post-infection (hpi) before cell lysates were harvested for immunoblotting. (**F**) Immunoblots of Delta SARS-CoV-2 S, S2, cleaved S2’, N and ACE2 proteins collected from HEK293T-ACE2 cells 24 hpi as described in (E). Blots are representative of three individual experiments. Numbers below the blots indicated the intensity of S2’ or N versus Tubulin. (**G**) Immunoblots of WT SARS-CoV-2 S, S2, cleaved S2’ and N proteins collected from Caco-2 cells 24 hpi as described in (E). Blots are representative of three individual experiments. Numbers below the blots indicated the intensity of S2’ or N versus β-Actin. (**H**) Immunofluorescent images showing morphology of SARS-CoV-2-infected Caco-2 cells pre-treated with or without IL-1β. Anti-SARS-CoV-2 N was stained with Alexa fluor 555, and nuclei were counterstained with DAPI, respectively. White arrow heads indicate syncytia formation or infected cells, scale bars are indicative of 50 μm and images are representative of three independent experiments. (**I**) Quantification of the infected area in (H). (**J**) Luciferase activity (RLU) measured from THP-1 supernatants-treated HEK293T-S and HEK293T-ACE2 in the presence or absence of IL-1RA. Data are representative of four individual repeats and displayed as individual points with mean ±SEM.

In order to validate the effect of IL-1β on cell-cell fusion during authentic SARS-CoV-2 infection, we pre-treated ACE2-expressing cells with IL-1β before inoculating Delta or WT authentic SARS-CoV-2. Cell lysates were used for the detection of SARS-CoV-2 spike and N protein 24 hpi **(Figure 2E)**. To this end, it was found that IL-1β reduced S2’ cleavage and N protein compared to the control group during such infection both in HEK293T-ACE2 **(Figure 2F and Figure 2—figure supplement 3A)** and Caco-2 cells **(Figure 2G)**. Meanwhile, IL-1β inhibited authentic SARS-CoV-2 induced syncytia formation **(Figure 2H and I, and Figure 2—figure supplement 3B and C)**. Thus, these results verified that IL-1β inhibits authentic SARS-CoV-2 induced cell-cell fusion in various target cells.

As expected, innate immune cells activated by TLR ligands secreted IL-1β into the cell culture supernatants **(Figure 2—figure supplement 4A and B).** We employed IL-1 receptor antagonist (IL-1RA) to block IL-1 receptor on target cells, and found that IL-1RA treatment reduced the inhibitory effect of PGN-stimulated-THP-1 cell culture supernatant on cell-cell fusion **(Figure 2J and Figure 2—figure supplement 4C)**. With another note, TLR2 was essential for THP-1 cells to release IL-1β in response to TLR2 ligands **(Figure 2—figure supplement 4D)**. More importantly, the cell culture supernatants of TLR2-knockout THP-1 cells stimulated by TLR2 Ligands had no effect on the bioluminescence signal, while the cell culture supernatants from WT THP-1 cells stimulated by the same TLR2 Ligands significantly reduced the bioluminescence signal **(Figure 2—figure supplement 4E)**. In addition, pre-treatment with TAK1 inhibitor (5Z-7) or IKKβ inhibitor (TPCA1) in WT THP-1 cells prevented IL-1β secretion after PGN stimulation **(Figure 2—figure supplement 4F)**, as well as eliminated the inhibitory effect of PGN-stimulated WT THP-1 cell culture supernatant on SARS-CoV-2 spike-induced cell-cell fusion **(Figure 2— figure supplement 4G)**. In parallel, pre-treatment with these inhibitors in PBMCs showed the same results **(Figure 2—figure supplement 4H and I)**. These data suggested that TLR-knockout or inhibitors targeting the respective TLR signaling prevented innate immune cells from releasing IL-1β into supernatants, which led to failed inhibition of SARS-CoV-2 spike induced cell-cell fusion. These findings thus further verify that IL-1β is an important host factor inhibiting SARS-CoV-2 induced cell-cell fusion.

To investigate the effector function of IL-1 on cells expressing SARS-CoV-2 spike (donor cells) and neighboring cells expressing ACE2 (acceptor cells), we pre-treated HEK293T-S or HEK293T-ACE2 cells or both with IL-1β, then co-cultured after washing with PBS; cells were then analyzed by the quantitative and qualitative models **(Figure 2—figure supplement 5A)**. Notably, pre-treatment of either HEK293T-S or HEK293T-ACE2 cells with IL-1β alone reduced bioluminescence signal and S2’ cleavage; when IL-1β pre-treatment on both HEK293T-S and HEK293T-ACE2 cells was applied, bioluminescence signal and S2’ cleavage were further reduced **(Figure 2—figure supplement 5B)**. Furthermore, we also applied Vero E6-overexpressing ACE2 cell line (Vero E6-ACE2) and human Calu-3 cells as acceptor cells; and found that pre-treatment of either HEK293T-S or Vero E6-ACE2 cells with IL-1β alone reduced part of S2’ cleavage, while IL-1β pre-treatment of both HEK293T-S and Vero E6-ACE2 cells led to further reduction of S2’ cleavage **(Figure 2—figure supplement 5C),** and the same results were observed in the case of Calu-3 as acceptor cells **(Figure 2—figure supplement 5D)**.

Accordingly, fluorescence imaging also showed that IL-1β significantly reduced the area of syncytia **(Figure 2—figure supplement 6A and B)**. Notably, IL-1β reduced the bioluminescence signal and S2’ cleavage in different SARS-CoV-2 variants **(Figure 2—figure supplement 6C-E)**. Therefore, these results suggest that IL-1β acts on both donor and acceptor cells to inhibit SARS-CoV-2 spike-induced cell-cell fusion in various cell lines.

Of note, SARS-CoV (Belouzard, Chu, & Whittaker, 2009) and MERS-CoV (Straus et al., 2020) spike proteins also induce cell-cell fusion in target cells. Therefore, we further explored whether IL-1β was also able to inhibit SARS-CoV and MERS-CoV spike-induced cell-cell fusion in ACE2- or Dipeptidyl peptidase-4 (DPP4)-expressing cells by bioluminescence assay, immunoblotting and a modified *stop-mCherry* fluorescent model, wherein mCherry reporter is only expressed when Cre excises the Stop cassette inside the fused syncytia **(Figure 2—figure supplement 7A)**. Similar to SARS-CoV-2 spike-induced cell-cell fusion, IL-1β also reduced bioluminescence signal **(Figure 2—figure supplement 7B and C)**, S2’ cleavage **(Figure 2—figure supplement 7D and E)** and the area of syncytium **(Figure 2—figure supplement 7F and G)** in these cell-cell fusion systems. Thus, IL-1β possesses a broad spectrum to inhibit cell-cell fusion induced by different coronaviruses.

### IL-1β inhibits SARS-CoV-2-induced cell-cell fusion through IL-1R1/MyD88/IRAK/TRAF6 pathway

To investigate the mechanism of IL-1β inhibition on SARS-CoV-2 induced cell-cell fusion, we performed gene knockout using CRISPR-Cas9 technology, in conjunction with inhibitors targeting the IL-1 receptor pathway **(Figure 3A)**. First of all, in the presence of IL-1RA, IL-1β was unable to reduce bioluminescence signal and S2’ cleavage **(Figure 3B)**. Next, as MyD88 is the downstream adaptor for IL-1R1, we generated MyD88 knockout HEK293T cell line, wherein IL-1β was unable to reduce bioluminescence signal **(Figure 3C)** and S2’ cleavage **(Figure 3—figure supplement 1A)**. In addition, we found that IL-1β was unable to reduce bioluminescence signal and S2’ cleavage in the presence of IRAK1/4 inhibitor **(Figure 3D)**. Furthermore, IL-1β was unable to reduce bioluminescence signal **(Figure 3E)** and S2’ cleavage **(Figure 3—figure supplement 1B)** in TRAF6 knockout HEK293T cell line. These results suggested that IL-1β inhibits SARS-CoV-2 spike induced cell-cell fusion through IL-1R1-MyD88-IRAK-TRAF6 pathway.

**Figure 3.**
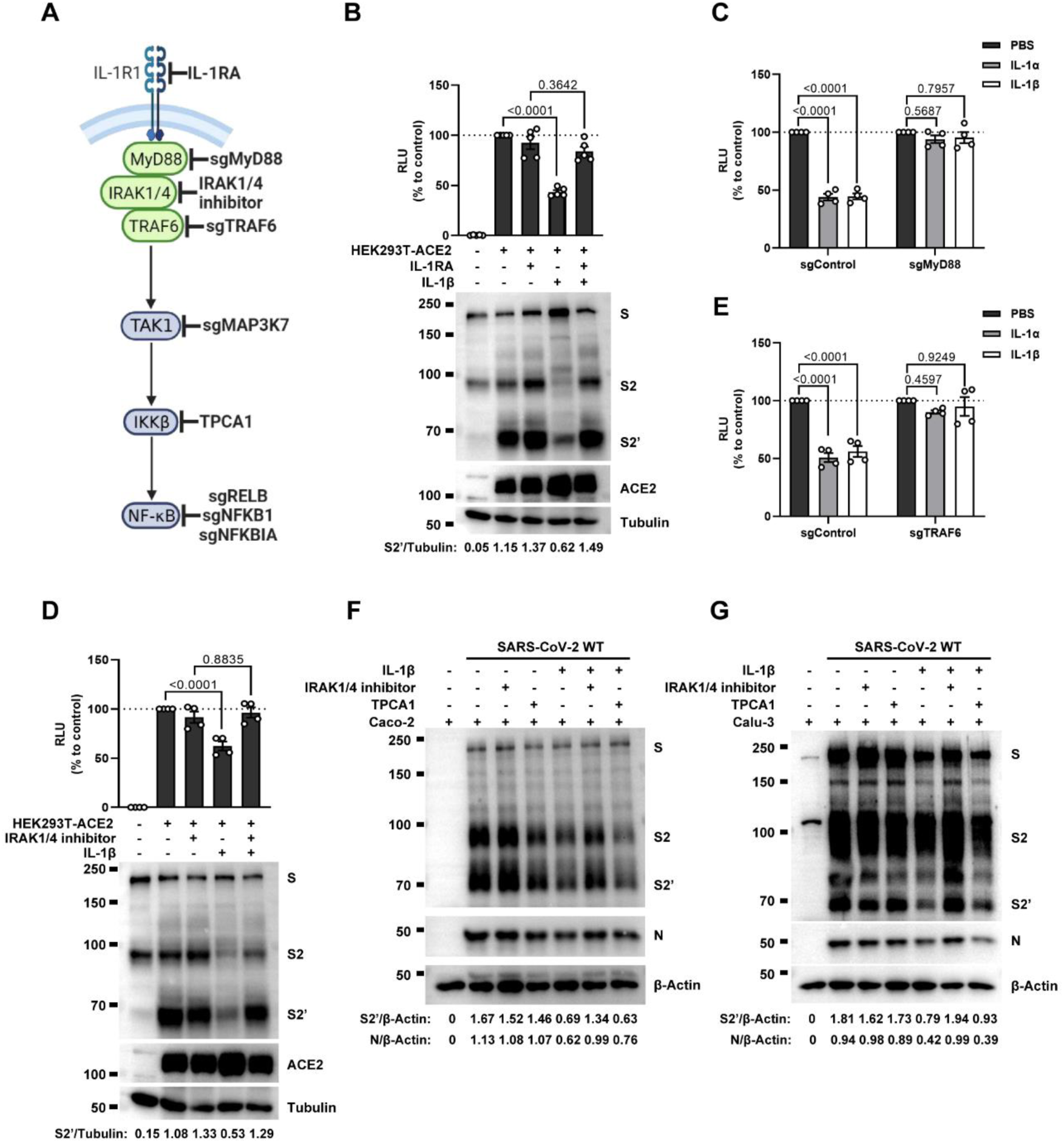
IL-1β inhibits SARS-CoV-2-induced cell-cell fusion through the IL-1R1/MyD88/IRAK/TRAF6 pathway. (**A**) Schematics of gene knockout or inhibitor treatment in the IL-1 receptor pathway. (**B**) Luciferase activity (RLU) measured from HEK293T cell lysates and immunoblots showing full-length spike, S2, cleaved S2’ and ACE2 collected from HEK293T-S and HEK293T-ACE2 pre-treated with 1000 ng/mL IL-1RA for 30 min, then treated with 1 ng/mL IL-1β for 16 hours. Data and blots are representative of five individual repeats. Numbers below the blots indicated the intensity of S2’ versus Tubulin. (**C**) Luciferase activity (RLU) measured from cell lysates collected from 10 ng/mL IL-1α or 1 ng/mL IL-1β-treated sgControl or sgMyD88 HEK293T cell-cell fusion system for 16 hours. Data are representative of four individual repeats and displayed as individual points with mean ±SEM. (**D**) Luciferase activity (RLU) measured from HEK293T cell lysates and immunoblots showing full-length spike, S2, cleaved S2’ and ACE2 collected from HEK293T-S and HEK293T-ACE2 pre-treated with 2μM IRAK1/4 inhibitor for 30 min, then treated with 1 ng/mL IL-1β for 16 hours. Data and blots are representative of four individual repeats. Numbers below the blots indicated the intensity of S2’ versus Tubulin. (**E**) Luciferase activity (RLU) measured from cell lysates collected from 10 ng/mL IL-1α or 1 ng/mL IL-1β-treated sgControl or sgTRAF6 HEK293T cell-cell fusion system for 16 hours. Data are representative of four individual repeats and displayed as individual points with mean ±SEM. (**F**) Immunoblots showing full-length spike, S2, cleaved S2’ and N collected from Caco-2 cells, which were pre-treated with 2μM IRAK1/4 inhibitor and 10 ng/mL IL-1β for 1 hour, then infected with authentic SARS-CoV-2 for 24 hours. Blots are representative of three independent experiments. Numbers below the blots indicated the intensity of S2’ or N versus β-Actin. (**G**) Immunoblots showing full-length spike, S2, cleaved S2’ and N collected from Calu-3 cells, which were infected with authentic SARS-CoV-2 for 1 hour, then washed with PBS before treated with 2μM IRAK1/4 inhibitor and 10 ng/mL IL-1β for 24 hours. Blots are representative of three independent experiments. Numbers below the blots indicated the intensity of S2’ or N versus β-Actin.

Intriguingly, when we tested TAK1, a downstream molecule of TRAF6 for the potential involvement in the signaling, it was found that IL-1β still reduced bioluminescence signal and S2’ cleavage in TAK1 knockout (sgMAP3K7) HEK293T cell line **(Figure 3—figure supplement 2A)**. Moreover, we found that in the presence of TPCA1, an IKKβ inhibitor, IL-1β still inhibited bioluminescence signal and S2’ cleavage as well **(Figure 3—figure supplement 2B)**. In addition, although IL-1β upregulated the mRNA transcription levels of NF-κB pathway-related genes, such as *RELB*, *NFKBIA*, and *NFKB1* **(Figure 3—figure supplement 2C)**, IL-1β still reduced the bioluminescence signal after these NF-κB pathway-related genes knockout **(Figure 3—figure supplement 2D)**. Taken together, these results demonstrated that IL-1β inhibits SARS-CoV-2 spike induced cell-cell fusion independent from the TAK1-IKKβ-NF-κB signaling cascade.

Furthermore, we validated these findings in authentic SARS-CoV-2 infected Caco-2 and Calu-3 cells. Consist with the results from HEK293T cells, IL-1β failed to reduce S2’ cleavage and N protein in the presence of IRAK1/4 inhibitor, whereas it still reduced S2’ cleavage and N protein in the presence of the IKKβ inhibitor TPCA1 in Caco-2 **(Figure 3F)** and Calu-3 cells **(Figure 3G)**.

### IL-1β inhibits SARS-CoV-2-induced cell-cell fusion through RhoA/ROCK mediated actin bundle formation at the cell-cell junction

It has been reported that IL-1β activates RhoA signaling via MyD88 and IRAK, which is a pathway independent from IKKβ (Chen, Zuraw, Liu, Huang, & Pan, 2002). As a major downstream effector of RhoA, ROCK phosphorylates substrates that are involved in the regulation of the actin cytoskeleton, cell attachment, and cell motility (Riento & Ridley, 2003). Therefore, we set out to detect the active level of RhoA through pull-down assay. To this end, we verified that IL-1β activated RhoA signaling in sgControl HEK293T cells but not in sgMyD88- or sgTRAF6-HEK293T cells **(Figure 4A)**. To directly visualize the distribution of endogenous GTP-RhoA (active RhoA), we used a location biosensor derived from the carboxy terminus of anillin (GFP-AHPH) (Priya et al., 2015; Sun et al., 2015). Interestingly, IL-1β significantly increased the fluorescence intensity of GFP-AHPH in sgControl HEK293T cells, but had no effect in sgMyD88- and sgTRAF6-HEK293T cells **(Figure 4 B and C)**.

**Figure 4.**
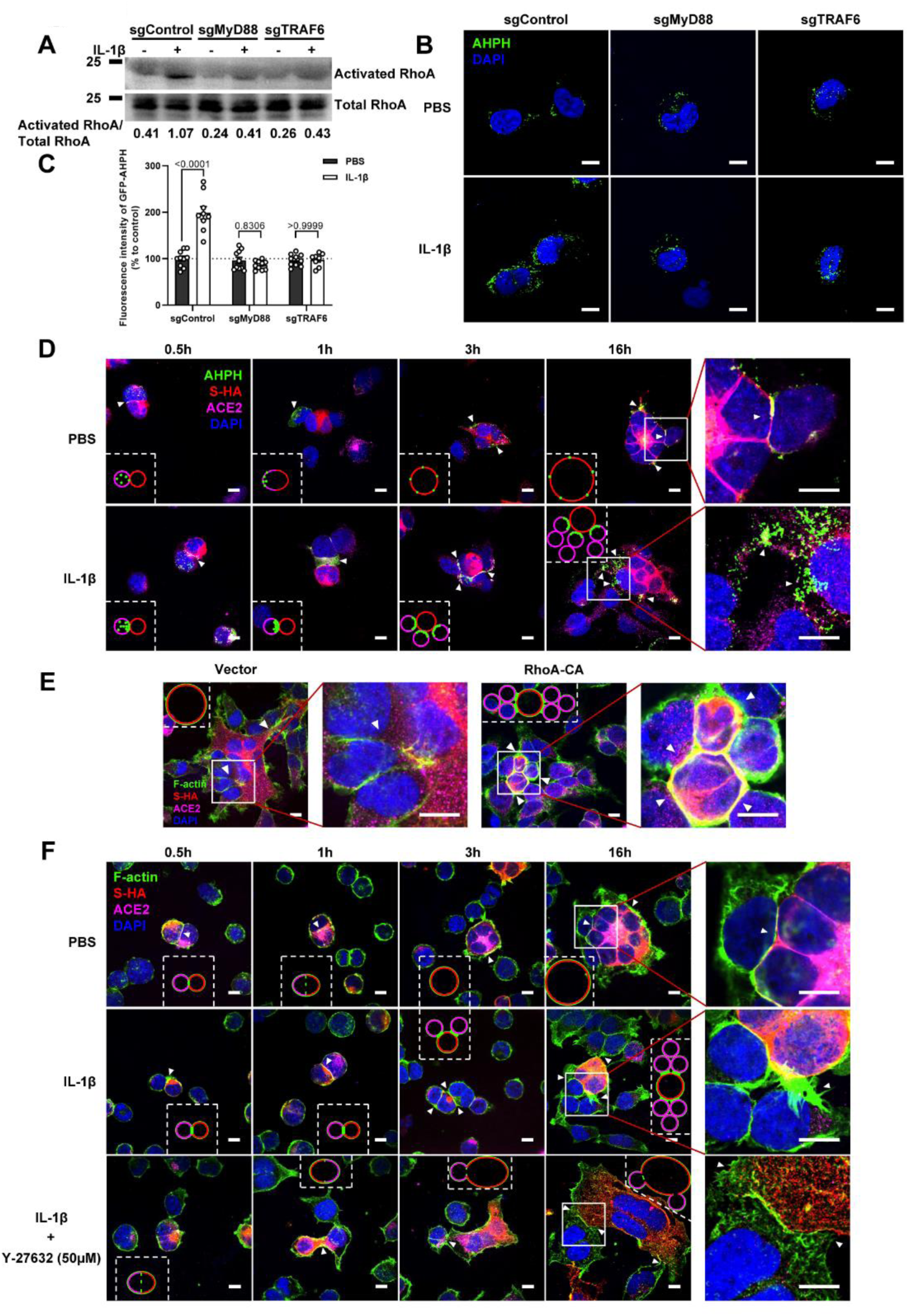
IL-1β inhibits SARS-CoV-2-induced cell-cell fusion through RhoA/ROCK mediated actin bundles assembly at cell-cell junction. (**A**) GTP-RhoA pull-down assay to detect the active level of RhoA in sgControl, sgMyD88 and sgTRAF6 HEK293T cells after 1 ng/mL IL-1β treatment for 30 min. Immunoblots showing activated RhoA and total RhoA. Blots are representative of three independent experiments. Numbers below the blots indicated the intensity of active RhoA versus total RhoA. (**B**) Representative confocal images of GFP-AHPH after 1 ng/mL IL-1β treatment for 30 min in sgControl, sgMyD88 and sgTRAF6 HEK293T cells. Scale bars, 10 μm. (**C**) Quantification of fluorescence intensity of GFP-AHPH in (B). Data are representative of eight individual repeats. (**D**) Representative confocal images of GFP-AHPH localization with or without 1 ng/mL IL-1β treatment at different time points of syncytia formation in HEK293T-S-HA and HEK293T-ACE2 cells. Schematics with green dots in the white dashed line boxes representing GFP-AHPH, red cycles representing S-expressing cells, and magenta cycles representing ACE2-expressing cells. White arrow heads indicate the localization of GFP-AHPH, scale bars, 10 μm. Images are representative of three independent experiments. (**E**) Representative confocal images of F-actin stained with phalloidin-488 in transfected vector or 20 ng RhoA-CA HEK293T-S-HA and HEK293T-ACE2 cells. Schematics with green lines in the white dashed line boxes representing actin bundles, red cycles representing S-expressing cells, and magenta cycles representing ACE2-expressing cells. Scale bars, 10 μm. Images are representative of three independent experiments. (**F**) Representative confocal images of F-actin stained with phalloidin-488 in the presence or absence of 1 ng/mL IL-1β or 50 μM Y-27632 treatment at different time points of syncytia formation in HEK293T-S-HA and HEK293T-ACE2 cells. Schematics with green lines in the white dashed line boxes representing actin bundles, red cycles representing S-expressing cells, and magenta cycles representing ACE2-expressing cells. White arrow heads (E and F) indicate the enrichment or disappearance of F-actin, scale bars, 10 μm. Images are representative of three independent experiments.

To investigate whether IL-1β inhibits SARS-CoV-2 spike-induced cell-cell fusion through the RhoA/ROCK pathway, we co-transfected GFP-AHPH in ACE2-expressing cells, then co-cultured with S-expressing cells at different time points. In the process of syncytia formation, cell-cell contact established between S-expressing cells and ACE2-expressing cells, and GFP-AHPH localized distally from cell-cell junction in the early stage of syncytia formation. With the enlargement of syncytium, GFP-AHPH is visualized at the periphery of syncytium **(Figure 4D, top panel and Figure 4—figure supplement 1A)**. However, in IL-1β treated group, GFP-AHPH foci is enriched to the cell-cell junction in the early stage. Over time, GFP-AHPH was recruited more to the cell-cell junction between S-expressing cells and ACE2-expressing cells, preventing further cell-cell fusion **(Figure 4D, bottom panel and Figure 4—figure supplement 1B)**. Cartoon schematics inserted in the imaging data illustrate such findings in a modeled manner.

It has been reported that RhoA initiates actin arc formation (Dupraz et al., 2019; Stern et al., 2021), so we further explored the changes of actin cytoskeleton during SARS-CoV-2 spike-induced cell-cell fusion. We co-transfected constitutively activated RhoA L63 (Nobes & Hall, 1999) (RhoA-CA) plasmid with spike or ACE2 in HEK293T cells, and found that constitutive activation of RhoA enriches actin filaments (F-actin) at cell-cell junction **(Figure 4E and Figure 4—figure supplement 2A)** and clearly reduces the bioluminescence signal and S2’ cleavage in a dose-dependent manner **(Figure 4—figure supplement 2B)**. Moreover, we observed that F-actin at cell-cell junction between S-expressing cells and ACE2-expressing cells was gradually disappeared along with cell-cell fusion in the early stages of syncytia formation. With the formation and enlargement of syncytium, F-actin of syncytium is preferably distributed peripherally **(Figure 4F, top panel and Figure 4—figure supplement 2C)**. However, IL-1β activated RhoA to initiate actin bundles formation at cell-cell junction, the formation of these actin bundles potentially generates barriers and prevents membrane fusion between S-expressing cells and ACE2-expressing cells. Even with the prolonged co-culture time, IL-1β-induced actin bundles formed at cell junctions consistently inhibited further syncytia formation **(Figure 4F, middle panel and Figure 4—figure supplement 2D)**. Of note, ROCK inhibitor Y-27632 prevents the formation of actin bundles (van der Heijden et al., 2008; Watanabe, Kato, Fujita, Ishizaki, & Narumiya, 1999). Here, we found that the ROCK inhibitor Y-27632 treatment prevented the formation of IL-1β-induced actin bundles at cell-cell junctions, thus promoted membrane fusion and cytoplasmic exchange between S-expressing cells and ACE2-expressing cells and restored syncytia formation **(Figure 4F, bottom panel and Figure 4—figure supplement 3A)**.

Importantly, upon authentic SARS-CoV-2 infection, we observed consistent results: immunofluorescence staining showed GFP-AHPH moving to the opposite of cell-cell junction and located peripherally with syncytia formation **(Figure 5A, top panel and Figure 5—figure supplement 1A)**, while upon IL-1β treatment, GFP-AHPH located to the cell-cell junction of infected cells and neighboring cells **(Figure 5A, bottom panel and Figure 5—figure supplement 1B)**. In parallel, staining results showed that F-actin at the cell-cell junction were disassembled during authentic SARS-CoV-2 infection; with the formation of syncytium, F-actin was mainly distributed peripherally. However, actin bundles formed at cell-cell junction upon IL-1β inhibition of membrane fusion and further syncytia formation **(Figure 5B and Figure 5—figure supplement 1C-E)**. Together, these data revealed that IL-1β induced the formation of actin bundles at the cell-cell junction of SARS-CoV-2 infected cells and neighboring cells through RhoA/ROCK pathway, which inhibited SARS-CoV-2 induced cell-cell fusion.

**Figure 5.**
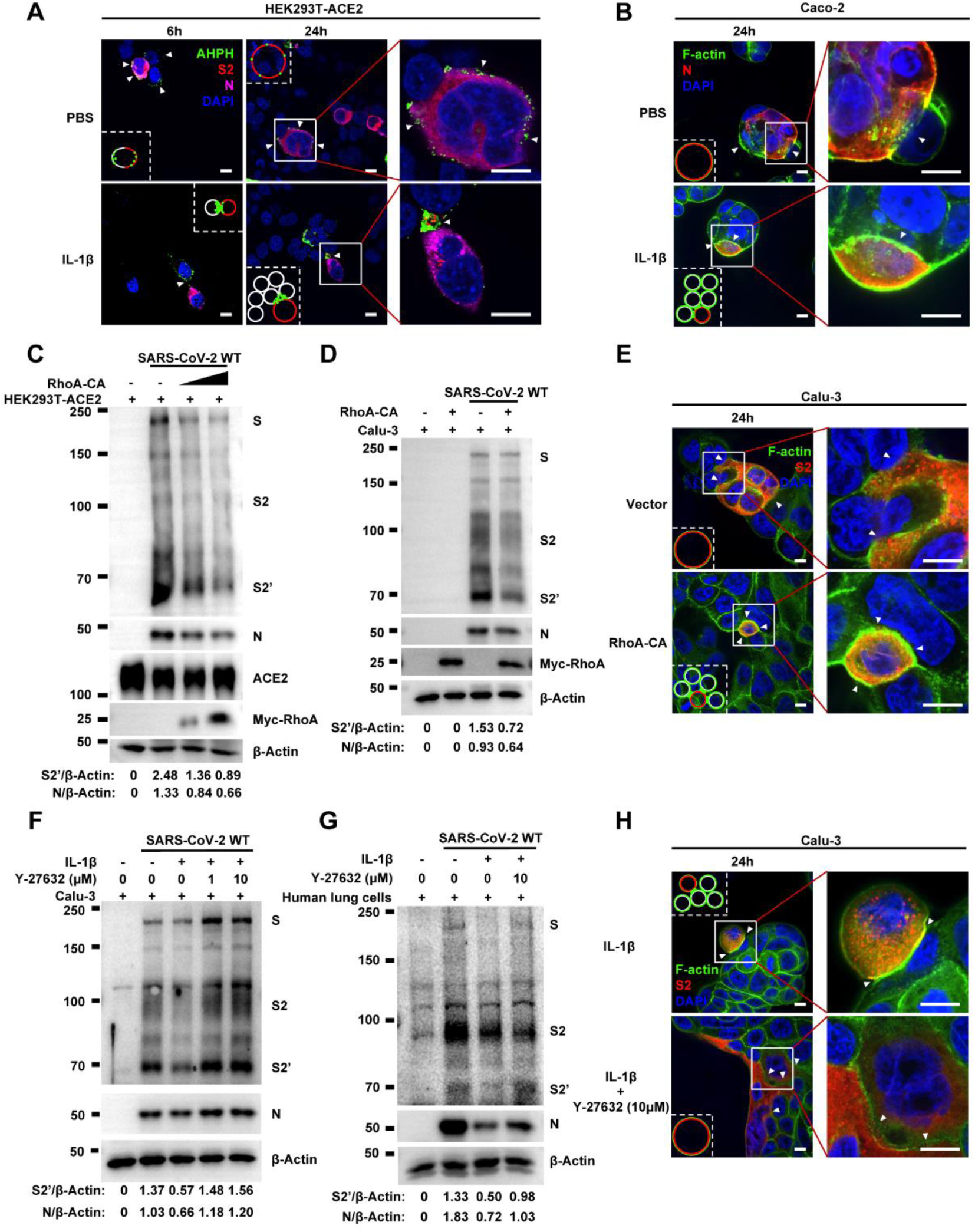
Activation of RhoA/ROCK pathway prevents authentic SARS-CoV-2-induced cell-cell fusion via forming actin bundles. (**A**) Representative confocal images of GFP-AHPH localization with or without 1 ng/mL IL-1β treatment in 0.5 MOI WT authentic SARS-CoV-2 infected HEK293T-ACE2 cells at 6 hpi and 24 hpi. Schematics with green dots in the white dashed line boxes representing GFP-AHPH, red cycles representing SARS-CoV-2 infected cells, and white cycles representing neighboring cells. White arrow heads indicate the localization of GFP-AHPH, scale bars, 10 μm. Images are representative of three independent experiments. (**B**) Representative confocal images of F-actin stained with phalloidin-488 in the presence or absence of 1 ng/mL IL-1β treatment upon 0.5 MOI WT authentic SARS-CoV-2 infection of Caco-2 cells at 24 hpi. Schematics with green lines in the white dashed line boxes representing actin bundles, red cycles representing SARS-CoV-2-infected cells, and white cycles representing neighboring cells. Scale bars, 10 μm. Images are representative of three independent experiments. (**C**) Immunoblots of WT SARS-CoV-2 S, S2, cleaved S2’, N and Myc-RhoA collected from HEK293T-ACE2 cells, which were transfected with vector, 10 ng or 20 ng RhoA-CA before infection with 0.5 MOI authentic SARS-CoV-2 WT strain for 24 hours. Blots are representative of three individual experiments. Numbers below the blots indicated the intensity of S2’ or N versus β-Actin. (**D**) Immunoblots of WT SARS-CoV-2 S, S2, cleaved S2’, N and Myc-RhoA collected from lentivirus-transduced Calu-3 cells expressing vector or RhoA-CA, infected with WT authentic SARS-CoV-2 for 24 hours. Blots are representative of three individual experiments. Numbers below the blots indicated the intensity of S2’ or N versus β-Actin. (**E**) Representative confocal images of F-actin stained with phalloidin-488 from Calu-3 cells described in (D). Schematics with green lines in the white dashed line boxes representing actin bundles, red cycles representing S-expressing cells, scale bars, 10 μm. Images are representative of four independent experiments. (**F, G**) Immunoblots of WT SARS-CoV-2 S, S2, cleaved S2’ and N collected from Calu-3 cells (**F**) or primary human lung cells (**G**), which were infected with authentic SARS-CoV-2 for 1 hour, then washed with PBS before being treated with different concentrations of Y-27632 and 10 ng/mL IL-1β for 24 hours. Blots are representative of three independent experiments. Numbers below the blots indicated the intensity of S2’ or N versus β-Actin. (**H**) Representative confocal images of F-actin stained with phalloidin-488 in Calu-3 cells described in (F). Schematics with green lines in the white dashed line boxes representing actin bundles, red cycles representing S-expressing cells. White arrow heads (B, E and H) indicate the enrichment or disappearance of F-actin, scale bars, 10 μm. Images are representative of four independent experiments.

To further investigate the role of RhoA/ROCK pathway in inhibiting SARS-CoV-2 induced cell-cell fusion, we found that HEK293T-ACE2 **(Figure 5C)**, Caco-2 **(Figure 5—figure supplement 2A)** and Calu-3 cells **(Figure 5D)** expressing RhoA-CA clearly reduced S2’ cleavage and N protein compared to the control group during authentic SARS-CoV-2 infection. Meanwhile, we observed that constitutive activation of RhoA enriches actin bundles at cell-cell junction, thus preventing SARS-CoV-2 induced cell-cell fusion in authentic SARS-CoV-2 infected Caco-2 **(Figure 5— figure supplement 2B and C)** and Calu-3 cells **(Figure 5E and Figure 5—figure supplement 2D)**. In addition, we examined the potential effect of RhoA-CA on ACE2 and found that it did not affect Spike protein binding to ACE2 **(Figure 5—figure supplement 2E)**, nor ACE2 distribution on the cell surface **(Figure 5—figure supplement 2F and G)**. We also observed that IL-1β treatment did not change ACE2 or Spike protein distribution on the cell surface **(Figure 5—figure supplement 3A-D)**.

Notably, ROCK inhibitor Y-27632 treatment increased bioluminescence signal and S2’ cleavage in a dose-dependent manner, promoting syncytia formation. When treated with lower concentrations of Y-27632, IL-1β eliminated Y-27632-enhanced cell-cell fusion. However, IL-1β was unable to inhibit cell-cell fusion in the presence of higher concentrations of Y-27632 **(Figure 5—figure supplement 3E)**. Furthermore, we verified that IL-1β was unable to reduce S2’ cleavage and N protein in the presence of Y-27632 in authentic SARS-CoV-2 infected Caco-2 **(Figure 5—figure supplement 3F)**, Calu-3 cells **(Figure 5F)** and primary human lung cells **(Figure 5G)**. Immunofluorescence results also confirmed that the elimination of IL-1β induced actin bundles by Y-27632 in Caco-2 **(Figure 5—figure supplement 3G and 4A)** and Calu-3 cells **(Figure 5H and Figure 5—figure supplement 4B)**. These results indicated that preventing the formation of RhoA/ROCK mediated actin bundles at cell-cell junction promotes SARS-CoV-2 induced cell-cell fusion.

### IL-1β restricts SARS-CoV-2 transmission via induction of actin bundles *in vivo*

To demonstrate the role of IL-1β in controlling SARS-CoV-2 transmission *in vivo*, BALB/c mice were infected with authentic SARS-CoV-2 B.1.351 after IL-1β or IL-1RA+IL-1β pre-treatment **(Figure 6—figure supplement 1A)**. Interestingly, the results of this experiment showed that in mice with IL-1β treatment, the body weight loss was less than in the PBS control group, while IL-1β was unable to improve body weight in the presence of IL-1RA **(Figure 6—figure supplement 1B)**. According to hematoxylin and eosin (H&E) staining, tissue histopathology analysis demonstrated that the mice with IL-1β treatment carry less pulmonary injury compared to the PBS control and IL-1RA+IL-1β groups **(Figure 6 A and B)**. In addition, the expression level of SARS-CoV-2 N gene in the lung from IL-1β-treated mice was significantly lower than in the PBS control and IL-1RA+IL-1β-treated mice **(Figure 6C)**. In addition, immunohistochemistry staining showed that the infected area in the epithelial linings of lung tissue was significantly reduced by IL-1β treatment compared to the PBS control and IL-1RA+IL-1β groups **(Figure 6 D and E)**, indicating that IL-1β restricted the transmission of SARS-CoV-2 in the lung. Moreover, fluorescence staining showed that SARS-CoV-2 infected lung epithelial cells fused with neighboring cells, promoting viral transmission in the airway epithelial cells, while IL-1β induced the formation of actin bundles to restrict the syncytia formation and further viral transmission **(Figure 6F and Figure 6—figure supplement 1C and D)**. In addition, we found that IL-1β-treated mice have no significant changes in body weight, nor liver and spleen weight compared to control mice **(Figure 6—figure supplement 2A-D)**, indicating that this dose of IL-1β did not cause toxicity *in vivo* in the mice. Of note, when we isolated tissue cells from the IL-1β-treated mice and infected with authentic SARS-CoV-2, it was found that S2’ cleavage and N protein were strongly reduced in IL-1β-treated mice-derived lung and intestine tissue cells compared to control **(Figure 6G and Figure 6—figure supplement 2E)**, suggesting that IL-1β may have protective effects on various tissue cells against SARS-CoV-2 infection *in vivo*.

**Figure 6.**
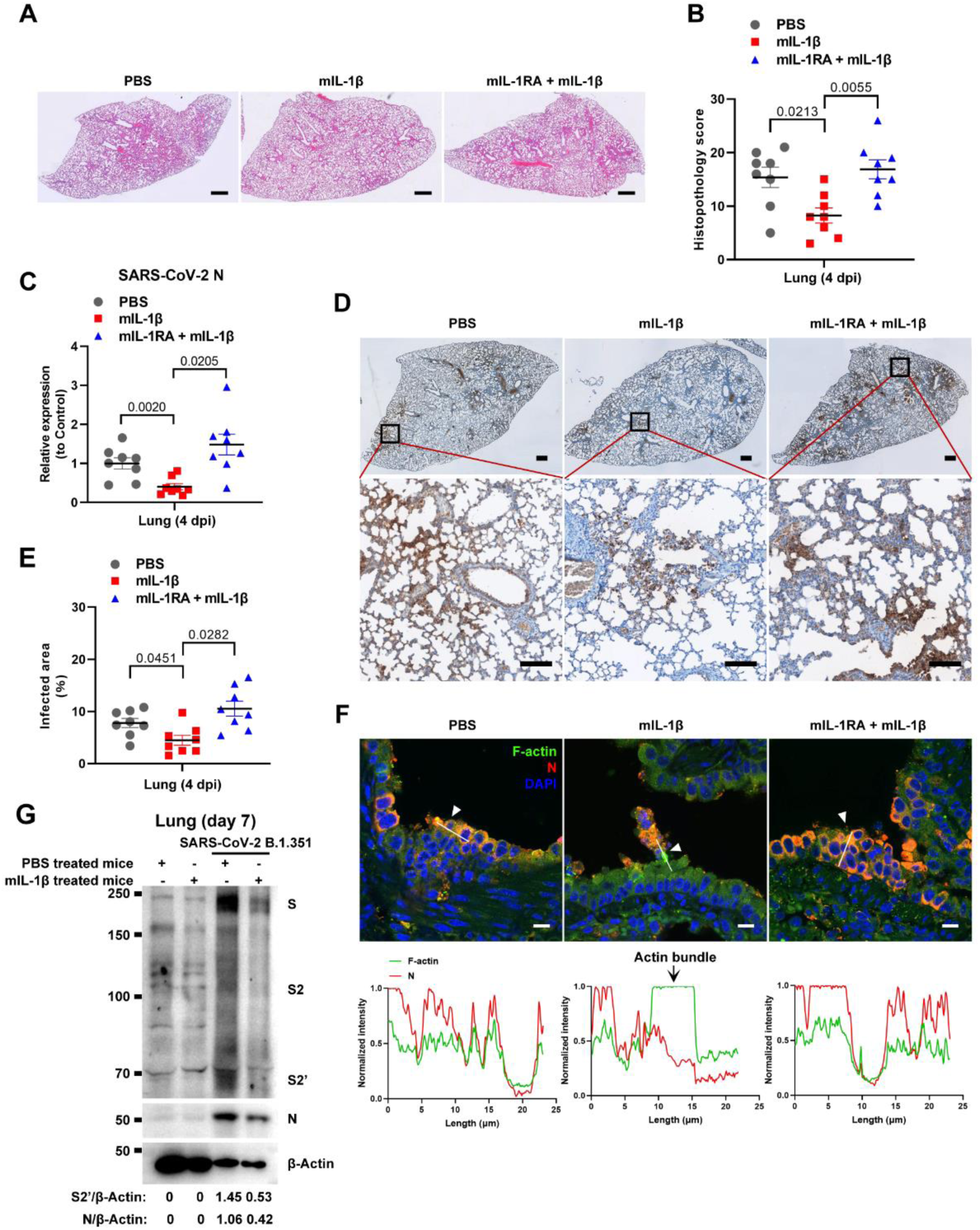
IL-1β restricts SARS-CoV-2 transmission via induction of actin bundles in the lung *in vivo*. (**A**, **B**) Representative images of H&E-stained lung sections (**A**) and histopathology scores (**B**) from PBS; 1 μg/kg mIL-1β; 150 μg/kg mIL-1RA + mIL-1β pre-treated mice infected with SARS-CoV-2 at 4 days post-infection (dpi), scale bars are indicative of 500 μm and images are representative of eight samples. (**C**) qPCR analysis of SARS-CoV-2 N mRNA collected from infected lung tissues at 4 dpi. (**D**) Immunohistochemistry analysis of SARS-CoV-2 N staining in the lung tissue slices at 4 dpi, scale bars are indicative of 500 μm (top panel), 50 μm (bottom panel) and images are representative of eight samples. (**E**) The percentages of SARS-CoV-2-infected area in (D) were quantified. (**F**) Representative confocal images of F-actin stained with phalloidin-488 and SARS-CoV-2 N in the area 1 of lung tissue at 4 dpi. White arrow heads indicate syncytia formation or infected cells, scale bars are indicative of 10 μm and images are representative of three samples (Top). White lines indicate SARS-CoV-2 cell-cell transmission and quantify with fluorescence intensity of F-actin and SARS-CoV-2 N (Bottom). (**G**) Immunoblots of SARS-CoV-2 S, S2, cleaved S2’ and N proteins collected from SARS-CoV-2 B.1.351-infected lung tissue cells, which were isolated from BALB/c mice treated with or without 1 μg/kg mIL-1β at day 7. Blots are representative of three individual mouse. Numbers below the blots indicated the intensity of S2’ or N versus β-Actin.

To further verify the function and mechanism of IL-1β in controlling SARS-CoV-2 transmission *in vivo*, BALB/c mice were infected with authentic SARS-CoV-2 B.1.351 after IL-1β or ROCK inhibitor Y-27632+IL-1β pre-treatment **(Figure 7—figure supplement 1A)**. Similar to IL-1RA, Y-27632 compromised the effect of IL-1β in preventing weight loss **(Figure 7—figure supplement 1B)**. In addition, H&E staining showed that Y-27632 treatment aggravated lung injury in IL-1β-treated mice upon SARS-CoV-2 infection **(Figure 7 A and B)**, although Y-27632+IL-1β did not cause weight loss or lung injury in uninfected mice **(Figure 7—figure supplement 1C and D)**. Moreover, Y-27632 treatment increased the expression level of SARS-CoV-2 N gene **(Figure 7C)** and infected area **(Figure 7D and E)** in the lungs of IL-1β-treated mice. Importantly, Y-27632 treatment prevented the formation of IL-1β-induced actin bundles at cell-cell junctions, thus promoted syncytia formation and further viral transmission **(Figure 7F and Figure 7—figure supplement 2A and B)**. Furthermore, we treated BALB/c mice with PBS, IL-1β or Y-27632 + IL-1β **(Figure 7—figure supplement 2C)**, then isolated the lung tissue cells for authentic SARS-CoV-2 infection. Here it was found that S2’ cleavage and N protein were clearly reduced in IL-1β treated mice compared to control at day 2, while Y-27632 treatment abolished the inhibitory effect of IL-1β **(Figure 7—figure supplement 2D)**. Of note, the lung tissue cells in IL-1β-treated mice remained resistant to SARS-CoV-2 infection at day 7, while the protective effect of IL-1β was abolished by Y-27632 treatment **(Figure 7G)**. Taken together, IL-1β prevents the transmission of SARS-CoV-2 through inducing the formation of actin bundles via the RhoA/ROCK pathway *in vivo*.

**Figure 7.**
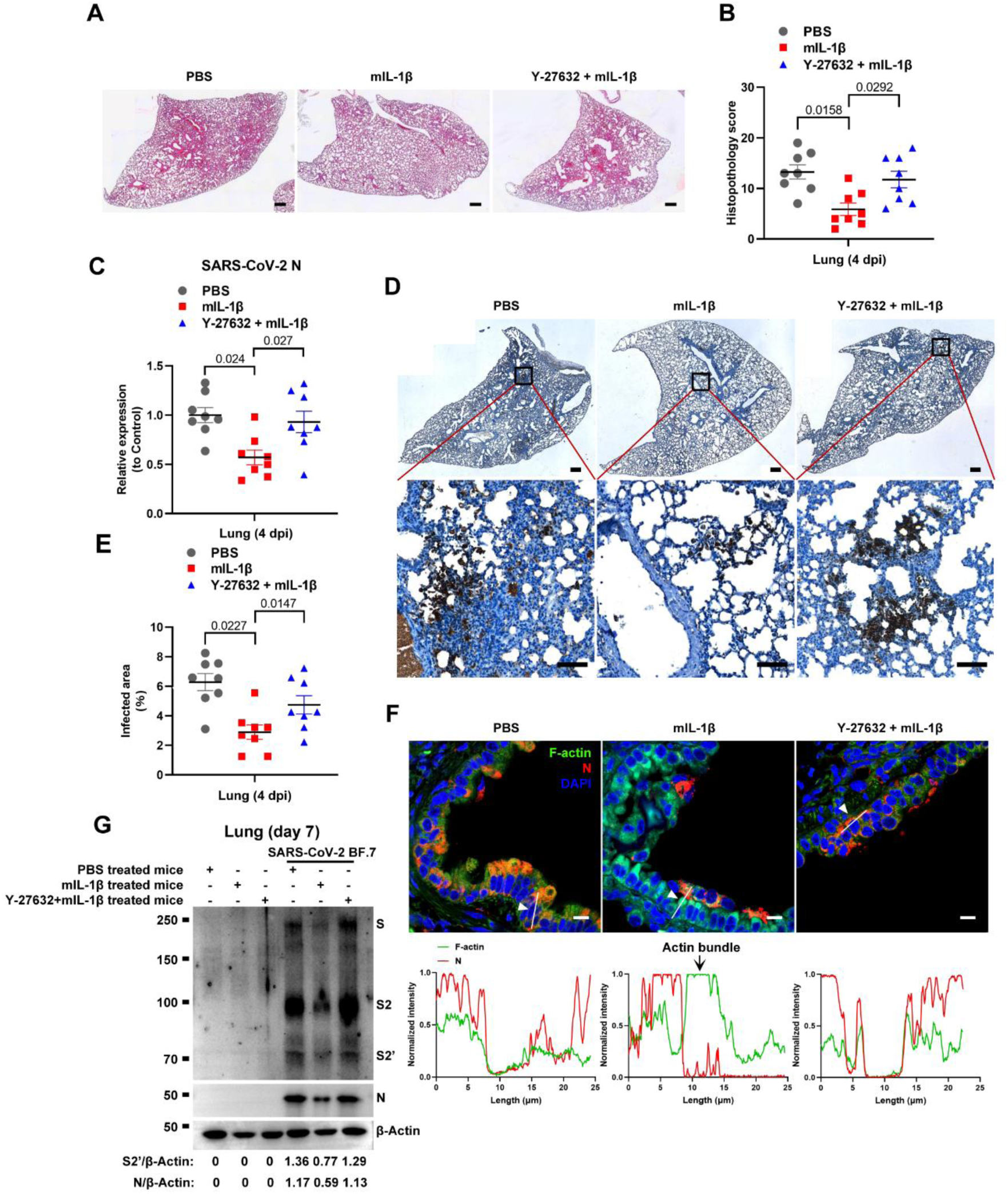
Prevention of IL-1β-induced actin bundles by ROCK inhibitor Y-27632 promotes SARS-CoV-2 transmission *in vivo*. (**A**, **B**) Representative images of H&E-stained lung sections (**A**) and histopathology scores (**B**) from PBS; 1 μg/kg mIL-1β; 1mg/kg Y-27632 + mIL-1β pre-treated mice infected with SARS-CoV-2 at 4 days post-infection (dpi), scale bars are indicative of 500 μm and images are representative of eight samples. (**C**) qPCR analysis of SARS-CoV-2 N mRNA collected from infected lung tissues at 4 dpi. (**D**) Immunohistochemistry analysis of SARS-CoV-2 N staining in the lung tissue slices at 4 dpi, scale bars are indicative of 500 μm (top panel), 50 μm (bottom panel) and images are representative of eight samples. (**E**) The percentages of SARS-CoV-2 infected area in (D) were quantified. (**F**) Representative confocal images of F-actin stained with phalloidin-488 and SARS-CoV-2 N in the area 1 of lung tissue at 4 dpi. White arrow heads indicate syncytia formation or infected cells, scale bars are indicative of 10 μm and images are representative of three samples (Top). White lines indicate SARS-CoV-2 cell-cell transmission and quantify with fluorescence intensity of F-actin and SARS-CoV-2 N (Bottom). (**G**) Immunoblots of SARS-CoV-2 S, S2, cleaved S2’ and N proteins collected from authentic SARS-CoV-2 BF.7-infected lung tissue cells, which were isolated from BALB/c mice treated with PBS, 1 μg/kg mIL-1β or 1mg/kg Y-27632 + 1 μg/kg mIL-1β at day 7. Blots are representative of three individual mouse. Numbers below the blots indicated the intensity of S2’ or N versus β-Actin.

## Discussion

In the present study, we explored the function of innate immune factors against SARS-CoV-2 infection. Notably, IL-1β inhibited various SARS-CoV-2 variants and other beta-coronaviruses spike-induced cell-cell fusion. Mechanistically, IL-1β activates and enriches RhoA to the cell-cell junction between SARS-CoV-2-infected cells and neighboring cells via the IL-1R-mediated signal to initiate actin bundle formation, preventing cell-cell fusion and viral spreading (**Figure 7—figure supplement 3**). These findings revealed a critical function for pro-inflammatory cytokines to control viral infection.

Elevated IL-1β levels in severe COVID-19 patients is central to innate immune response as it induces the expression of other pro-inflammatory cytokines (Tahtinen et al., 2022). In addition, IL-1α is also secreted during SARS-CoV-2 infection (Xiao et al., 2021). Of note, several therapeutic strategies have employed the inhibition of IL-1 signal in an attempt to treat SARS-CoV-2 infection (Huet et al., 2020; Ucciferri et al., 2020). Intriguingly, although anakinra, a recombinant human IL-1 receptor antagonist, improved clinical outcomes and reduced mortality in severe COVID-19 patients (Cavalli et al., 2020), it did not reduce mortality in mild-to-moderate COVID-19 patients, and even increased the probability of serious adverse events (Tharaux et al., 2021). With another note, IL-1 blockade significantly decreased the neutralizing activity of serous anti-SARS-CoV-2 antibodies in severe COVID-19 patients (Della-Torre et al., 2021). According to our finding that both IL-1β and IL-1α are able to inhibit SARS-CoV-2-induced cell-cell fusion, inhibition of IL-1 signaling may have abolished the antiviral function of IL-1, thus failing to restrict virus-induced syncytia formation and transmission.

Notably, IL-1β plays a key role in triggering vaccine-induced innate immunity, suggesting that innate immune responses play important roles in the antiviral defense by enhancing the protective efficacy of vaccines (Eisenbarth, Colegio, O’Connor, Sutterwala, & Flavell, 2008; Tahtinen et al., 2022). In addition, vaccination with Bacillus Calmette-Guérin (BCG) has been reported to confer nonspecific protection against heterologous pathogens, including protection against SARS-CoV-2 infection in humans and mice (Hilligan et al., 2022; A. Lee et al., 2023; Rivas et al., 2021). Moreover, Lipid nanoparticle (LNP) in mRNA vaccine (Han et al., 2023) and penton base in adenovirus vaccine (Di Paolo et al., 2009) can both activate innate immune cells to amplify the protective effect of vaccines, which may also be attributed to IL-1β-mediated inhibition of SARS-CoV-2-induced cell-cell fusion on top of adaptive immune responses induced by the vaccines.

With another note, patients with inherited MyD88 or IRAK4 deficiency have been reported to be selectively vulnerable to COVID-19 pneumonia. It was found that these patients’ susceptibility to SARS-CoV-2 can be attributed to impaired type I IFN production, which do not sense the virus correctly in the absence of MyD88 or IRAK4 (García-García et al., 2023). In our study, MyD88 or IRAK4 deficiency abolished the inhibitory effect of IL-1β on SARS-CoV-2 induced cell-cell fusion, suggesting that these innate immune molecules are critical to contain SARS-CoV-2 infection, and this may be another mechanism accounting for the disease of those patients. Moreover, MyD88 signaling was essential for BCG-induced innate and type 1 helper T cell (TH1 cell) responses and protection against SARS-CoV-2, which is consistent with our fundings.

Of note, cell-cell fusion is not limited to the process of viral infection, both normal and cancerous cells can utilize this physiological process in tissue regeneration or tumor evolution (Delespaul et al., 2020; Powell et al., 2011). For example, myoblast fusion is the key process of skeletal muscle terminal differentiation, inactivation of RhoA/ROCK signaling is crucial for myoblast fusion (Nishiyama, Kii, & Kudo, 2004). Our current work revealed that inhibition of RhoA/ROCK signaling promoted virus-induced cell-cell fusion, possibly due to the virus hijacking of such biological process. In turn, activated RhoA/ROCK signaling inhibits virus-induced cell-cell fusion, so it can be targeted for future therapeutic development to control viral transmission. Cell-cell fusion is mediated by actin cytoskeletal rearrangements, the dissolution of F-actin focus is essential for cell-cell fusion; in contrast, syncytia formation cannot proceed if disassembly of actin filaments or bundles is prevented (Doherty et al., 2011; Rodríguez-Pérez et al., 2021). We uncovered that preventing actin bundles dissolution inhibited virus-induced cell-cell fusion, and IL-1β induced RhoA/ROCK signal promotes actin bundle formation at cell-cell junctions. As RhoA is ubiquitously expressed by all cell types, it is currently unclear whether IL-1-mediated RhoA activation is specific towards viral infection-associated cytoskeleton modification, or may regulate other RhoA-related processes, which is a limitation of the current work and remains to be investigated in future.

In summary, this study demonstrated the function and mechanism of IL-1β in inhibiting SARS-CoV-2 induced syncytia formation, and highlighted the function of innate immune factors including cytokines against coronaviruses transmission, thus provide potential therapeutic targets for viral control.

## Materials and Methods

### Reagents and plasmids

The antibodies used for immunoblotting include: rabbit anti-SARS-CoV-2 S2 (Sino Biological, 40590-T62, 1:2000), mouse anti-SARS-CoV-2 N (Sino Biological, 40143-MM05, 1:1000), rabbit anti-ACE2 (Proteintech, 21115-1-AP, 1:2000), rabbit anti-MERS-CoV S2 (Sino Biological, 40070-T62, 1:1000), rabbit anti-MyD88 (Cell Signaling Technology, 4283, 1:1000), rabbit anti-TRAF6 (Abcam, ab33915, 1:5000), rabbit anti-TAK1 (Cell Signaling Technology, 4505, 1:1000), mouse anti-Myc-Tag (Abclonal, AE010,1:2000), HRP-conjugated β-tubulin (Abclonal, AC030, 1:5000), mouse anti-β-actin (Proteintech, 66009-1-Ig, 1:5000), anti-rabbit/anti-mouse (Jackson Immuno Research, 111-035-003, 1:5000). The antibodies and regents used for immunofluorescence include: rabbit anti-SARS-CoV-2 S2 (Sino Biological, 40590-T62, 1:200), mouse anti-SARS-CoV-2 N (Sino Biological, 40143-MM05, 1:200), rabbit anti-ACE2 (Proteintech, 21115-1-AP, 1:200), mouse anti-HA-Tag (Abclonal, AE008, 1:200). Actin-Tracker Green-488 (Beyotime, C2201S,1:100), goat anti-mouse IgG-555 (Invitrogen, A-21424, 1: 400) and goat anti-rabbit IgG-647 (Invitrogen, A-21236, 1: 400), DAPI (Abcam, ab228549, 1:2000) and antifade mounting medium (vectorlabs, H-1400-10). Purified LTA from S. aureus (Invitrogen, tlrl-pslta), Pam3CSK4 (Invitrogen, tlrl-pms), Peptidoglycan from S. aureus (Sigma-Aldrich, 77140), LPS (Invitrogen, tlrl-eklps), TPCA1 (Selleck, S2824), 5Z-7-Oxozeaenol (Sigma-Aldrich, O9890), IRAK1/4 inhibitor (Selleck, S6598), Y-27632 (Selleck, S6390). Inhibitors were dissolved in dimethyl sulfoxide (DMSO, Sigma-Aldrich, D2650), and DMSO was added as solvent control. Recombinant human IL-1α (200-01A), human IL-1β (200-01B), mouse IL-1β (211-11B), human IL-1RA (200-01RA), human IL-6 (200-06) and human IL-8 (200-08M) were purchased from Peprotech. Mouse IL-1RA (769706) was purchased from BioLegend. IL-1β concentrations in supernatants from THP-1 and PBMCs were determined using ELISA kit, according to the manufacturer’s instructions (R&D Systems, DY201). RhoA pull-down activation assay Biochem kit (BK036-S) was purchased from Cytoskeleton, Inc. Collagenase, type I (17100017) was purchased from Gibco and B-ALI Growth Media (00193516) was purchased from Lonza Bioscience.

SARS-CoV-2 spike (Wild type, GenBank: QHD43419.1) was homo sapiens codon-optimized and generated *de novo* into pVAX1 vector by recursive polymerase chain reaction (PCR). WT, Alpha, Beta and Delta variants containing point and deletion mutations were generated via stepwise mutagenesis using spike construct containing the truncated 19 amino acids at the C-terminal (CTΔ19). The latest human codon optimized Omicron was purchased from Genescripts, and subcloned into the pVAX1 backbone with CTΔ19 for comparison. Human ACE2 assembled in a pcDNA4.0 vector was used for transient expression of ACE2. GFP-AHPH (Addgene plasmid # 71368; http://n2t.net/addgene:71368; RRID: Addgene_71368) and pRK5myc RhoA L63 (Addgene plasmid # 15900; http://n2t.net/addgene:15900; RRID: Addgene_15900) were from Addgene.

### Cell culture and stimulation

HEK293T cells were purchased from the National Science & Technology Infrastructure (NSTI) cell bank (www.cellbank.org.cn). Human colon epithelial carcinoma cell line Caco-2 (catalog no. SCSP-5027) cells were obtained from Cell Bank/Stem Cell Bank, Chinese Academy of Sciences. Human lung cancer cell line Calu-3 and Vero E6-ACE2 cells were gifted from Prof. Dimitri Lavillette (Applied Molecular Virology Laboratory, Discovery Biology Department, Institut Pasteur Korea). Human monocytic cell line THP-1 (TIB-202; ATCC) was authenticated at Genetic Testing Biotechnology Corporation (Suzhou, China) using Short Tandem Repeat (STR) analysis as described in 2012 in ANSI Standard (ASN-0002) by the ATCC Standards Development Organization. HEK293T and Vero E6-ACE2 cells were cultured in Gibco Dulbecco’s Modified Eagle Medium (DMEM) (GE Healthcare) supplemented with 10% fetal bovine serum (FBS) (Sigma) and 1% Penicillin/streptomycin (P/S) (Life Technologies) at 37°C with 5% CO_2_ in a humidified incubator. Caco-2 and Calu-3 cells were cultured in Minimum Essential Medium (MEM) supplemented with 10% FBS, 1% non-essential amino acids and 1% P/S at 37°C with 5% CO_2_ in a humidified incubator. THP-1 cells were cultured in Roswell Park Memorial Institute (RPMI) 1640 supplemented with 10% FBS, 1% P/S and 50 μM 2-ME at 37°C with 5% CO_2_ in a humidified incubator. All cells were routinely tested for mycoplasma contamination; passages between 4 ^th^ to 25 ^th^ were used. Human PBMCs were isolated from the peripheral blood of healthy doners (Shanghai Blood Center). This study was performed in accordance with the International Ethical Guidelines for Biomedical Research Involving Human Subjects and the principles expressed in the Declaration of Helsinki. Briefly, fresh human PBMCs were separated using Ficoll-Paque PLUS reagent (cytiva, 17144003) at 1200 g for 10 min at room temperature with SepMateTM-50 (SepMate, 86450). PBMCs were washed three times with filtered PBS containing 0.5% BSA and 2 mM EDTA. PBMCs were counted and resuspended in RPMI 1640 medium supplemented with 1% FBS and 1% P/S.

For stimulation, THP-1 cells were seeded at 2 × 10^6^ cells per ml in FBS free RPMI 1640 and PBMCs were seeded at 1 × 10^7^ cells per ml in 1% FBS RPMI 1640, then stimulated with LTA (10 μg/ml), Pam3CSK4 (1 μg/ml), PGN (2 μg/ml), LPS (1 μg/ml) for 24 hours, cell culture supernatants were collected after centrifugation at 2000 g for 5 min for subsequent experiments.

### Transient transfection and cell-cell fusion assays

For transient transfections, HEK293T cells were seeded in 24-well plates at 0.5 x 10^6^ cells /mL overnight. 250 ng plasmids encoding SARS-CoV-2 spike mutants or ACE2 variants were packaged in Lipofectamine 2000 (Life technologies) and transfected for 24 hours. For luciferase assays, Spike-mediated membrane fusion, a *Cre-loxp* Firefly luciferase (*Stop-Luc*) co-expression system was introduced to enable the detection of DNA recombination events during cell-cell fusion. 200 ng Cre plasmids were co-transfected into HEK293T-S cells and 200 ng *Stop-Luc* plasmid were co-transfected into HEK293T ± ACE2 cells, respectively. For visualization of syncytia formation, 100 ng ZsGreen plasmid was co-transfected with spike variants. HEK293T cells in the 24-well plates were then detached using ice-cold calcium-free PBS in the absence of trypsin and centrifuged at 600 g for 4 min.

For cell-cell fusion assays, cell pellets were resuspended into complete DMEM and mixed with control HEK293T cells, or HEK293T-ACE2, Vero E6-ACE2 or Calu-3 cells at 1:1 ratio before adhesion to the 48-well or 96-well plates, cell mixes were incubated for 16 hours at 37°C. Quantification of cell-cell fusion was performed by measuring luciferase expression as relative luminescence units (RLU) 1 min by mixing cell lysates with the Bright-Glo luciferase substrate (E2610, Promega) on a Synergy H1 plate reader (Biotek). Fluorescent images showing syncytia formation were captured at endpoint using a 10x objective and 12-bit monochrome CMOS camera installed on the IX73 inverted microscope (Olympus). Attached cells and syncytia were lysed in a NP40 lysis buffer containing 0.5% (v/v) NP40, 25 mM Tris pH 7.3, 150 mM NaCl, 5% glycerol and 1x EDTA-free protease inhibitor cocktail (PIC) (Roche).

### Immunoblotting

Tissue culture plates containing adherent syncytia and cell mixes were directly lysed on ice in 2x reducing Laemmli loading buffer before boiled at 95°C for 5 min. Protein samples were separated by standard Tris-glycine SDS-PAGE on 7.5% or 9.5% Tris-glycine polyacrylamide gels. Proteins were then transferred onto 0.45 μm PVDF membranes (Millipore) for wet transfer using Towbin transfer buffer. All membranes were blocked in PBS supplemented with 0.1% Tween20 (PBST) and 2.5% bovine serum albumin (BSA) or 5% non-fat dry milk, before overnight incubation in primary antibodies at 4°C. Blots were labelled with HRP-tagged secondary antibodies (Jackson ImmnuoResearch) and visualized with PicoLight substrate enhanced chemiluminescence (ECL) solution (Epizyme Scientific). Immunoblot images were captured digitally using a 5200 chemiluminescent imaging system (Tanon) with molecular weight markers indicated.

### Real time PCR

0.5 x 10^6^ cells /mL HEK293T cells were seeded in 24 well plates overnight. After the cells were about 80% covered, specified stimulant was added. Upon harvesting, cells were washed with PBS for three times, and 1 mL TRIzol Reagent (15596018; Thermo Fisher Scientific) was added for full lysis at room temperature for 5 min. 250 μL chloroform was added, fully mixed at room temperature for 5 min, centrifuged at 10000 r/min, 4°C for 10 min. After carefully removing the aqueous phase using a pipette into another 1.5 mL Eppendorf tube, some of the aqueous phase (about 1 mm above DNA layer to prevent DNA contamination) was remained. 550 µL isopropanol was added in the aqueous phase and mixed gently, then placed at −20°C for 30 min. The tubes were centrifuged at 14000 r/min, 4°C for 20 min, and washed with 75% ethanol twice before dissolved in 30 µL DEPC water. RNA was reverse transcribed to cDNA using a GoSript Reverse Transcription Kit (Promega). Real-time PCR was performed using SYBR Green Realtime PCR Master Mix (TOYOBO) on ABI QuantStudio 6 flex Real-time PCR System (Thermo Fisher Scientific). The RT-qPCR Primer sequences for targeting genes are displayed in Table S1. Target genes’ relative quantification was normalized to GAPDH as relative unit (RU).

### CRISPR/Cas9-Mediated Gene Targeting

Gene-deficient THP-1 or HEK293T cells were generated using CRISPR/Cas9-mediated gene targeting technology. Briefly, LentiCRISPR v2 (52961; addgene) containing sgRNA specifically targeting indicated genes were constructed. The sgRNA sequences for targeting respective genes are displayed in Table S2. The Lentiviral particles were produced in HEK293T cells by transfection with LentiCRISPR v2-sg gene, psPAX2, VSV-G at 2:1.5:1 ratio using Lipofectamine 2000. The lentiviral particles were employed to infect THP-1 or HEK293T cells. One day post infection, the cells were subjected to puromycin selection at a concentration of 2 μg/ml for 72 hours. Survived cells were subjected to limiting dilution in 96-well plates to obtain single clones stably knocking-out respective genes.

### RhoA pull-down assay

RhoA pull-down activation assay Biochem kit was applied for this experiment. In brief, after 1 ng/mL IL-1β treatment for 30 min, HEK293T cells were placed on ice and the culture media was aspirated off before washing cells with ice cold PBS, then washed cells were transferred into 1.5 mL Eppendorf tubes followed with a centrifugation 600 g, 4°C for 5 min. Then the cell lysis buffer with protease inhibitor cocktail was added. The tubes were immediately centrifuged at 10000 g, 4°C for 1 min, then 20 µL of the lysate was saved for total RhoA, and the remaining lysate was used for pull-down assay. For pull-down assay, 10 μL rhotekin-RBD beads were mixed with 600 µg total protein, then the tubes were incubated at 4°C on a rotator for 1 hour before centrifuged at 5000 g, 4°C for 1 min. Next, 90% of the supernatant were carefully removed before washing beads with 500 μL wash buffer. Then the tubes were centrifuged at 5000 g, 4°C for 3 min, and supernatant was carefully removed before adding 20 μL of 2 x Laemmli sample buffer, then the beads were thoroughly resuspended and boiled for 2 min and analyzed through Immunoblotting.

### Immunostaining and confocal microscopy

HEK293T-ACE2 cells were seeded onto sterilized poly-D-lysine (100 ug/mL) (Beyotime, ST508) treated 12 mm coverslips (fisher scientific, 1254580) in 24-well plates. After co-culture with HEK293T-S, cells were washed with PBS once before fixing with 4% (w/v) paraformaldehyde (PFA) for 20 min. Then, cells were washed twice with PBS and permeabilized with 0.1% Triton at room temperature for 10 min (For WGA staining, cells were not treated with Triton). Next, cells were washed twice with PBS and blocked with Immunol Staining Blocking Buffer (Beyotime, P0102) at room temperature for 1 hour. Primary antibodies were incubated at room temperature for 1 hour. Coverslips were then washed twice with PBS before incubation with Actin-Tracker Green-488 or secondary antibodies for 1 hour at room temperature. Coverslips were washed twice with PBS before DAPI staining for 10 min or being mounted in antifade mounting medium. Fluorescent images covering various areas on the coverslips were captured at 12-bit depth in monochrome using a 100x oil immersion objective on the Olympus SpinSR10 confocal microscope and subsequently processed using imageJ software (NIH) with scale bars labeled.

### Authentic SARS-CoV-2 Infection of cells

All experiments involving authentic SARS-CoV-2 virus *in vitro* were conducted in the biosafety level 3 (BSL3) laboratory of the Shanghai municipal center for disease control and prevention (CDC). The experiments and protocols in this study were approved by the Ethical Review Committee of the Shanghai CDC. Briefly, HEK293T-ACE2 or Caco-2 cells were seeded into 24-well or 96-well plates at a density of 4 x 10^5^ cells per mL overnight, then pre-treated with different reagents for 1 hour before infection with 0.5 multiplicity of infection (MOI) Delta or WT authentic SARS-CoV-2 (B.1.617.2 and WT) for 24 hours. Calu-3 cells were seeded into 24-well or 96-well plates at a density of 4 x 10^5^ cells per mL overnight, infected with 0.5 MOI WT authentic SARS-CoV-2 for 1 hour, then washed with PBS before treating with different reagents for 24 hours. Brightfield images were captured to indicate the syncytia formation, cell lysates were collected for spike S2’ cleavage and N protein immunoblots.

For primary mouse tissue cells, specific pathogen-free 6-week-old female BALB/c mice were lightly anesthetized with isoflurane and intranasal treated with PBS, mIL-1β (1 μg/kg) or Y-27632 (1 mg/kg) + mIL-1β (1 μg/kg) at day 0, then mice were Intraperitoneal injected with PBS, mIL-1β (1 μg/kg) or Y-27632 (1 mg/kg) + mIL-1β (1 μg/kg) at day 1 and 2. At day 7, mice were anesthetized by intraperitoneal injection of Avertin (2,2,2-tribromoethanol, Sigma-Aldrich), the thoracic cavity and abdominal cavity were opened, an outlet was cut in the left ventricle of the mice, and then the right ventricle was perfused with phosphate buffer saline (PBS) through the pulmonary artery to remove blood cells in the lung. Next, the lung digestive solution with HBSS 1ml, 1 mg/ml collagenase IA, DnaseI (200 mg/ml; Roche), DispaseII (4 U/ml; Gibco) and 5% FBS was injected into the lung cavity, then the lungs were peeled off and digested for 30 min with shaking at 37°C. For the intestinal cell isolation, the intestines are peeled off and put into the intestine digestive solution containing DMEM 10mL, 1 μM DTT, 0.25 μM EDTA and 5% FBS, then digested for 30 min with shaking at 37°C. After digestion, the lung and intestinal cells were resuspended in DMEM supplemented with 10% FBS and 1% P/S, subsequently, the cells were infected with 1 MOI authentic SARS-CoV-2 B.1.351 or BF.7 for 24 hours. All procedures were conducted in compliance with a protocol approved by the Institutional Animal Care and Use Committee (IACUC) at Shanghai Institute of Immunity and Infection, Chinese Academy of Sciences.

For primary human lung cells, the human lung tissues were cut into small pieces of about 2 mm^3^ and washed three times with HBSS solution containing 1% PS, digested with collagenase type I (100 mg+50 mL PBS) in an incubator at 37°C for 4 hours. Then filtered through a 70 μm filter and centrifuged at 500 g for 5 min at room temperature. Lysed with 3 mL Red Blood Cell Lysis Buffer for 5min at room temperature, and then centrifuged at 500 g for 5 min at room temperature, washed twice with HBSS solution containing 1% PS. Human lung cells were resuspended in B-ALI Growth Media and seeded into 96-well plates at a density of 4 x 10^5^ cells per mL overnight, infected with 0.5 MOI WT authentic SARS-CoV-2 for 1 hour, then washed with PBS before treating cells with different reagents for 24 hours. The experiments and protocols were approved by the Ethical Review Committee of the Shanghai CDC.

### Authentic SARS-CoV-2 infection of BALB/c mice

Specific pathogen-free 6-week-old female BALB/c mice were lightly anesthetized with isoflurane and intranasal treated with PBS, mIL-1β (1 μg/kg), mIL-1RA (150 μg/kg) + mIL-1β (1 μg/kg); or PBS, mIL-1β (1 μg/kg), Y-27632 (1 mg/kg) + mIL-1β (1 μg/kg) for 1 hour, then intranasal challenged with 5 × 10^4^ FFU of SARS-CoV-2 B.1.351. For booster injection, mice were Intraperitoneal injected with PBS, mIL-1β (1 μg/kg), mIL-1RA (150 μg/kg) + mIL-1β (1 μg/kg); or PBS, mIL-1β (1 μg/kg), Y-27632 (1 mg/kg) + mIL-1β (1 μg/kg) at 1- and 2-days post infection (dpi). Mice were monitored daily for weight loss. Lungs were removed into Trizol or 4%PFA at 4 dpi. All protocols were approved by the Institutional Animal Care and Use Committee of the Guangzhou Medical University.

### Pulmonary histopathology

Lungs were collected from mice infected with SARS-CoV-2 at 4 dpi and fixed in 4% PFA (Bioss) for 12 hours followed by dehydrating, embedded in paraffin for sectioning, then stained with hematoxylin and eosin (H&E), immunohistochemistry (IHC) or immunofluorescence (IF). H&E and IHC data were analyzed by PerkinElmer Vectra 3, IF results were analyzed by Olympus SpinSR10 confocal microscope. The pathological scores were judged according to previous work (Curtis, Warnock, Arraj, & Kaltreider, 1990).

## Statistics analysis

Bar graphs were presented as mean values ± standard error of mean (SEM) with individual data points. All statistical analyses were carried out with the Prism software v8.0.2 (GraphPad). Data with multiple groups were analyzed using matched one-way ANOVA followed by Sidak’s post hoc comparisons. Statistical significance P values were indicated between compared groups and shown on Figures.

## Acknowledgments

We thank Qiuhong Guo, Prof. Dimitri Lavillette and Prof. Gary Wong for their experimental supports and reagents used in this work. This study is supported by grants from Natural Science Foundation of China (92269202, 82825001, 92054104, 82402593), National Key R&D Program of China (2022YFC2304700, 2022YFC2303200, 2022YFC2303502), Strategic Priority Research Program of the Chinese Academy of Sciences (XDB0940102),Three-Year Initiative Plan for Strengthening Public Health System Construction in Shanghai (2023-2025) Key Discipline Project (GWVI-11.1-09), Shanghai Municipal Science and Technology Major Project (2019SHZDZX02), Shanghai Natural Science Foundation Projects (24ZR1476700, 24ZR1476600), State Key Laboratory of Respiratory Disease Project (J24411029), Guangdong Basic and Applied Basic Research Foundation (2023A1515010152) and Young Scientists Fund of the Guangzhou National Laboratory (QNPG23-03).

## Supplemental Figures

**Figure 1—figure supplement 1.**
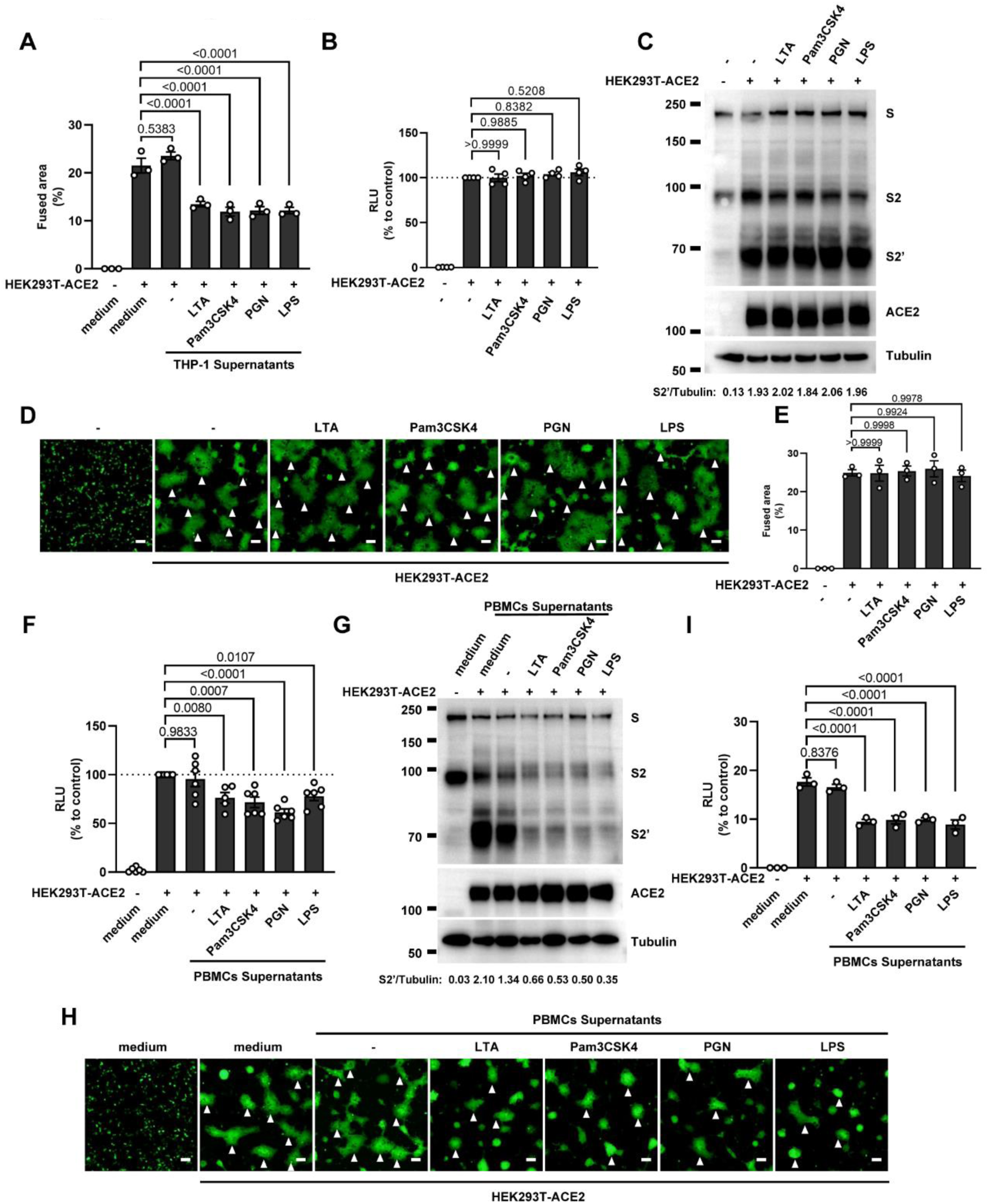
Host factors secreted by activated PBMCs inhibit SARS-CoV-2 spike-induced cell-cell fusion, but TLR ligands alone have no effect. (**A**) Quantification of the fused area in Figure 1D. (**B**) Luciferase activity (RLU) measured from HEK293T cell lysates collected from TLR ligands treated HEK293T-S and HEK293T-ACE2 for 16 hours. Data are representative of four individual repeats and displayed as individual points with mean ± standard error of the mean (SEM). (**C**) Immunoblots showing full-length spike, S2, cleaved S2’ and ACE2 collected from TLR ligands-treated HEK293T-S and HEK293T-ACE2 for 16 hours. Blots are representative of three independent experiments. Numbers below the blots indicated the intensity of S2’ versus Tubulin. (**D**) Representative fluorescent image captured at 488 nm from TLR ligands treated HEK293T-S-Zsgreen and HEK293T-ACE2 for 16 hours. (**E**) Quantification of the infected area in (D). (**F**) Luciferase activity (RLU) measured from HEK293T cell lysates collected from PBMCs supernatants-treated HEK293T-S and HEK293T-ACE2 for 16 hours. 1% FBS RPMI 1640 served as medium control. Data are representative of five individual repeats and displayed as individual points with mean ±SEM. (**G**) Immunoblots showing full-length spike, S2, cleaved S2’ and ACE2 collected from PBMCs supernatants-treated HEK293T-S and HEK293T-ACE2 for 16 hours. Blots are representative of three independent experiments. Numbers below the blots indicated the intensity of S2’ versus Tubulin. (**H**) Representative fluorescent image captured at 488 nm from PBMCs supernatants-treated HEK293T-S-Zsgreen and HEK293T-ACE2 for 16 hours. White arrow heads (D and H) indicate syncytia formation. Scale bars, 50 μm. Images are representative of three individual repeats. (**I**) Quantification of the fused area in (H).

**Figure 1—figure supplement 2.**
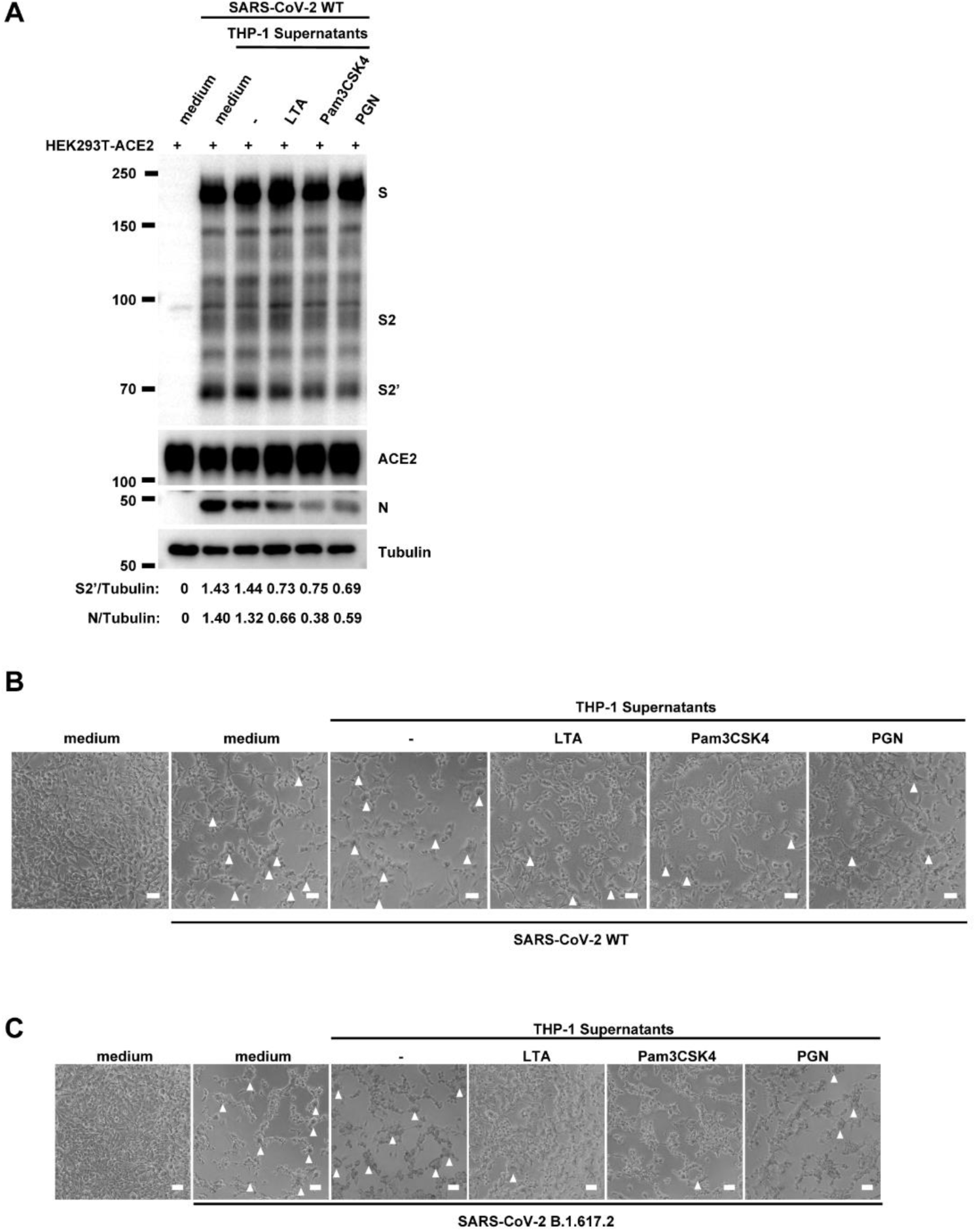
Host factors secreted by activated THP-1 cells inhibit authentic SARS-CoV-2-induced cell-cell fusion. (**A**) Immunoblots of WT SARS-CoV-2 S, S2, cleaved S2’ and N proteins collected from HEK293T-ACE2 cells 24 hpi as described in Figure 1E. Blots are representative of three individual experiments. Numbers below the blots indicated the intensity of S2’ or N versus Tubulin. (**B, C**) Bright field images of 0.5 MOI WT (**B**) or Delta (**C**) SARS-CoV-2-infected HEK293T-ACE2 cells pre-treated with THP-1 supernatants. White arrow heads indicate syncytia formation, scale bars are indicative of 50 μm, and images are representative of three independent experiments.

**Figure 2—figure supplement 1.**
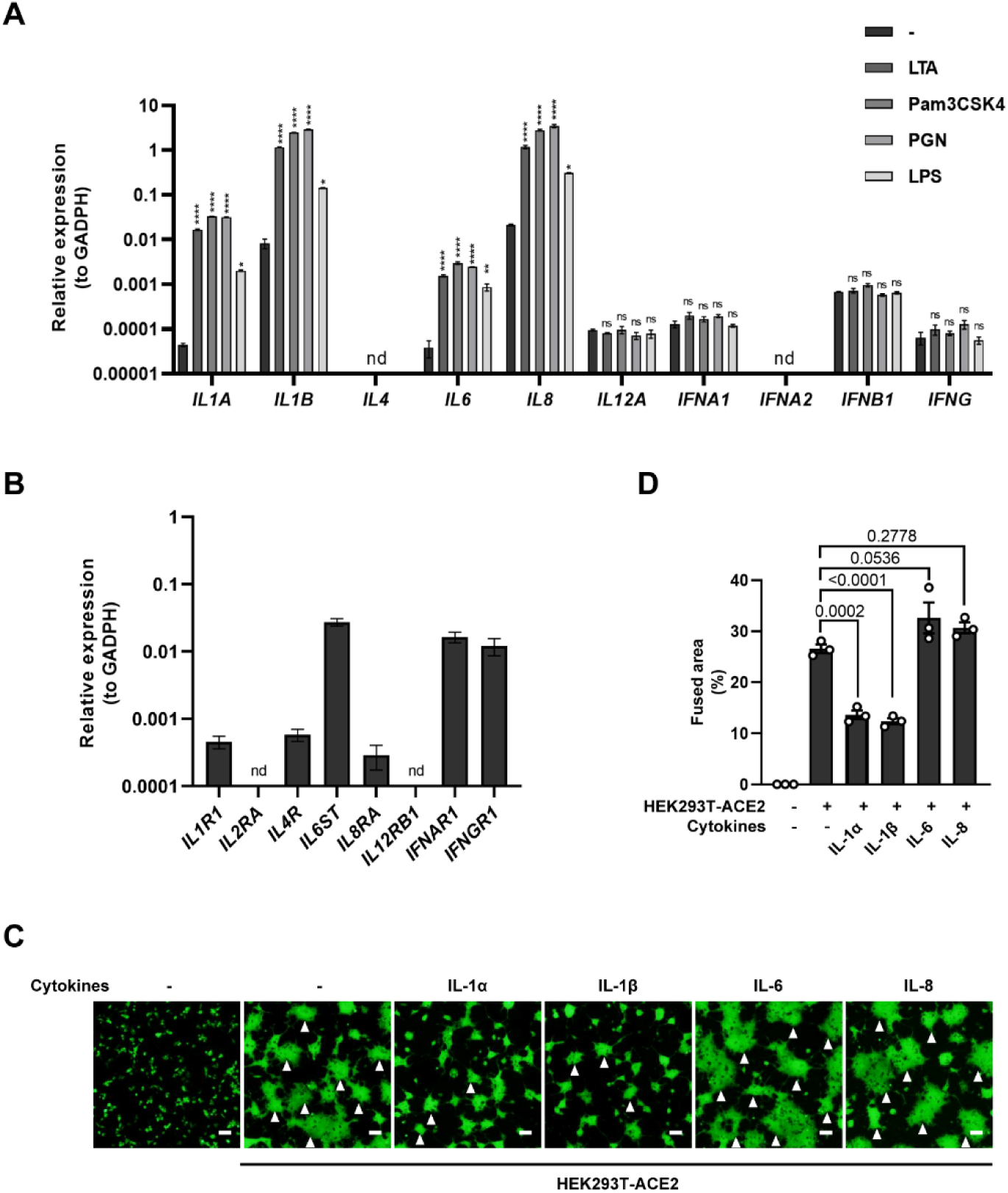
mRNA levels of different cytokine genes in THP-1 cells and selected cytokine receptor genes in HEK293T cells, as well as fluorescent images indicating cell-cell fusion in the presence of indicated cytokines. (**A**) mRNA levels of different cytokine genes in THP-1 cells after TLR ligands stimulation for 4 hours. Data are representative of three individual repeats. (**B**) mRNA levels of indicated cytokine receptor genes in HEK293T cells. Data are representative of five individual repeats. (**C**) Representative fluorescent image captured at 488 nm from different cytokines-treated HEK293T-S-Zsgreen and HEK293T-ACE2 for 16 hours. White arrow heads indicate syncytia formation. Scale bars, 50 μm. Images are representative of three individual repeats. (**D**) Quantification of the fused area in (C).

**Figure 2—figure supplement 2.**
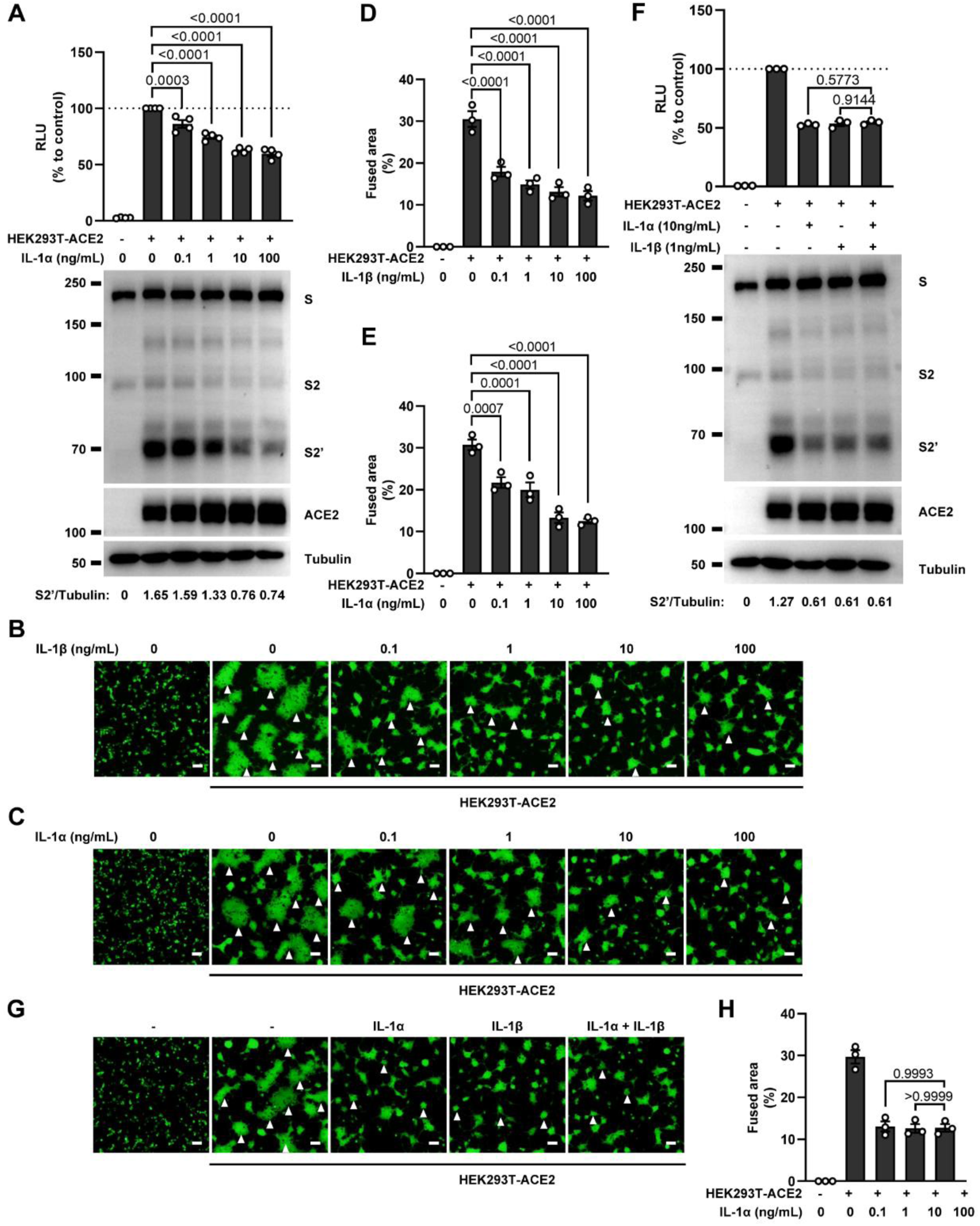
No synergistic inhibition of SARS-CoV-2 spike-induced cell-cell fusion by IL-1α and IL-1β co-treatment. (**A**) Luciferase activity (RLU) measured from HEK293T cell lysates and immunoblots showing full-length spike, S2, cleaved S2’ and ACE2 collected from different concentrations of IL-1α-treated HEK293T-S and HEK293T-ACE2 for 16 hours. Data and blots are representative of four individual repeats. Numbers below the blots indicated the intensity of S2’ versus Tubulin. (**B, C**) Representative fluorescent image captured at 488 nm from different concentrations of IL-1β (**B**) or IL-1α (**C**) treated HEK293T-S-Zsgreen and HEK293T-ACE2 for 16 hours. (**D**) Quantification of the fused area in (B). (**E**) Quantification of the fused area in (C). (**F**) Luciferase activity (RLU) measured from HEK293T cell lysates and immunoblots showing full-length spike, S2, cleaved S2’ and ACE2 collected from IL-1α and IL-1β co-treated HEK293T-S and HEK293T-ACE2 for 16 hours. Data and blots are representative of three individual repeats. Numbers below the blots indicated the intensity of S2’ versus Tubulin. (**G**) Representative fluorescent image captured at 488 nm from IL-1α and IL-1β co-treated HEK293T-S-Zsgreen and HEK293T-ACE2 for 16 hours. White arrow heads (B, C and G) indicate syncytia formation. Scale bars, 50 μm. Images are representative of three individual repeats. (**E**) Quantification of the fused area in (G).

**Figure 2—figure supplement 3.**
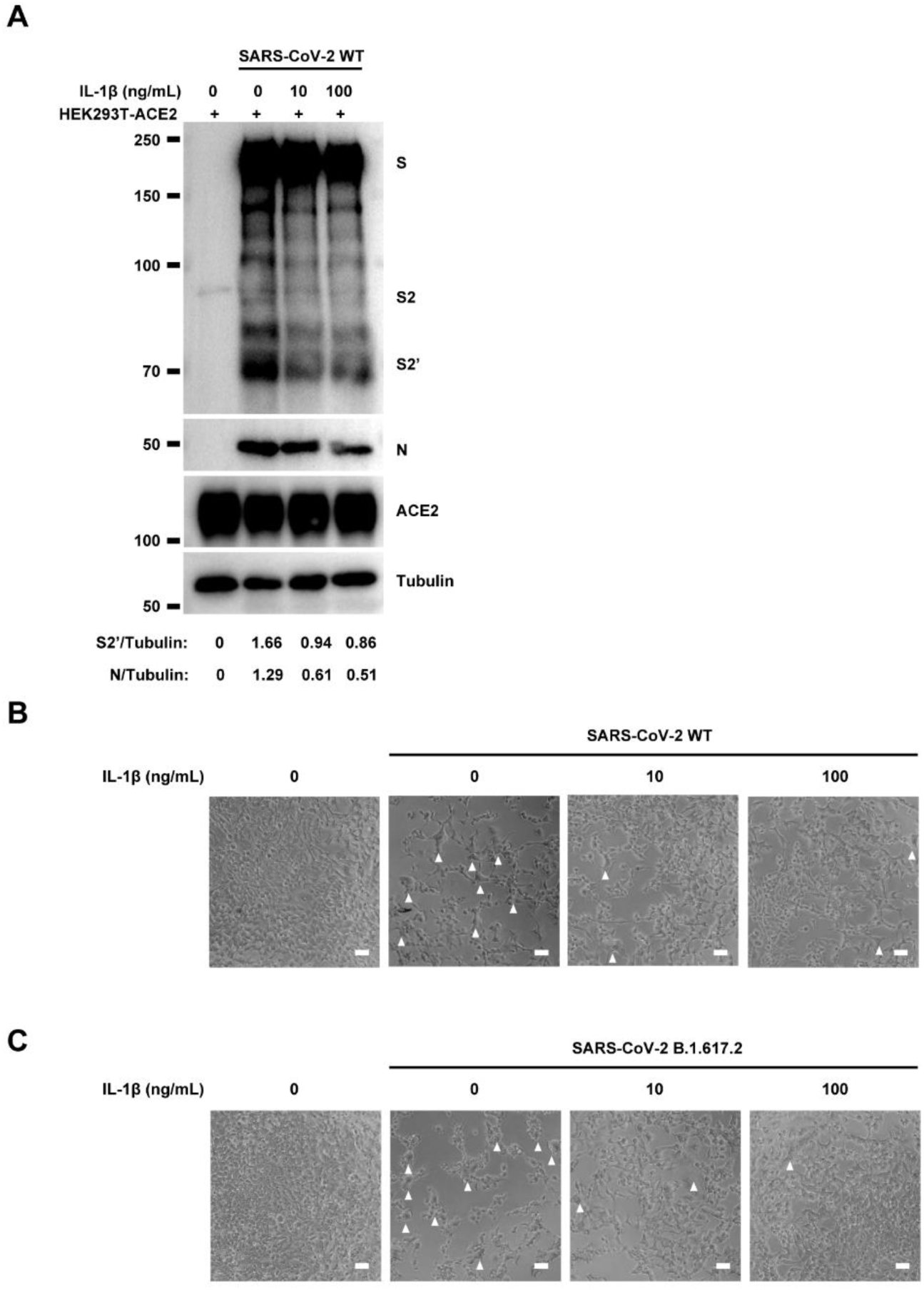
IL-1β inhibits authentic SARS-CoV-2-induced cell-cell fusion. (**A**) Immunoblots of WT SARS-CoV-2 S, S2, cleaved S2’ and N proteins collected from HEK293T-ACE2 cells 24 hpi as described in Figure 2E. Blots are representative of three individual experiments. Numbers below the blots indicated the intensity of S2’ or N versus Tubulin. (**B, C**) Bright field images of 0.5 MOI WT (**B**) or Delta (**C**) SARS-CoV-2 infected HEK293T-ACE2 cells pre-treated with different concentrations of IL-1β. White arrow heads indicate syncytia formation, scale bars are indicative of 50 μm, and images are representative of three independent experiments.

**Figure 2—figure supplement 4.**
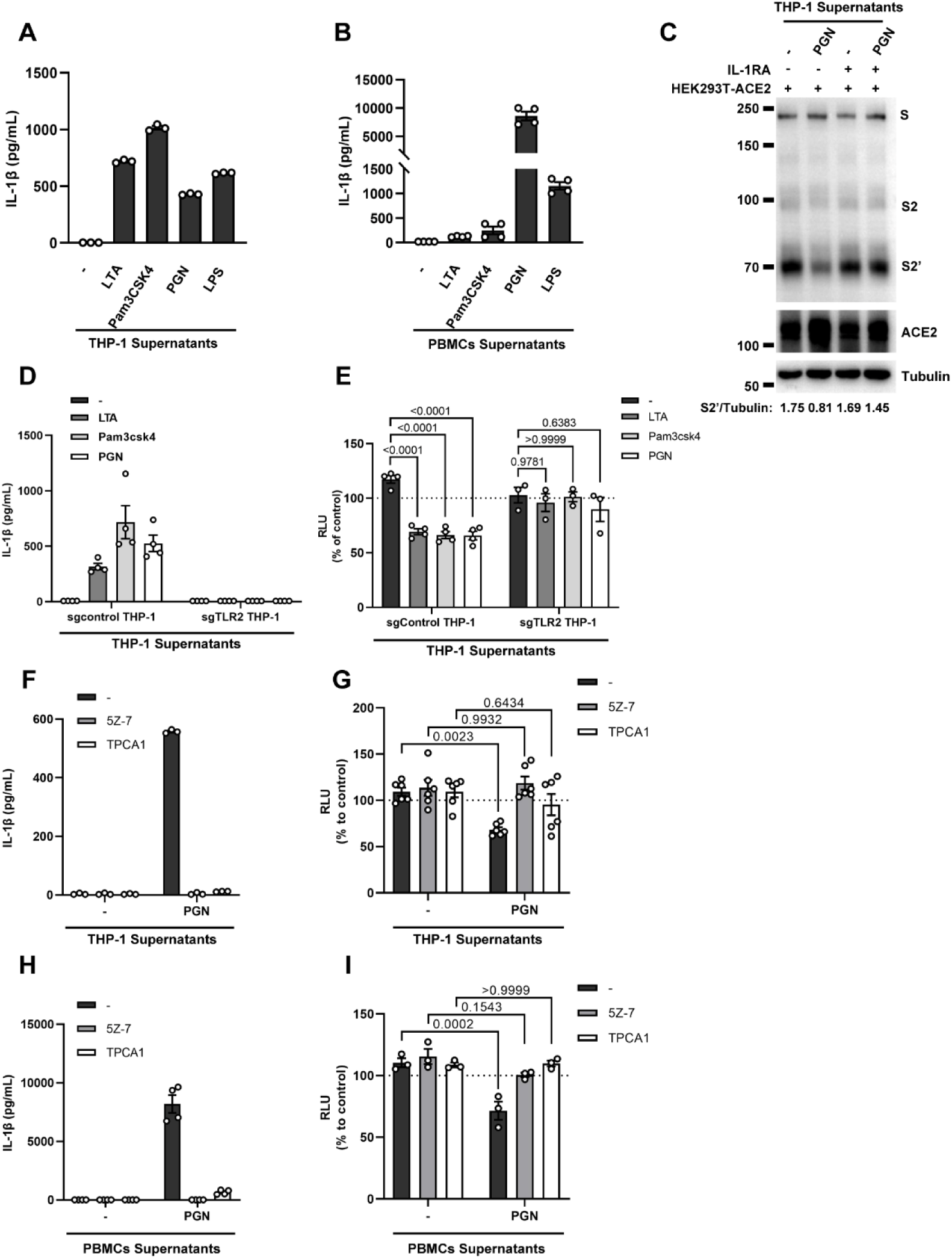
IL-1β is an important host factor from innate immune cells inhibiting SARS-CoV-2 spike-induced cell-cell fusion. (**A, B**) ELISA of IL-1β concentrations in supernatants from THP-1 (**A**) and PBMCs (**B**) after TLR ligands stimulation. Data are representative of three individual repeats. (**C**) immunoblots showing full-length spike, S2, cleaved S2’ and ACE2 collected from THP-1 supernatants-treated HEK293T-S and HEK293T-ACE2 in the presence or absence of IL-1RA. Blots are representative of three individual repeats. Numbers below the blots indicated the intensity of S2’ versus Tubulin. (**D**) ELISA of IL-1β concentrations in supernatants from sgControl and sgTLR2 THP-1 cells after TLR ligands stimulation. Data are representative of four individual repeats. (**E**) Luciferase activity (RLU) measured from HEK293T cell lysates collected from sgControl and sgTLR2 THP-1 supernatants-treated HEK293T-S and HEK293T-ACE2 for 16 hours. FBS free RPMI 1640 served as medium control. Data are representative of three individual repeats and displayed as individual points with mean ±SEM. (**F**) ELISA of IL-1β concentrations in supernatants from TAK1/IKKβ inhibitors-treated THP-1 cells after PGN stimulation. Data are representative of three individual repeats. (**G**) Luciferase activity (RLU) measured from HEK293T cell lysates collected from TAK1/IKKβ inhibitors pre-treated THP-1 supernatants added onto HEK293T-S and HEK293T-ACE2 cells for 16 hours. FBS free RPMI 1640 served as medium control. Data are representative of three individual repeats and displayed as individual points with mean ±SEM. (**H**) ELISA of IL-1β concentrations in supernatants from TAK1/IKKβ inhibitors-treated PBMCs after PGN stimulation. Data are representative of three individual repeats. (**I**) Luciferase activity (RLU) measured from HEK293T cell lysates collected from TAK1/IKKβ inhibitors pre-treated PBMCs supernatants added onto HEK293T-S and HEK293T-ACE2 cells for 16 hours. 1% FBS RPMI 1640 served as medium control. Data are representative of three individual repeats and displayed as individual points with mean ± SEM.

**Figure 2—figure supplement 5.**
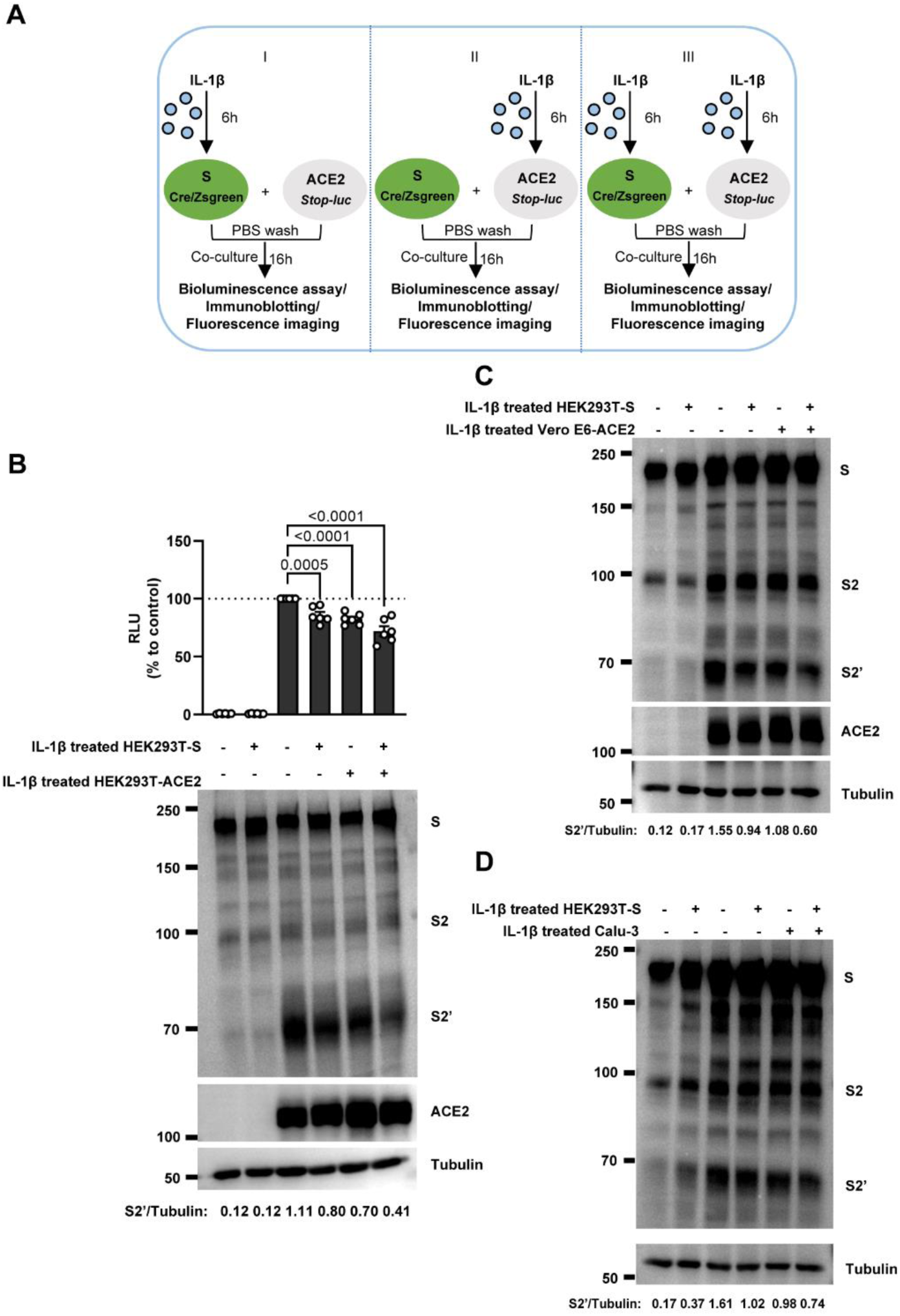
IL-1β inhibits SARS-CoV-2-induced cell-cell fusion through acting on both donor and acceptor cells. (**A**) Schematics of the cell-cell fusion model used to determine cell types affected by IL-1β. Pre-treated HEK293T-S or HEK293T-ACE2 cells or both with 1 ng/mL IL-1β for 6 hours, then co-cultured for 16 hours after washing with PBS. Cells co-expressing SARS-CoV-2 spike and Cre, were co-cultured with ACE2 and *Stop-luc* co-expressing HEK293T cells for 16 hours, before cell lysates were collected for bioluminescence assay and immunoblotting. Cells co-expressing SARS-CoV-2 spike and Zsgreen, were co-cultured with ACE2 expressing HEK293T cells for 16 hours before fluorescence imaging. (**B**) Luciferase activity (RLU) measured from HEK293T cell lysates and immunoblots showing full-length spike, S2, cleaved S2’ and ACE2 collected from different treatments of IL-1β described in (A) for 16 hours. Data and blots are representative of six individual repeats. Numbers below the blots indicated the intensity of S2’ versus Tubulin. (**C**) Immunoblots showing full-length spike, S2, cleaved S2’ and ACE2 collected from 1 ng/mL IL-1β-treated HEK293T-S and Vero E6-ACE2 for 16 hours. Blots are representative of three independent experiments. Numbers below the blots indicated the intensity of S2’ versus Tubulin. (**D**) Immunoblots showing full-length spike, S2 and cleaved S2’ collected from 1 ng/mL IL-1β-treated HEK293T-S and Calu-3 for 16 hours. Blots are representative of three independent experiments. Numbers below the blots indicated the intensity of S2’ versus Tubulin.

**Figure 2—figure supplement 6.**
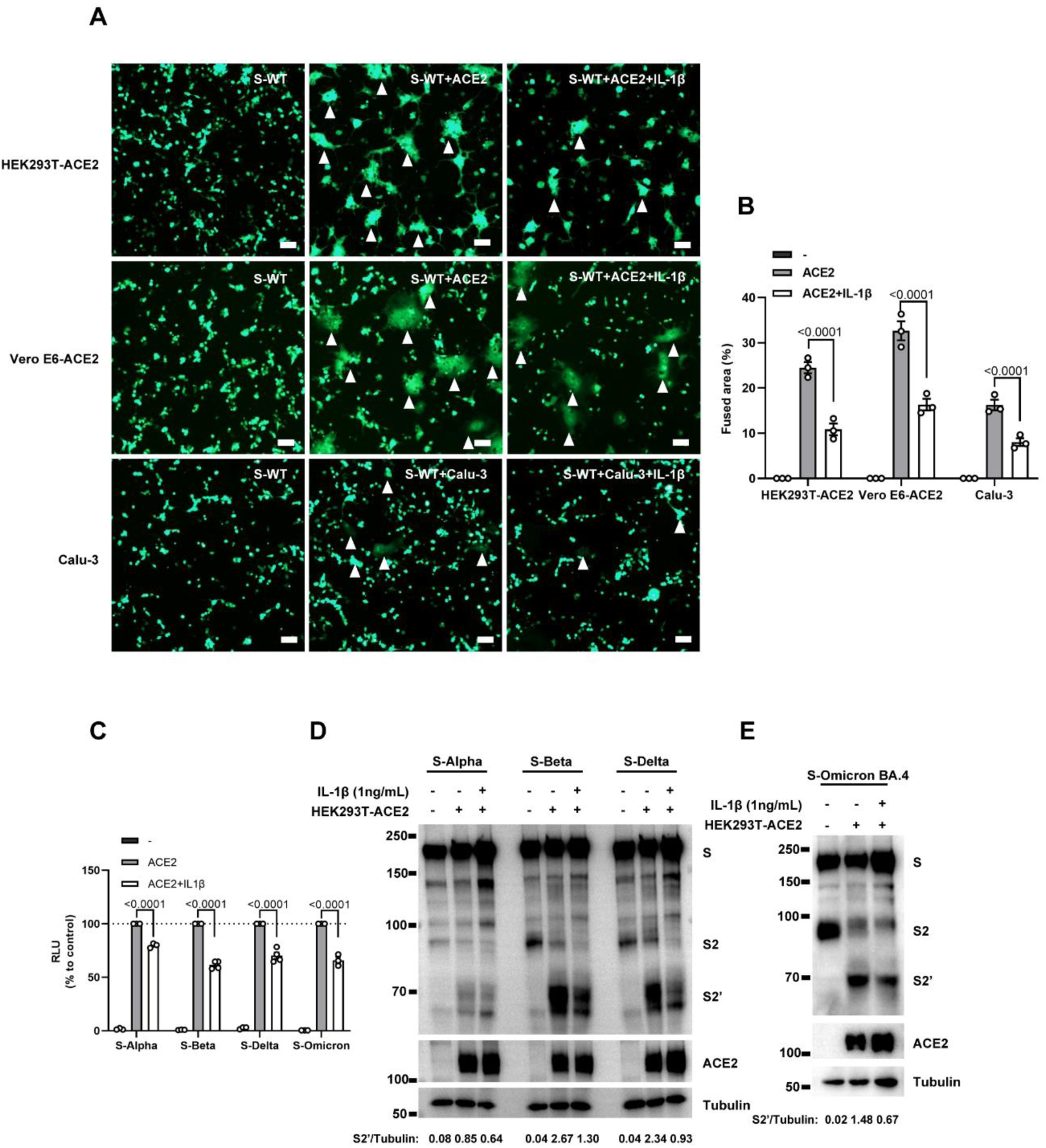
IL-1β inhibits SARS-CoV-2 spike-induced syncytia formation in different cells. (**A**) Representative fluorescent image captured at 488 nm from HEK293T-S-Zsgreen co-cultured with HEK293T-ACE2, Vero E6-ACE2 and Calu-3 for 16 hours with or without 1 ng/mL IL-1β. White arrow heads indicate syncytia formation. Scale bars, 50 μm. Images are representative of three individual repeats. (**B**) Quantification of the fused area in (A). (**C**) Luciferase activity (RLU) measured from HEK293T cell lysates from 1 ng/mL IL-1β-treated HEK293T-S (S-Alpha, S-Beta, S-Delta, S-Omicron BA.4) and HEK293T-ACE2 for 16 hours. Data are representative of four individual repeats and displayed as individual points with mean ±SEM. (**D**, **E**) Immunoblots showing full-length spike, S2, cleaved S2’ and ACE2 collected from 1 ng/mL IL-1β-treated HEK293T-S (S-Alpha, S-Beta, S-Delta (**D**), S-Omicron BA.4 (**E**)) and HEK293T-ACE2 for 16 hours. Blots are representative of three independent experiments. Numbers below the blots indicated the intensity of S2’ versus Tubulin.

**Figure 2—figure supplement 7.**
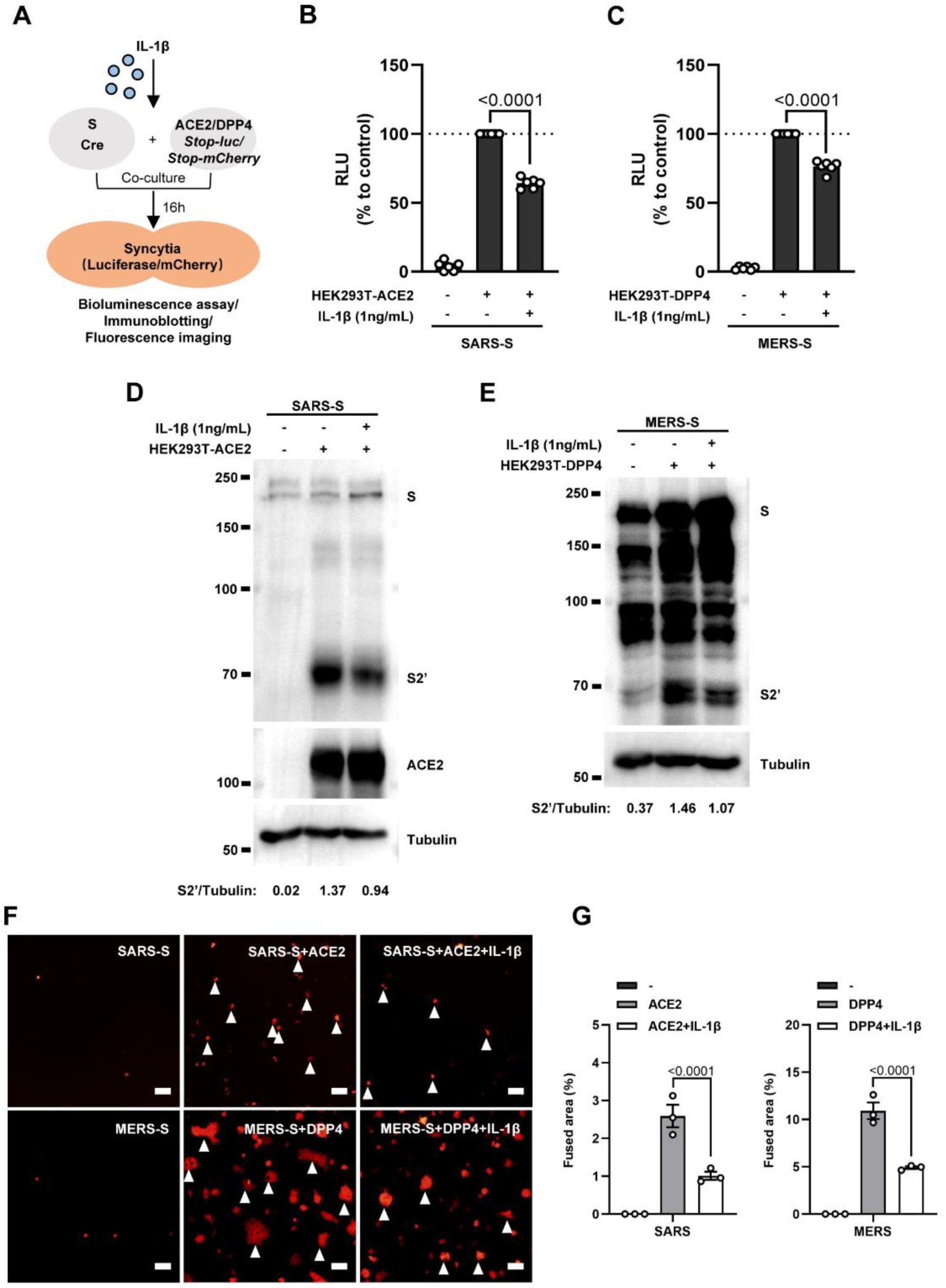
IL-1β inhibits SARS-CoV and MERS-CoV spike-induced cell-cell fusion. (**A**) Schematics of the cell-cell fusion model used to quantify SARS-CoV and MERS-CoV spike-mediated syncytia formation upon IL-1β treatment. Cells co-expressing SARS-CoV or MERS-CoV spike and Cre, were co-cultured with ACE2 or DPP4 and *Stop-luc* co-expressing HEK293T cells for 16 hours, before cell lysates were collected for bioluminescence assay and immunoblotting. Cells co-expressing SARS-CoV or MERS-CoV spike and Cre, were co-cultured with ACE2 or DPP4 and *Stop-mCherry* co-expressing HEK293T cells for 16 hours before fluorescence imaging. (**B**) Luciferase activity (RLU) measured from HEK293T cell lysates from 1 ng/mL IL-1β treated HEK293T-S (SARS-S) and HEK293T-ACE2 for 16 hours. Data are representative of four individual repeats and displayed as individual points with mean ±SEM. (**C**) Luciferase activity (RLU) measured from HEK293T cell lysates from 1 ng/mL IL-1β-treated HEK293T-S (MERS-S) and HEK293T-DPP4 for 16 hours. Data are representative of four individual repeats and displayed as individual points with mean ±SEM. (**D**) Immunoblots showing SARS-CoV full-length spike, cleaved S2’ and ACE2 collected from 1 ng/mL IL-1β-treated HEK293T-S (SARS-S) and HEK293T-ACE2 for 16 hours. Blots are representative of three independent experiments. Numbers below the blots indicated the intensity of S2’ versus Tubulin. (**E**) Immunoblots showing MERS-CoV full-length spike, cleaved S2’ collected from 1 ng/mL IL-1β-treated HEK293T-S (MERS-S) and HEK293T-DPP4 for 16 hours. Blots are representative of three independent experiments. Numbers below the blots indicated the intensity of S2’ versus Tubulin. (**F**) Representative fluorescent images captured at 594 nm from 1 ng/mL IL-1β-treated HEK293T-S (SARS-S) and HEK293T-ACE2 or HEK293T-S (MERS-S) and HEK293T-DPP4 for 16 hours. White arrow heads indicate syncytia formation. Scale bars, 50 μm. Images are representative of three individual experiments. (**G**) Quantification of the fused area in (F).

**Figure 3—figure supplement 1.**
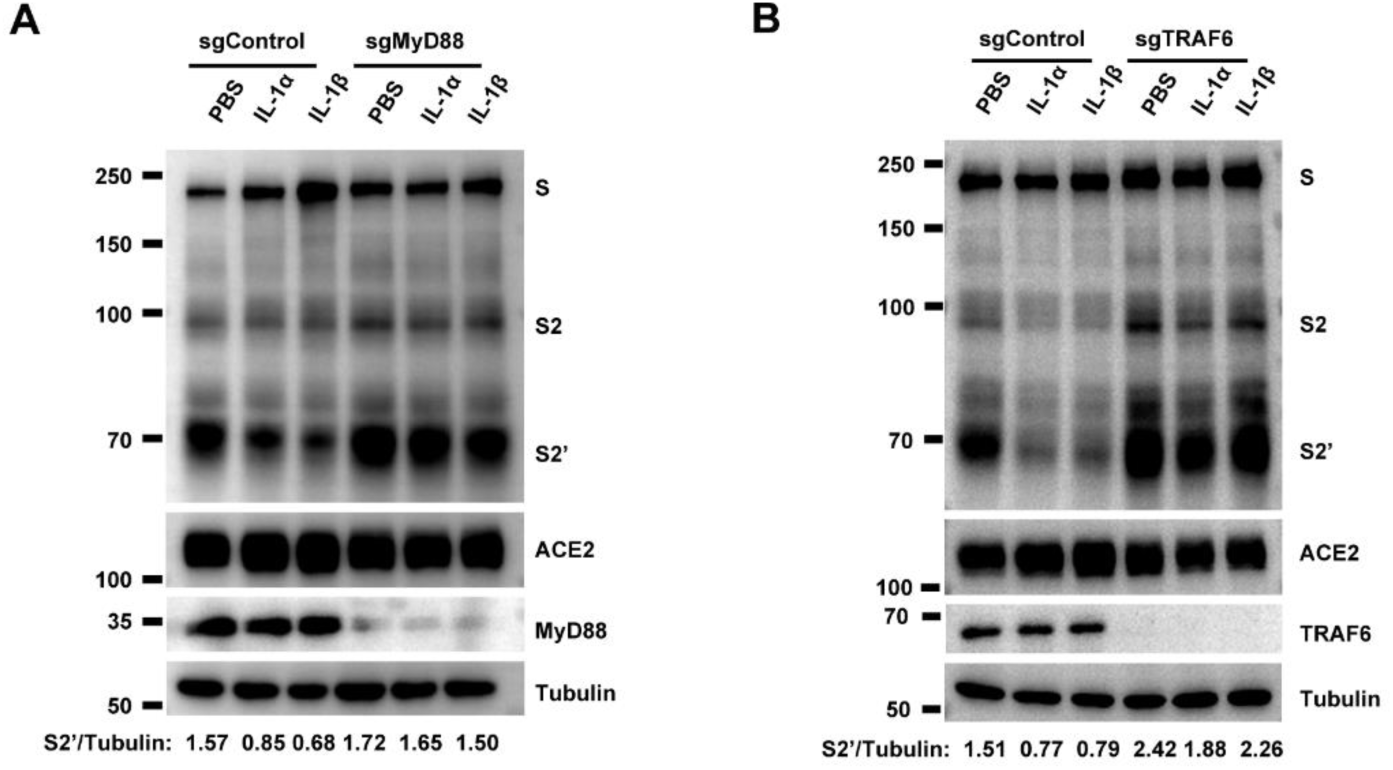
IL-1β inhibits SARS-CoV-2 spike-induced cell-cell fusion through the IL-1R1/MyD88/IRAK/TRAF6 pathway. (**A**) Immunoblots showing full-length spike, S2 and cleaved S2’, ACE2 and MyD88 collected from 10 ng/mL IL-1α or 1 ng/mL IL-1β treated sgControl or sgMyD88 HEK293T cell-cell fusion system for 16 hours. Blots are representative of three independent experiments. Numbers below the blots indicated the intensity of S2’ versus Tubulin. (**B**) Immunoblots showing full-length spike, S2 and cleaved S2’, ACE2 and TRAF6 collected from 10 ng/mL IL-1α or 1 ng/mL IL-1β treated sgControl or sgTRAF6 HEK293T cell-cell fusion system for 16 hours. Blots are representative of three independent experiments. Numbers below the blots indicated the intensity of S2’ versus Tubulin.

**Figure 3—figure supplement 2.**
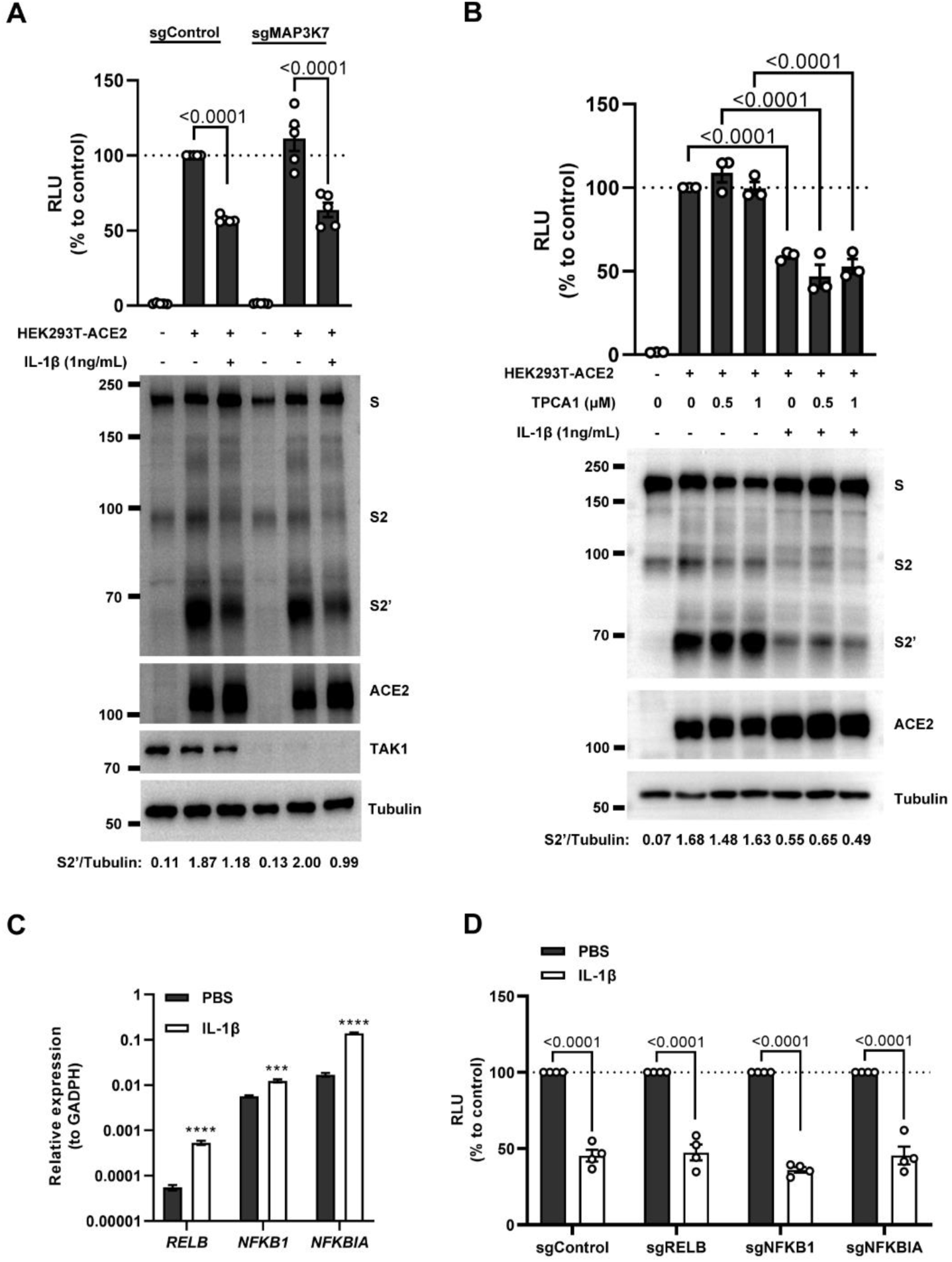
IL-1β inhibits SARS-CoV-2 spike-induced cell-cell fusion independent from the TAK1/IKKβ/NF-κB pathway. (**A**) Luciferase activity (RLU) measured from cell lysates and immunoblots showing full-length spike, S2 and cleaved S2’, ACE2 collected from 1 ng/mL IL-1β treated sgControl or sgTAK1 HEK293T cell-cell fusion system for 16 hours. Data and blots are representative of five individual repeats. Numbers below the blots indicated the intensity of S2’ versus Tubulin. (**B**) Luciferase activity (RLU) measured from HEK293T cell lysates and immunoblots showing full-length spike, S2 and cleaved S2’, ACE2 collected from HEK293T-S and HEK293T-ACE2 pre-treated with different concentrations of TPCA1 for 30 min, then treated with 1 ng/mL IL-1β for 16 hours. Data and blots are representative of three individual repeats. Numbers below the blots indicated the intensity of S2’ versus Tubulin. (**C**) mRNA levels of NF-κB pathway-related genes in HEK293T cells after 1 ng/mL IL-1β for 4 hours. Data are representative of three individual repeats. (**D**) Luciferase activity (RLU) measured from cell lysates collected from 1 ng/mL IL-1β-treated sgControl, sgRELB, sgNFKB1 or sgNFKBIA HEK293T cell-cell fusion system for 16 hours. Data are representative of four individual repeats and displayed as individual points with mean ±SEM.

**Figure 4—figure supplement 1.**
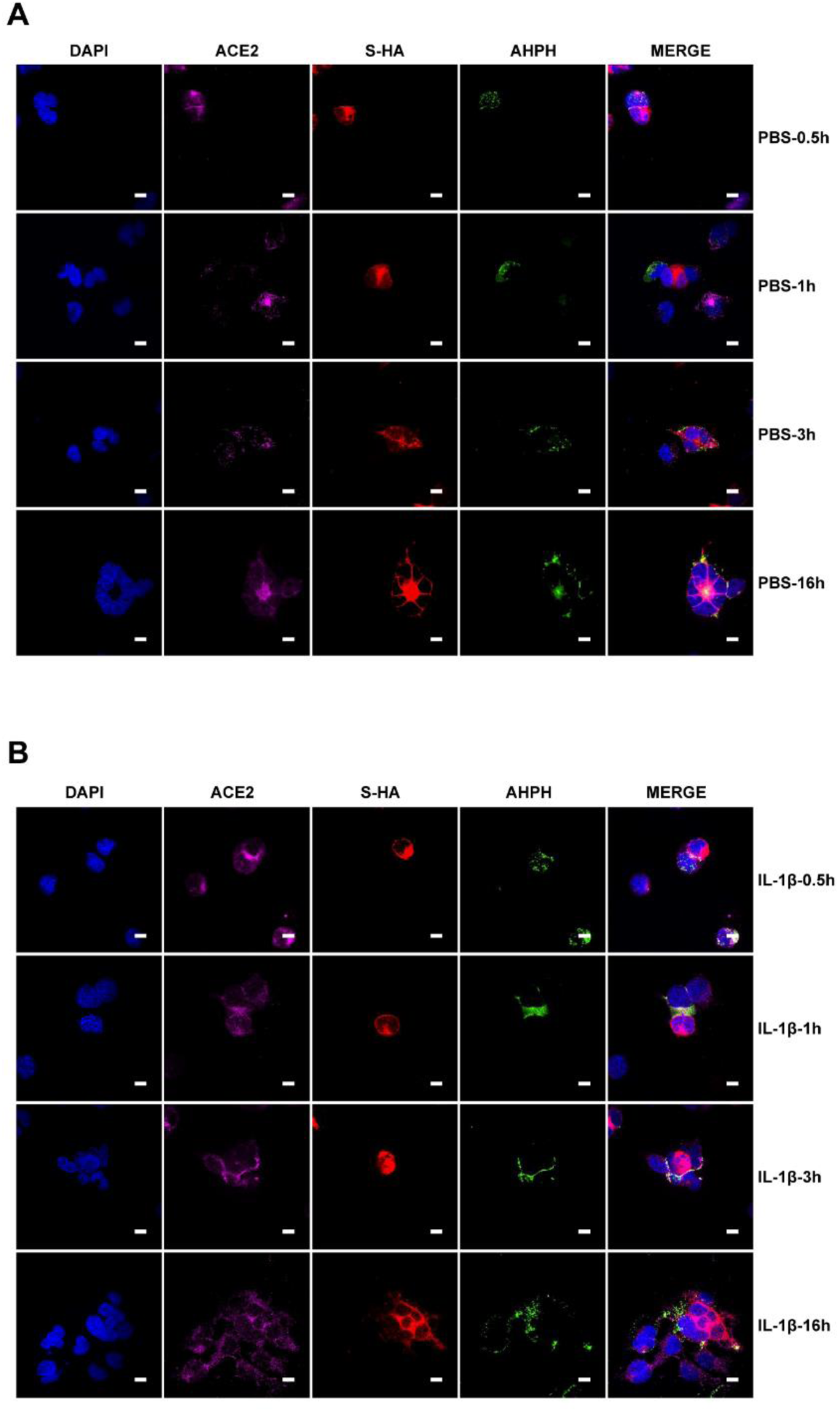
Single-channel confocal images. (**A**) Single-channel confocal images of Figure 4D top panel. (**B**) Single-channel confocal images of Figure 4D bottom panel.

**Figure 4—figure supplement 2.**
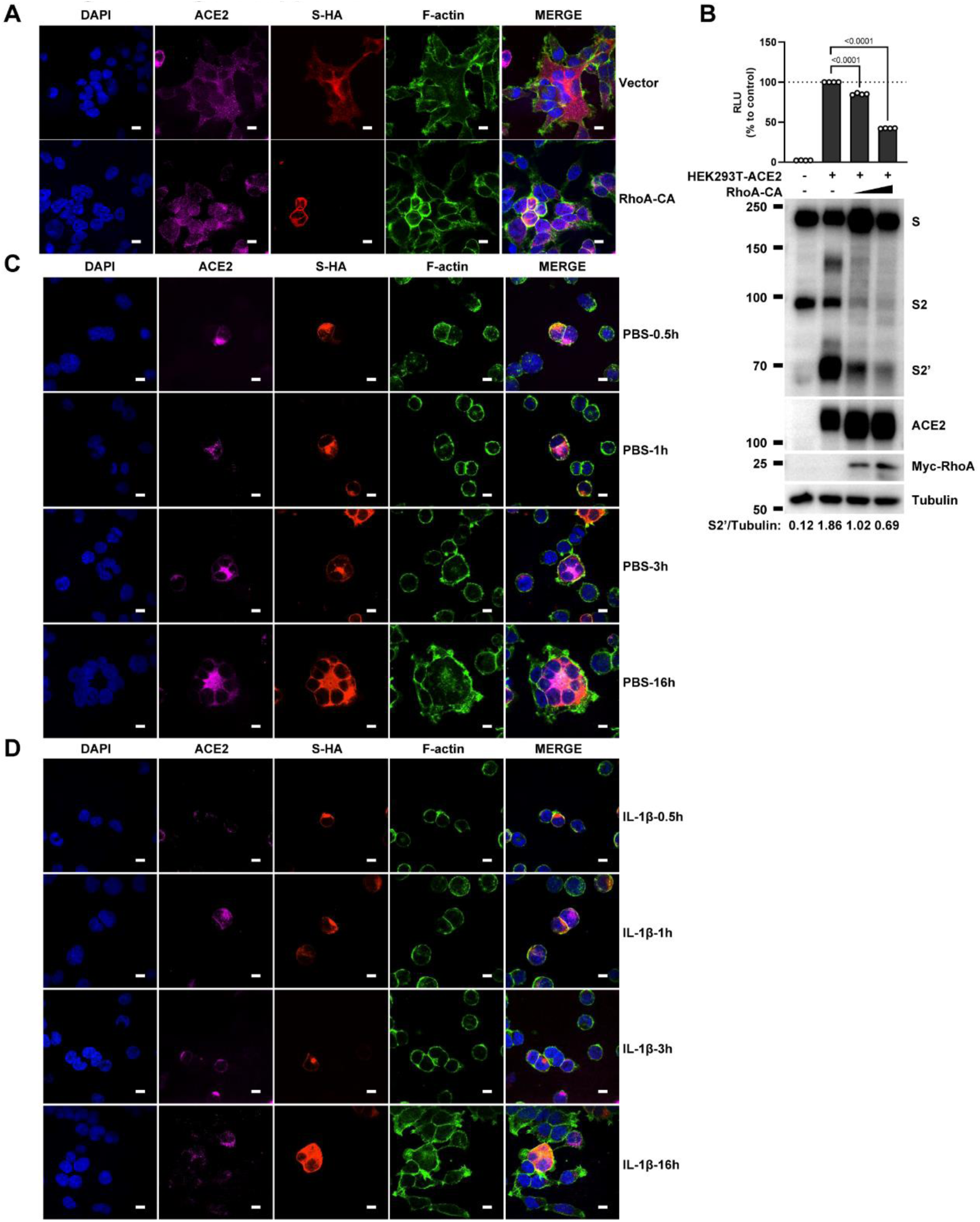
Single-channel confocal images. (**A**) Single-channel confocal images of Figure 4E. (**B**) Luciferase activity (RLU) measured from HEK293T cell lysates and immunoblots showing full-length spike, S2, cleaved S2’, ACE2 and Myc-RhoA collected from transfected vector, 10 ng or 20 ng constitutively active RhoA mutant (RhoA-CA) both in HEK293T-S and HEK293T-ACE2 cells. Data and blots are representative of four individual repeats. Numbers below the blots indicated the intensity of S2’ versus Tubulin. (**C**) Single-channel confocal images of Figure 4F top panel. (**D**) Single-channel confocal images of Figure 4F middle panel.

**Figure 4—figure supplement 3.**
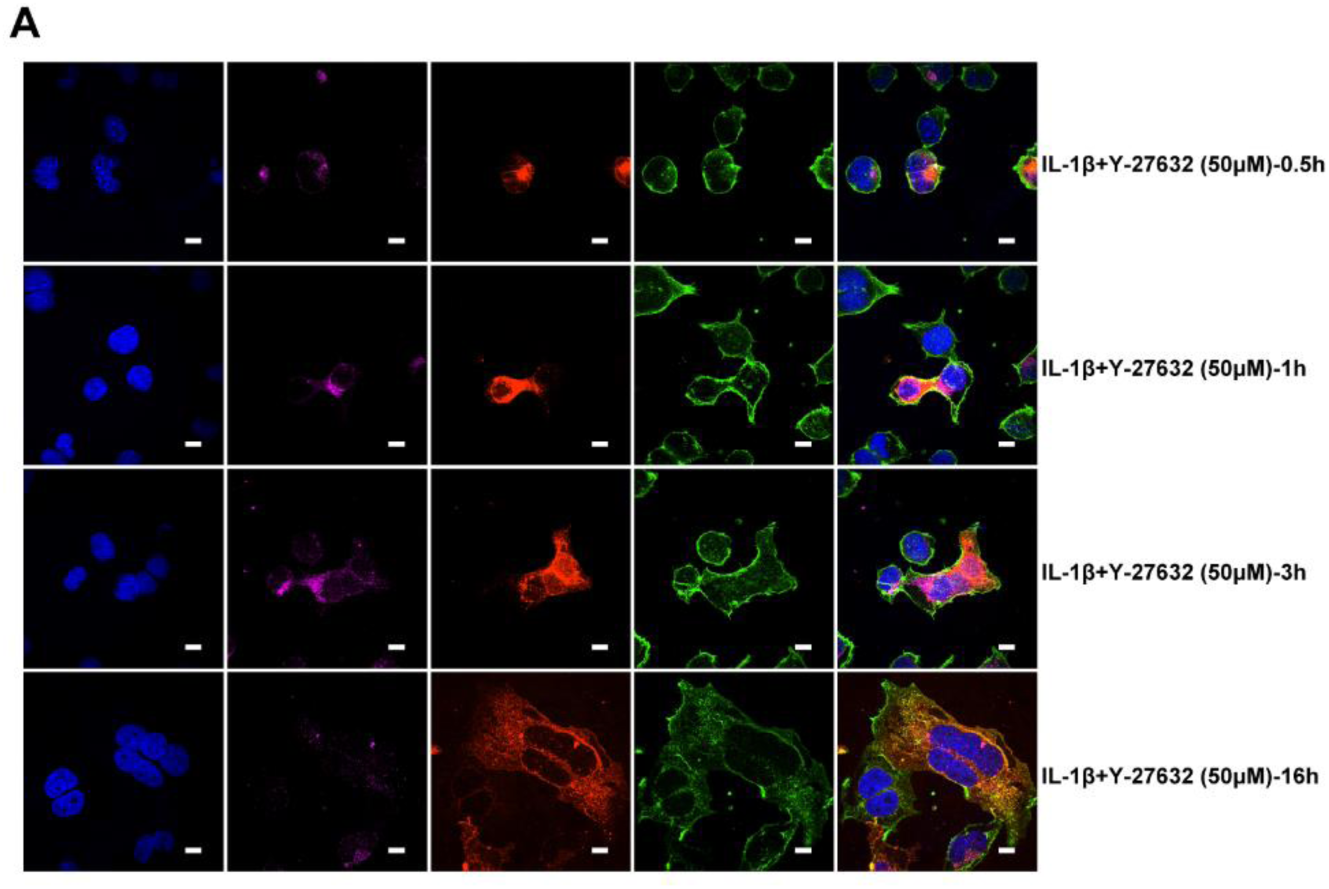
Single-channel confocal images. **(A)** Single-channel confocal images of Figure 4F bottom panel.

**Figure 5—figure supplement 1.**
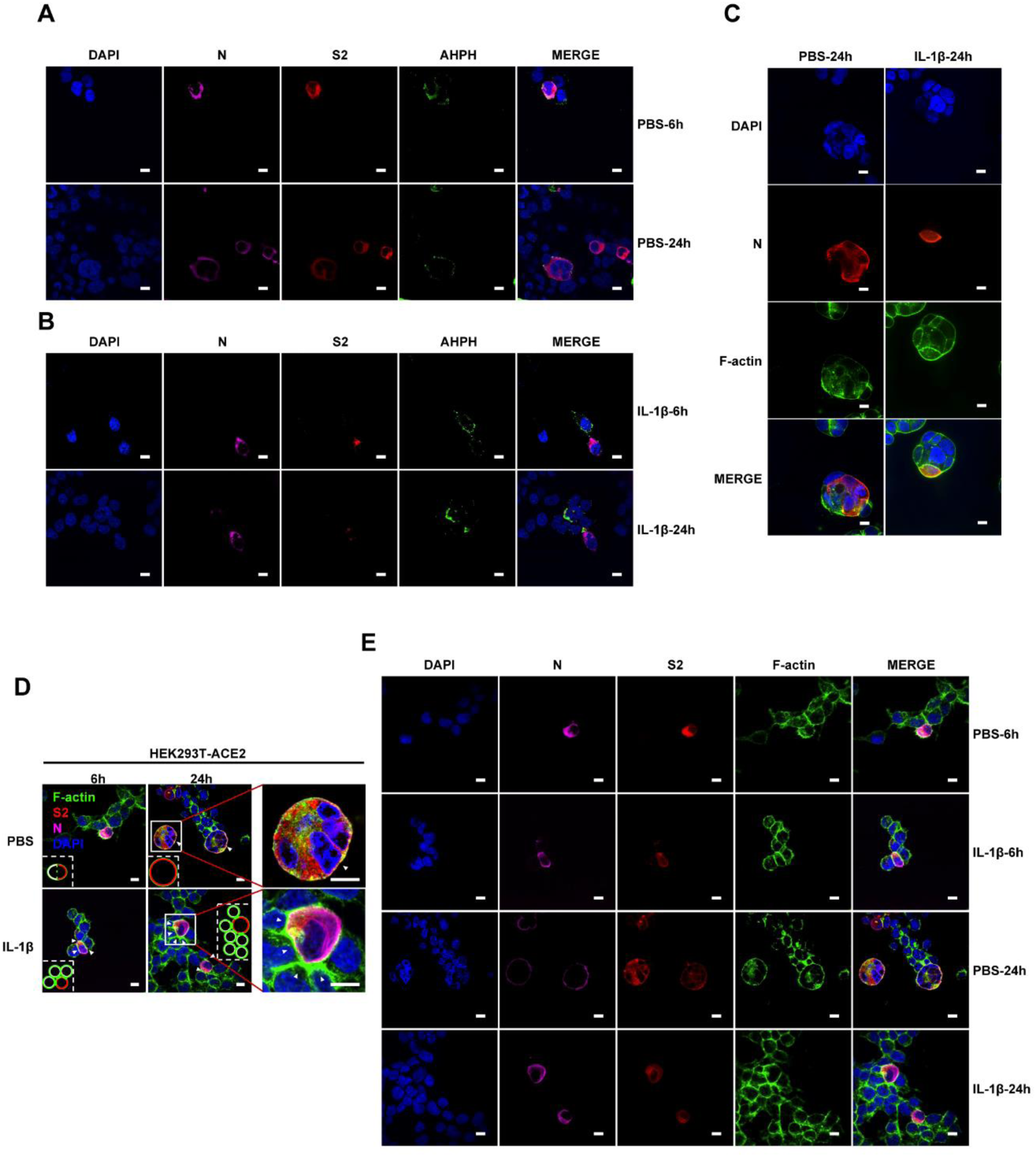
Single-channel confocal images. (**A**) Single-channel confocal images of Figure 5A top panel. (**B**) Single-channel confocal images of Figure 5A bottom panel. (**C**) Single-channel confocal images of Figure 5B. (**D**) Representative confocal images of F-actin stained with phalloidin-488 in the presence or absence of 1 ng/mL IL-1β treatment in 0.5 MOI WT authentic SARS-CoV-2 infected HEK293T-ACE2 cells at 6 or 24 hpi. Schematics with green lines in the white dashed line boxes representing actin bundles, red cycles representing SARS-CoV-2 infected cells, and white cycles representing neighboring cells. White arrow heads indicate the enrichment or disappearance of F-actin, scale bars, 10 μm. Images are representative of four independent experiments. (**E**) Single-channel confocal images of (D).

**Figure 5—figure supplement 2.**
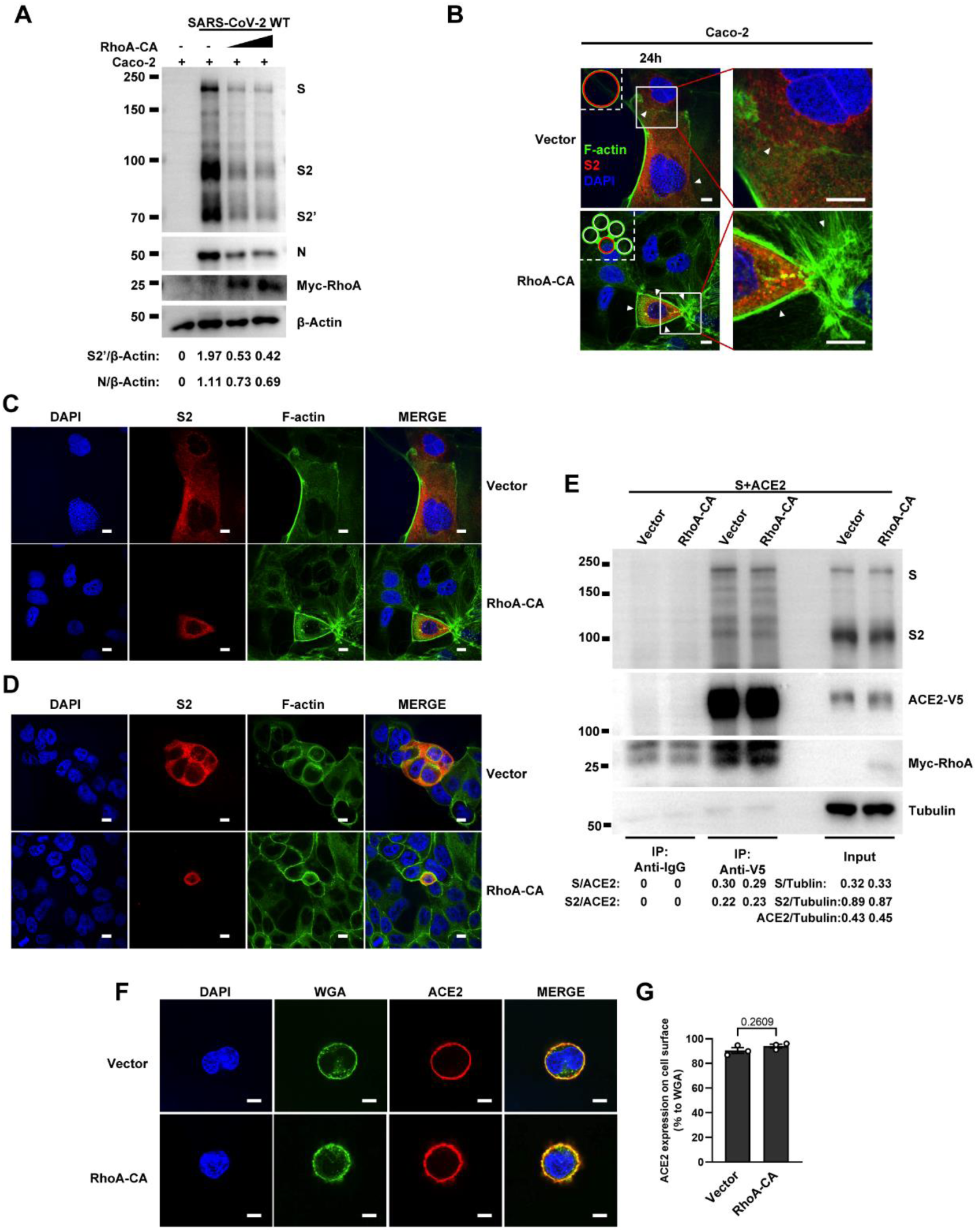
RhoA-CA does not affect Spike protein binding to ACE2 or ACE2 distribution on the cell surface. (**A**) Immunoblots of WT SARS-CoV-2 S, S2, cleaved S2’, N and Myc-RhoA collected from Caco-2 cells, which were transfected with vector, 10 ng or 20 ng RhoA-CA before infection with 0.5 MOI WT authentic SARS-CoV-2 for 24 hours. Blots are representative of three individual experiments. Numbers below the blots indicated the intensity of S2’ or N versus β-Actin. (**B**) Representative confocal images of F-actin stained with phalloidin-488 from Caco-2 cells described in (A). Schematics with green lines in the white dashed line boxes representing actin bundles, red cycles representing S-expressing cells. White arrow heads indicate the enrichment or disappearance of F-actin, scale bars, 10 μm. Images are representative of four independent experiments. (**C**) Single-channel confocal images of (B). (**D**) Single-channel confocal images of Figure 5E. (**E**) Co-immunoprecipitation (IP) and input controls of full-length spike protein after anti-V5 or anti-IgG pulldown from cell lysates mixed between HEK293T cells expressing ACE2-V5-6his or Spike protein with vector or RhoA-CA (10 ng). Blots are representative of three individual experiments. Numbers below the blots indicated the intensity of S or S2 versus ACE2 in IP group and S, S2 or ACE2 versus Tubulin in input group. (**F**) Representative confocal images of Wheat Germ Agglutinin (WGA, cell surface marker) and ACE2 from HEK293T cells co-transfected with ACE2 and vector or RhoA-CA (10 ng). Scale bars, 10 μm. Images are representative of three independent experiments. (**G**) Quantification of the relative ACE2 expression on the cell surface in (F).

**Figure 5—figure supplement 3.**
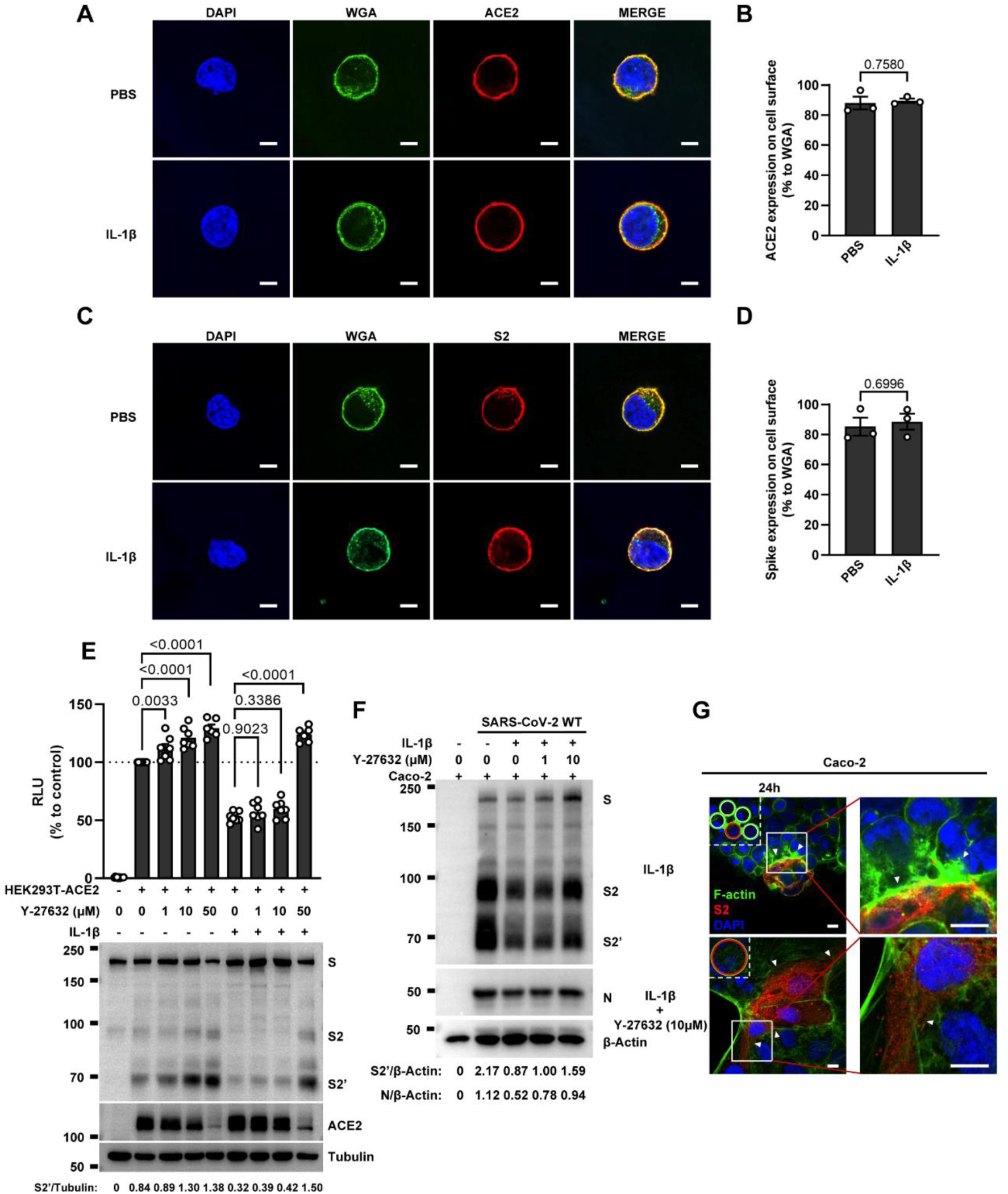
IL-1β does not affect ACE2 and Spike distribution on the cell surface. (**A**) Representative confocal images of Wheat Germ Agglutinin (WGA) and ACE2 from HEK293T cells transfected with ACE2 and treated with PBS or IL-1β. Scale bars, 10 μm. Images are representative of three independent experiments. (**B**) Quantification of the relative ACE2 expression on the cell surface in (A). (**C**) Representative confocal images of WGA and Spike from HEK293T cells transfected with Spike and treated with PBS or IL-1β. Scale bars, 10 μm. Images are representative of three independent experiments. (**D**) Quantification of the relative Spike expression on the cell surface in (C). (**E**) Luciferase activity (RLU) measured from HEK293T cell lysates and immunoblots showing full-length spike, S2 and cleaved S2’, ACE2 collected from HEK293T-S and HEK293T-ACE2 pre-treated with different concentrations of Y-27632 for 30 min, then treated with 1 ng/mL IL-1β for 16 hours. Data and blots are representative of three independent experiments. Numbers below the blots indicated the intensity of S2’ versus Tubulin. (**F**) Immunoblots of WT SARS-CoV-2 S, S2, cleaved S2’ and N collected from Caco-2 cells, which were treated with different concentrations of Y-27632 and 10 ng/mL IL-1β for 1 hour, then infected with 0.5 MOI WT authentic SARS-CoV-2 for 24 hours. Blots are representative of three individual experiments. Numbers below the blots indicated the intensity of S2’ or N versus β-Actin. (**G**) Representative confocal images of F-actin stained with phalloidin-488 in Caco-2 cells described in (F). Schematics with green lines in the white dashed line boxes representing actin bundles, red cycles representing S-expressing cells. White arrow heads indicate the enrichment or disappearance of F-actin, scale bars, 10 μm. Images are representative of four independent experiments.

**Figure 5—figure supplement 4.**
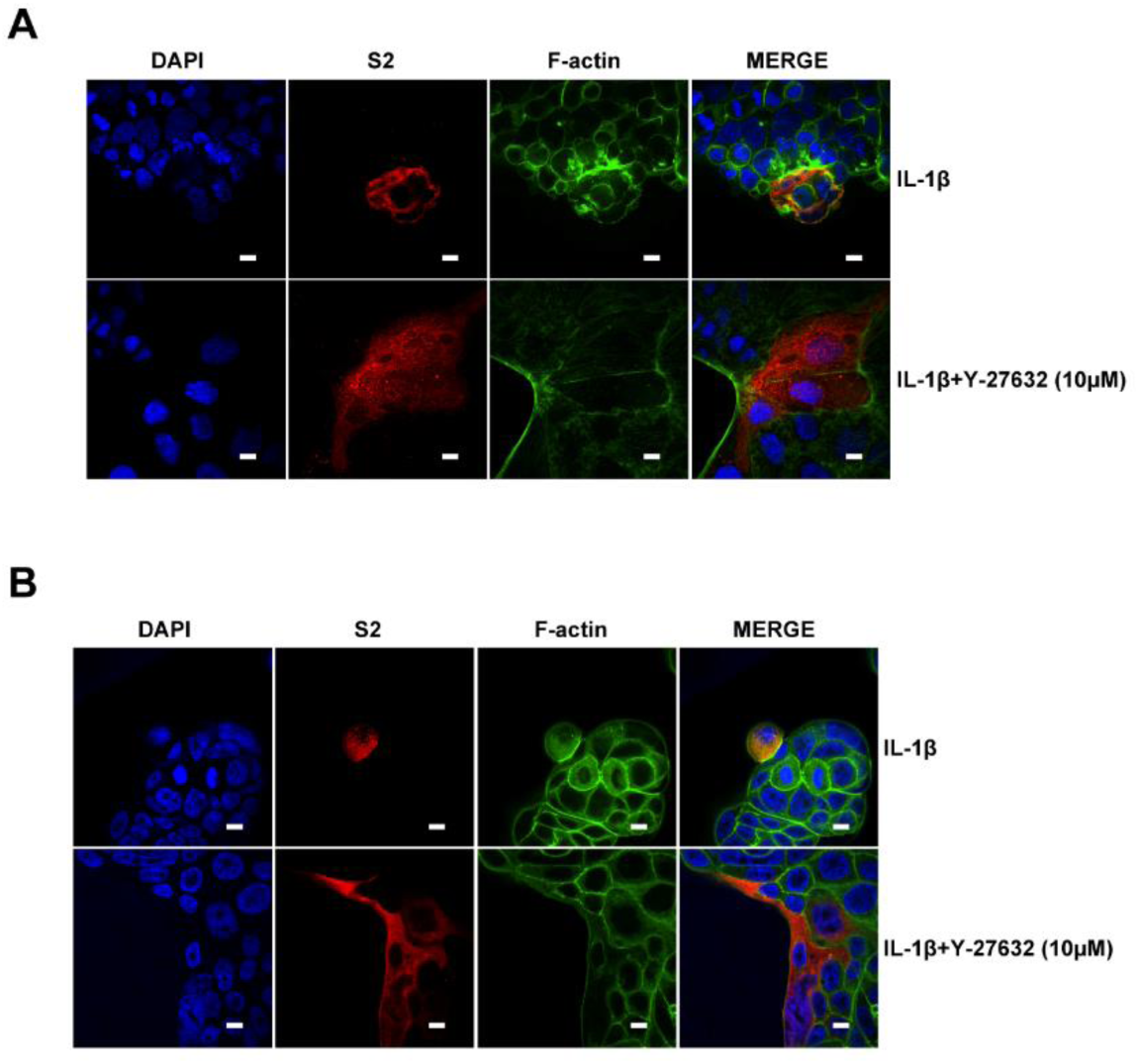
Single-channel confocal images. (**A**) Single-channel confocal images of Figure 5—figure supplement 3G. (**B**) Single-channel confocal images of Figure 5H.

**Figure 6—figure supplement 1.**
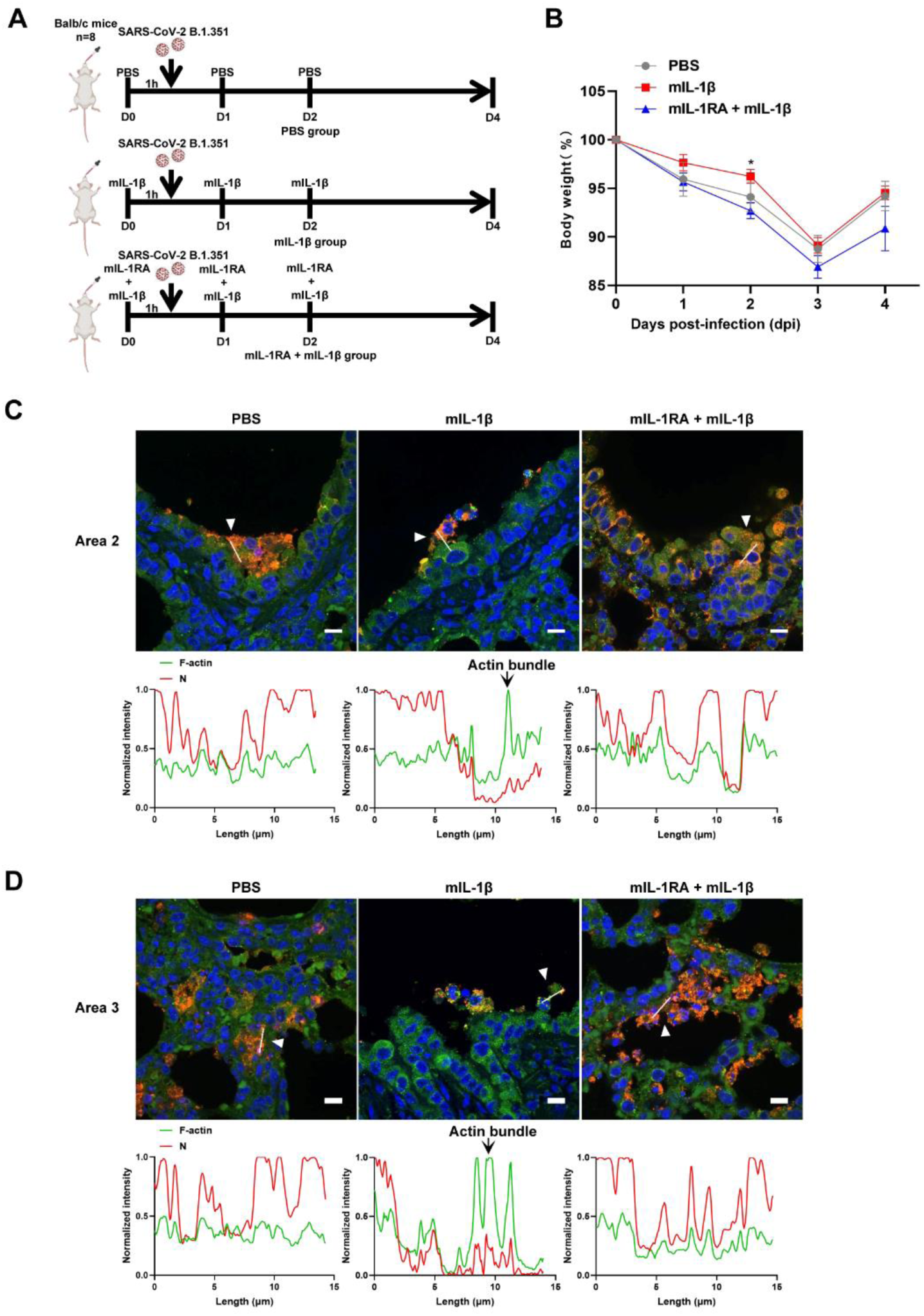
IL-1β reduces the weight loss and viral transmission upon SARS-CoV-2 infection *in vivo*. (**A**) Schematic for a murine model of authentic SARS-CoV-2 infection, PBS control (n=8); 1 μg/kg mIL-1β (n=8); 150 μg/kg mIL-1RA + 1 μg/kg mIL-1β (n=8) were administered 1 hour before intranasal challenge with 5 × 10^4^ FFU of SARS-CoV-2 B.1.351; mice were then intraperitoneal injection with PBS, mIL-1β and mIL-1RA + mIL-1β at 1 dpi and 2 dpi, before sacrificed at 4 dpi. (**B**) The body weights were assessed daily for weight loss after SARS-CoV-2 infection. (**C**, **D**) Representative confocal images of F-actin stained with phalloidin-488 and SARS-CoV-2 N in the area 2 and area 3 of lung tissue at 4 dpi. White arrow heads indicate syncytia formation or infected cells, scale bars are indicative of 10 μm and images are representative of three samples (Top). White lines indicate SARS-CoV-2 cell-cell transmission and quantify with fluorescence intensity of F-actin and SARS-CoV-2 N (Bottom).

**Figure 6—figure supplement 2.**
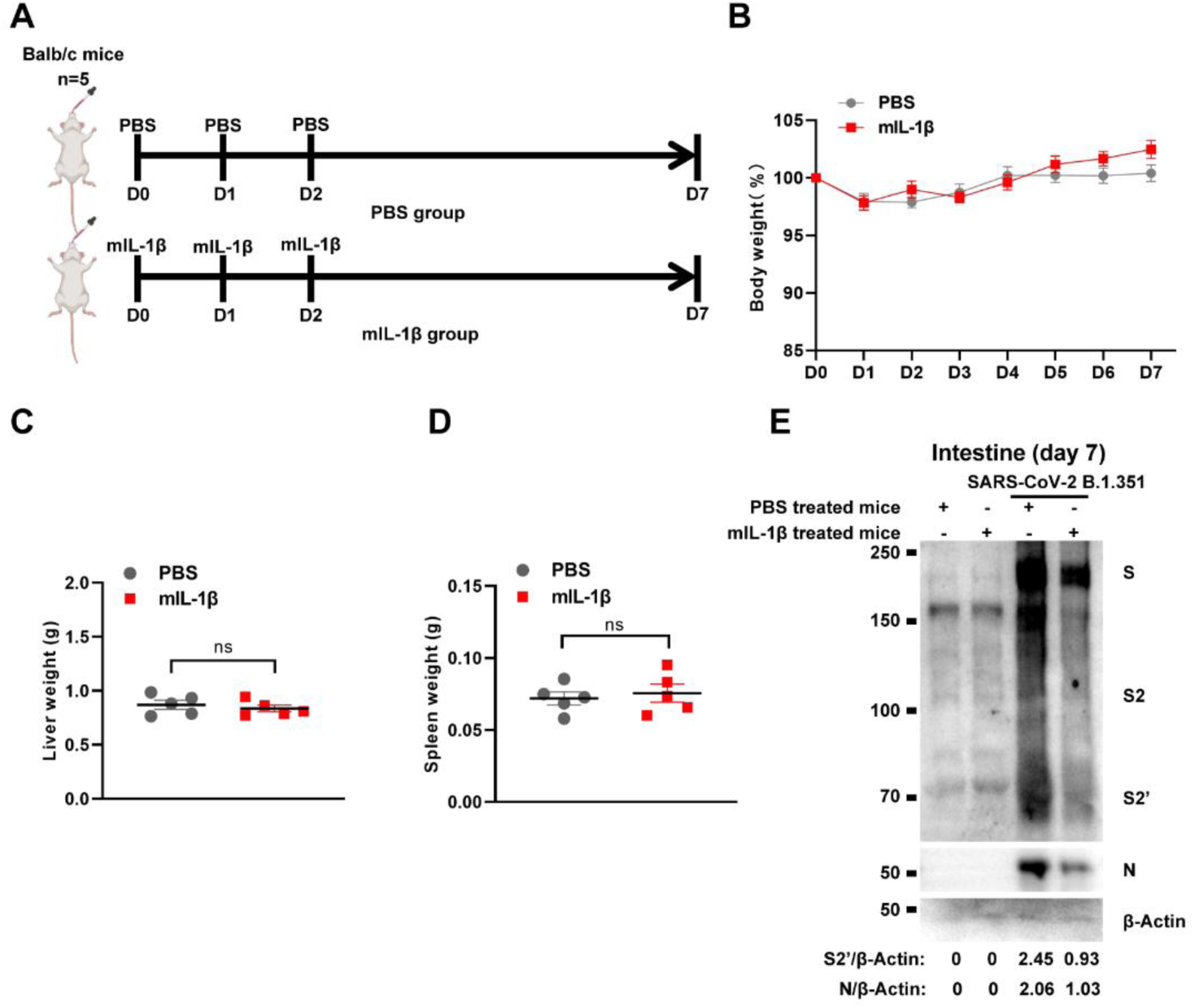
Treatment with 1 μg/kg mIL-1β protects mouse lung and intestinal tissue cells against SARS-CoV-2 infection without toxicity *in vivo*. (**A**) Schematic of PBS (n=5) and mIL-1β (n=5) treated BALB/c mice. (**B**) The body weights were assessed daily for weight loss. Liver (**C**) and spleen (**D**) weights were assessed at day 7. (**E**) Immunoblots of SARS-CoV-2 S, S2, cleaved S2’ and N proteins collected from SARS-CoV-2 B.1.351 infected intestine tissue cells, which isolated from BALB/c mice treated with or without 1 μg/kg mIL-1β at day 7. Blots are representative of three individual mouse. Numbers below the blots indicated the intensity of S2’ or N versus β-Actin.

**Figure 7—figure supplement 1.**
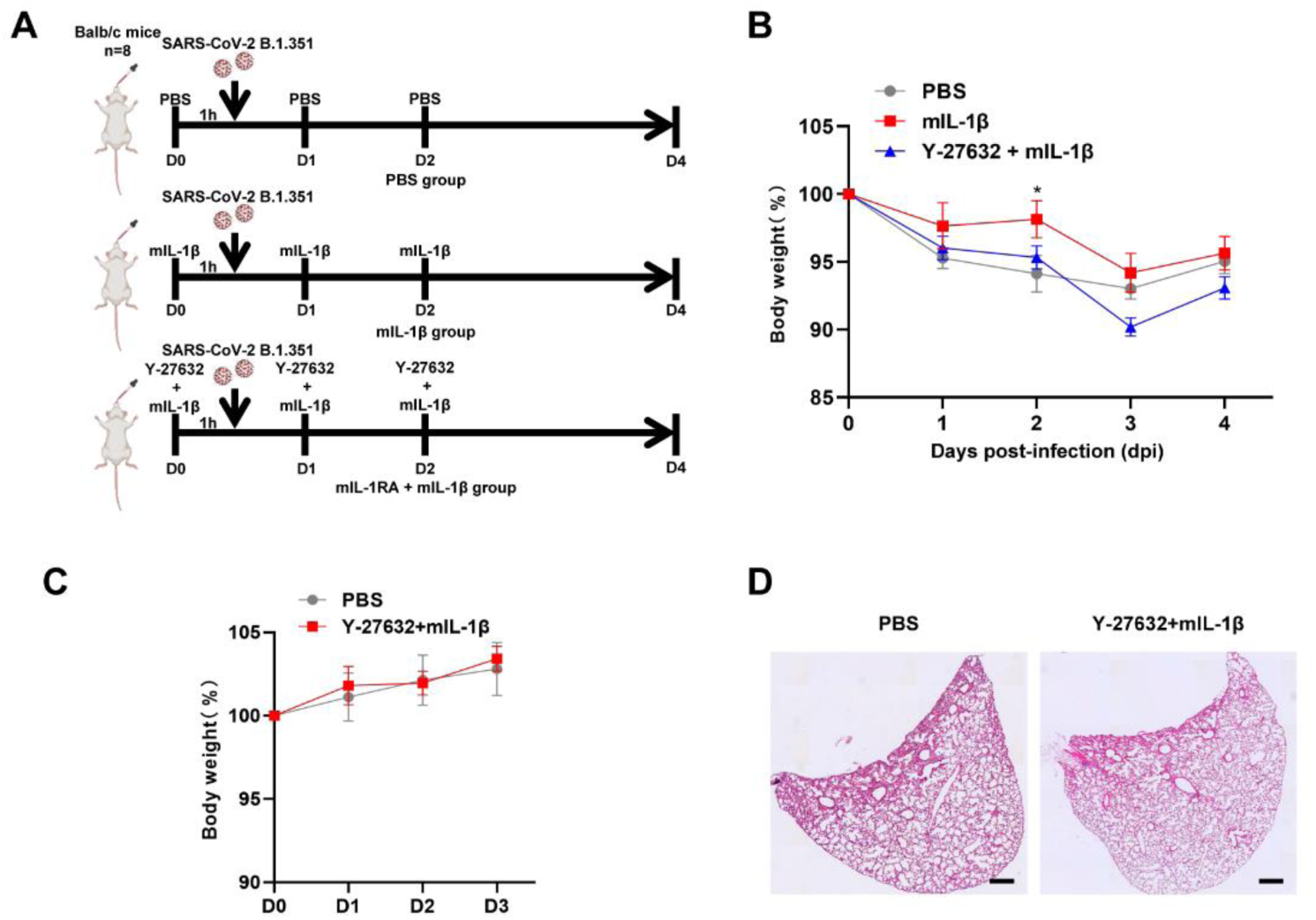
Y-27632+IL-1β do not cause weight loss and lung injury in uninfected BALB/c mice. (**A**) Schematic for a murine model of authentic SARS-CoV-2 infection, PBS control (n=8); 1 μg/kg mIL-1β (n=8); 1mg/kg Y-27632 + 1 μg/kg mIL-1β (n=8) were administered 1 hour before intranasal challenge with 5 × 10^4^ FFU of SARS-CoV-2 B.1.351; mice were then intraperitoneal injection with PBS, mIL-1β and Y-27632 + mIL-1β at 1 dpi and 2 dpi, before sacrificed at 4 dpi. (**B**) The body weights were assessed daily for weight loss after SARS-CoV-2 infection. (**C**) The body weights were assessed daily from PBS or Y-27632 + mIL-1β treated uninfected mice. (**D**) Representative images of H&E-stained lung sections from PBS or Y-27632 + mIL-1β-treated uninfected mice at day7, scale bars are indicative of 500 μm and images are representative of four samples.

**Figure 7—figure supplement 2.**
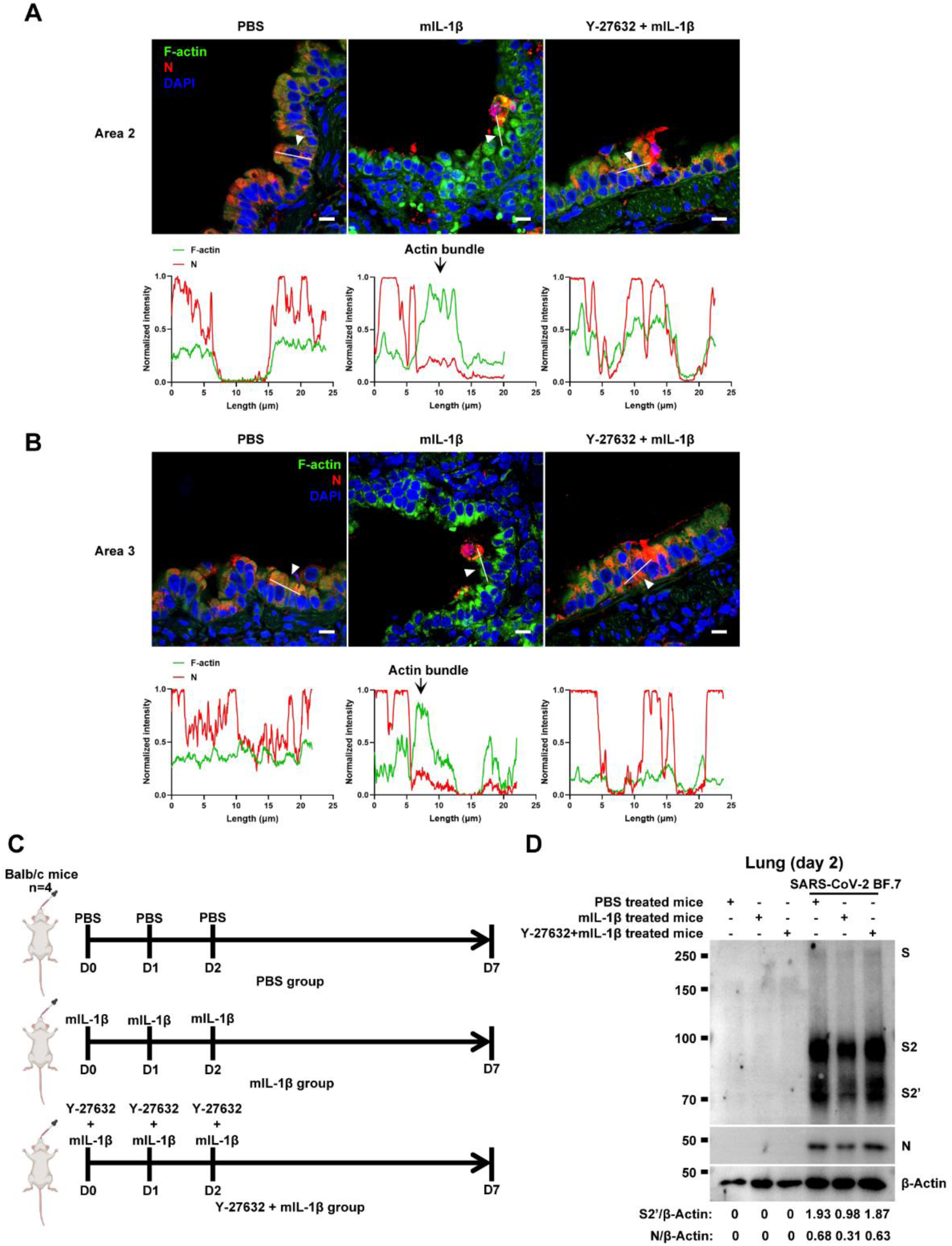
Y-27632 treatment prevents the formation of IL-1β-induced actin bundles at cell-cell junctions *in vivo*. (**A**, **B**) Representative confocal images of F-actin stained with phalloidin-488 and SARS-CoV-2 N in the area 2 and area 3 of lung tissue at 4 dpi. White arrow heads indicate syncytia formation or infected cells, scale bars are indicative of 10 μm and images are representative of three samples (Top). White lines indicate SARS-CoV-2 cell-cell transmission and quantify with fluorescence intensity of F-actin and SARS-CoV-2 N (Bottom). (**C**) Schematic of PBS, mIL-1β and Y-27632 + mIL-1β treated BALB/c mice. (**D**) Immunoblots of SARS-CoV-2 S, S2, cleaved S2’ and N proteins collected from authentic SARS-CoV-2 BF.7 infected lung tissue cells, which were isolated from BALB/c mice treated with PBS, 1 μg/kg mIL-1β or 1mg/kg Y-27632 + 1 μg/kg mIL-1β at day 2. Blots are representative of three individual mouse. Numbers below the blots indicated the intensity of S2’ or N versus β-Actin.

**Figure 7—figure supplement 3.**
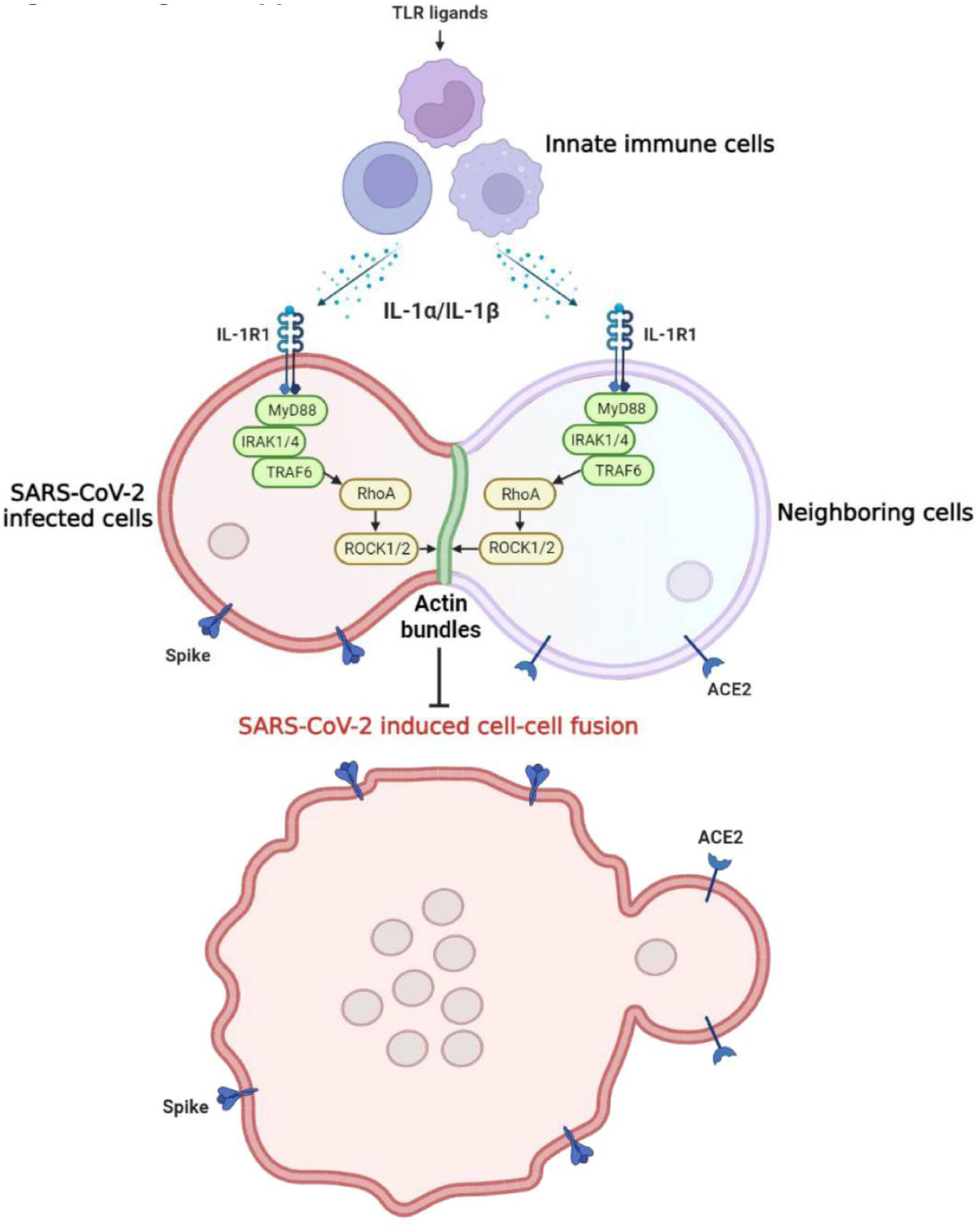
Graphical abstract. Host factors secreted from innate immune cells upon TLR ligands stimulation, including interleukin-1β (IL-1β) and IL-1α, act on both SARS-CoV-2 infected cells expressing spike protein and neighboring cells expressing ACE2 receptor via IL-1R1-MyD88-IRAK-TRAF6 signaling pathway, which leads to strong enrichment of activated RhoA at cell-cell junction, resulting in the formation of ROCK-mediated actin bundle to prevent SARS-CoV-2 induced cell-cell fusion and further viral transmission through syncytia formation.

**Table S1.**
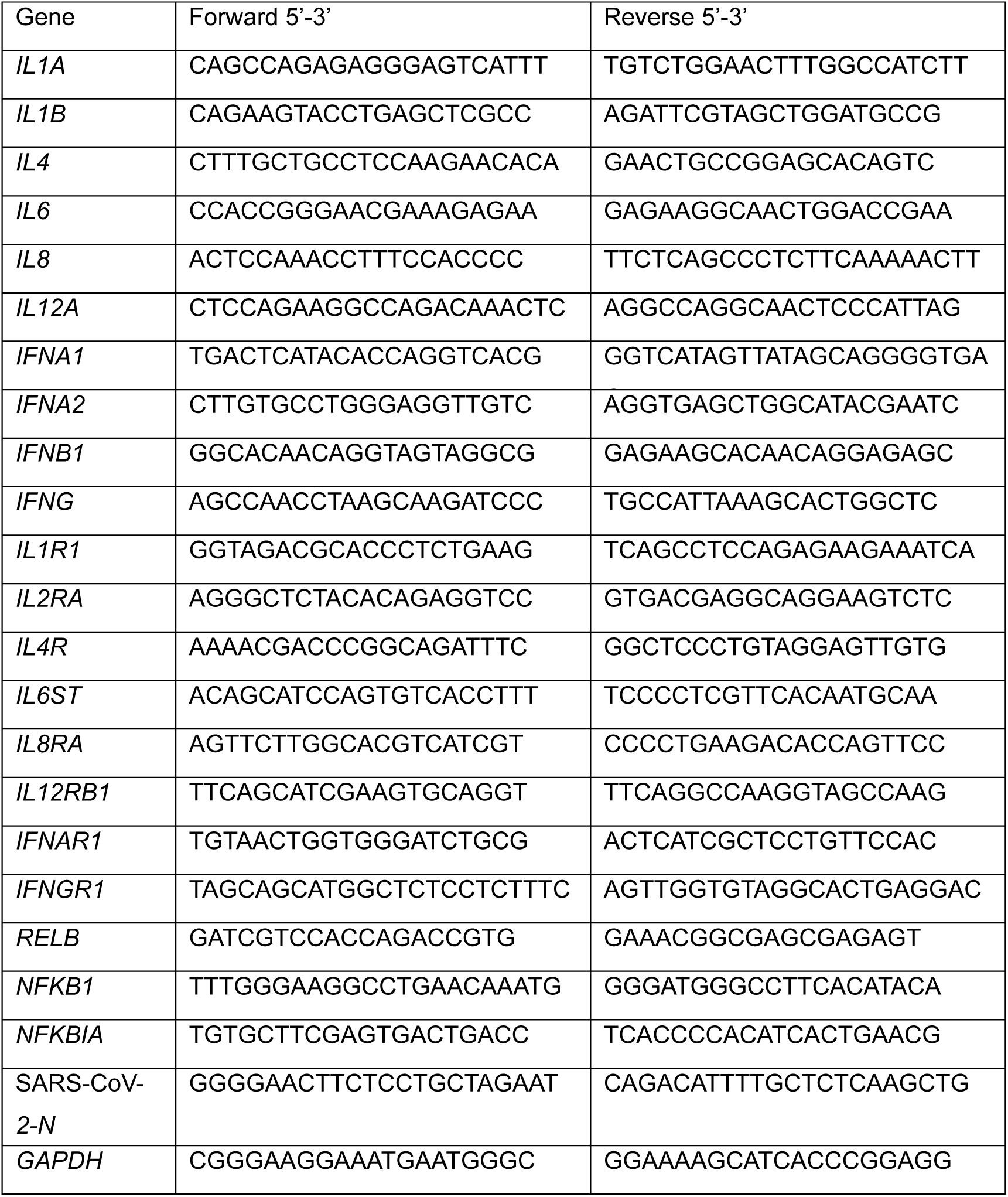
Primer sequences for RT-qPCR.

**Table S2.**
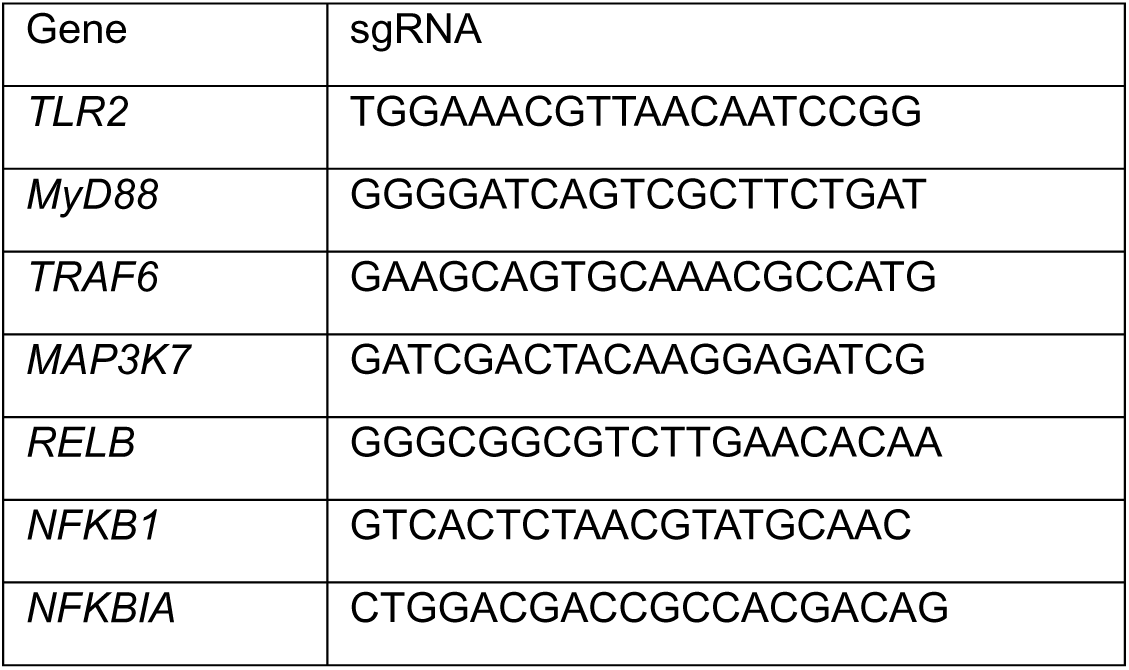
Primer sequences for sgRNA.

